# A standard knockout procedure alters expression of adjacent loci at the translational level

**DOI:** 10.1101/2021.08.21.457210

**Authors:** Artyom A. Egorov, Alexander I. Alexandrov, Valeriy N. Urakov, Desislava S. Makeeva, Roman O. Edakin, Artem S. Kushchenko, Vadim N. Gladyshev, Ivan V. Kulakovskiy, Sergey E. Dmitriev

## Abstract

The *S. cerevisiae* gene deletion collection is widely used for functional gene annotation and genetic interaction analyses. However, the standard G418-resistance cassette used to produce knockout mutants delivers strong regulatory elements into the target genetic loci. To date, its side effects on the expression of neighboring genes have never been systematically assessed. Here, using ribosome profiling data, RT-qPCR, and reporter expression, we investigated perturbations induced by the KanMX module. Our analysis revealed significant alterations in the transcription efficiency of neighboring genes and, more importantly, severe impairment of their mRNA translation, leading to changes in protein abundance. In the “head-to-head” orientation of the neighbor and the deleted gene, knockout often led to a shift of the transcription start site of the neighboring gene, introducing new uAUG codon(s) into the expanded 5’ untranslated region (5’ UTR). In the “tail-to-tail” arrangement, knockout led to activation of alternative polyadenylation signals in the neighboring gene, thus altering its 3’ UTR. These events may explain the so-called neighboring gene effect (NGE), i.e. false genetic interactions of the deleted genes. We estimate that in as much as ∼1/5 of knockout strains the expression of neighboring genes may be substantially (>2-fold) deregulated at the level of translation.

## INTRODUCTION

The haploid genome of the budding yeast *Saccharomyces cerevisiae* contains approximately 6,000 genes (1). Almost two decades ago, the complete gene knockout library was produced by the Saccharomyces Genome Deletion Project (SGDP) (2,3) with the goal of assigning a function to each ORF through phenotypic analysis of the mutants. For this, a PCR-based gene deletion strategy was used (4) to generate a complete start-to-stop-codon deletion of every ORF in the yeast genome. The ORF was replaced with a KanMX4 cassette (5) harboring the G418 resistance gene under control of the strong eEF1A promoter. The eEF1A terminator, adjacent vector-derived sequences, as well as additional artificial barcodes, were inserted along with the gene (Figure 1A).

**Figure 1.**
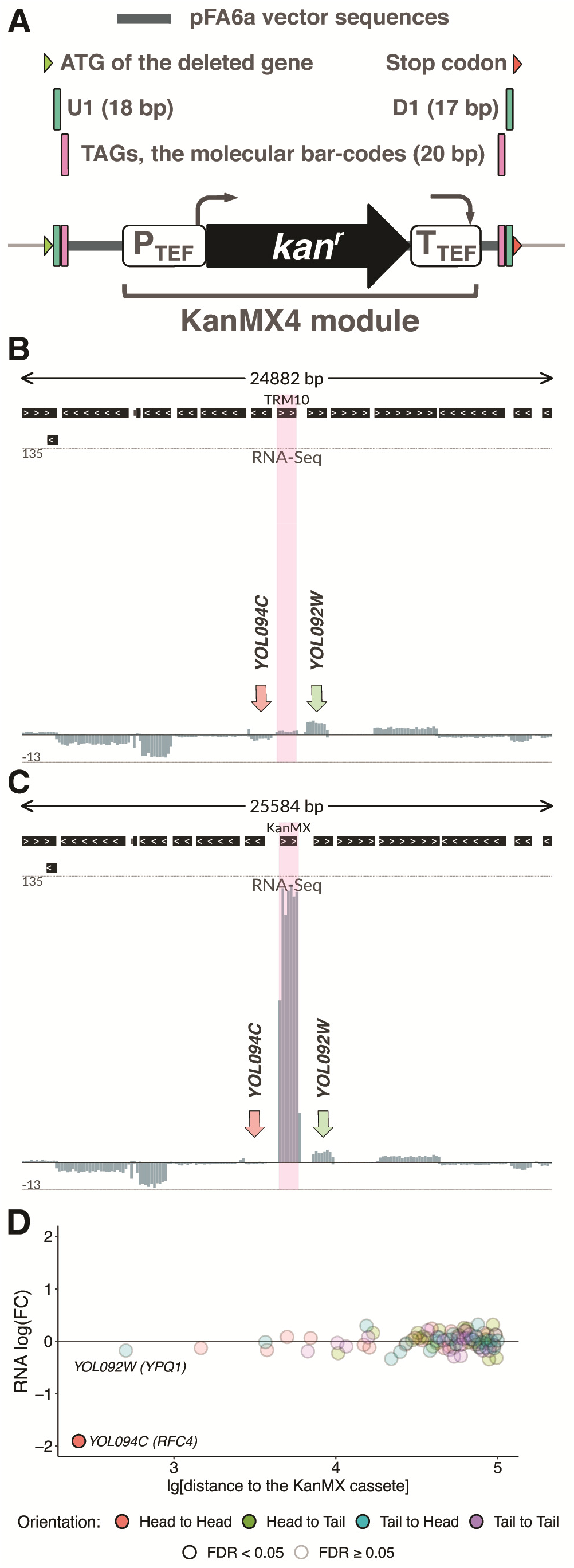
KanMX cassette used to generate gene deletions is excessively expressed within the mutated locus. (A) A schematic representation of the KanMX cassette replacing the knockout target gene. The cassette harbors the G418 resistance gene under the control of the strong *A*.*gossypii* eEF1A promoter and eEF1A terminator, as well as an adjacent vector-derived sequences and barcodes. (B,C) Excessive *kan* gene expression in the mutant strain, as compared to other genes in the vicinity. RNA-Seq data for the *TRM10* locus (including 15 genes in the vicinity) in *wt* (B) and *trm10*Δ (C) strains are shown as normalized total read coverage (positive and negative values correspond to the direct and reverse complementary DNA strands, respectively). The pink highlight indicates the *TRM10* CDS (*wt* strain) and the KanMX module (*trm10*Δ strain). RNA-Seq coverage of the neighboring genes, *RFC4* (*YOL094C*) and *YPQ1* (*YOL092W*), is indicated by arrows. (D) Differential expression of genes within the surrounding chromosome region (±100 kb from the KanMX module) at the transcriptional level, *trm10*Δ compared to *wt*. FDR: false discovery rate.

The creation of the SGDP yeast deletion collection revolutionized the field of eukaryotic molecular genetics. Since then, a huge amount of information was generated using the knockout strains. In particular, many genes have been functionally characterized according to the analysis of phenotype and genetic interactions of the corresponding deletion strains (6-8).

However, the target genetic loci were substantially modified during the procedure of knockout strain production. As *S. cerevisiae* has a compact genome with very short intergenic regions, the introduction of a highly expressed gene module could, in principle, significantly alter transcription within the loci, affecting the expression of neighboring genes. There were a few studies of the neighboring gene effects (NGE) in genetic interactions (9-11) but the mechanisms of such effects have never been analyzed.

Here, using RNA-Seq and ribosome profiling (Ribo-Seq) data available for a number of knockout strains, we revealed drastic changes of gene expression at a number of loci in which the target gene and its neighbor are in close proximity. Among the observed side-effects there were major alterations in transcription start sites (TSS) and polyadenylation site (PAS) usage of neighboring genes. These changes led to dramatic deregulation of the neighboring gene expression mainly at the level of mRNA translation that was further confirmed by analysis of protein abundance. Based on several dozen cases analyzed in this study, we roughly estimate that in as much as 10-20% of knockout strains in the yeast deletion library, the presence of the KanMX cassette substantially (>2-4-fold) deregulates expression of neighboring genes at the translational level.

## MATERIALS AND METHODS

### Data sources

Ribosome profiling (Ribo-Seq) and RNA-Seq data for knockout and wild-type *S. cerevisiae* strains (12,13) were downloaded from GEO [GSE100626, GSE122039]. Genome reference annotation and sequence for the BY4741 wt strain (version R64-1-1) was downloaded from Ensembl (14). Coordinates of alternative polyadenylation sites were taken from (15).

### Data processing

Knockout strain genome reference annotations and sequences were obtained with reform (https://github.com/gencorefacility/reform, (16)) through modification of the Ensembl (R64-1-1) annotation and sequences. SRA data were downloaded and transformed into fastq with SRA toolkit v2.9.4 (17). The adapters were trimmed with cutadapt v1.18 (18).

Read counting and mapping to the original and modified genome annotations were performed with STAR v2.7.0 (19). BedGraph profiles were produced from SAM data with samtools (20) and bedtools v2.27.1 (21). Coverage profiles were normalized using normalization factor and library size estimates from differential expression analysis (see below) separately for each bedGraph profile. Genomic signal tracks were visualized with svist4get (22).

### Ribo-Seq and RNA-Seq differential expression analysis

Raw gene counts processing and statistical analysis was performed in R using edgeR Bioconductor package (23). Genes not reaching 2 read count per million (CPM) in at least 3 RNA-Seq and 4 Ribo-Seq libraries were excluded from analysis. To detect diffentially expressed genes (for RNA-Seq, Ribo-Seq, and ribosome occupancy), we used the generalized linear model (glmQLFit, glmQLFTest of the edgeR package) with the strain as a categorical variable.

### Yeast strains

Yeast strains producing proteins tagged with C- and N-terminal Green Fluorescent Protein (GFP) were taken from systematic collections (24,25) and modified in order to delete genes adjacent to the gene of interest using the same strategy as in the original SGDP, except that the deletion cassette was obtained via 1 round of PCR with long primers. A list of the strains and primers used for the gene deletions is presented in Supplementary Tables 1 and 2, respectively. The gene deletion was verified by PCR of yeast genomic DNA using the primers listed in Supplementary Table 2.

### PCR-based quantification of normal and extended transcripts

Yeast strains with deletions of particular genes were grown in 100 ml of YPD to OD_600_ 0.5 at 30°C, collected and frozen in liquid nitrogen. Total RNA was extracted according to Collart et al. (26) using the hot acid phenol method. RNA samples were treated with RQ1 DNase (Promega). 1 mcg of total RNA was used for reverse transcription with MMLV-RT kit (Evrogen) according to the recommended protocol with oligo(dT)_15_ and random (dN)_10_ primers at a ratio of 1:3. Real-time PCR for detection of normal length and extended transcripts was performed with Eva Green I, kit №R-441 (Syntol), with primers listed in Supplementary Table 2. PCR protocol: 95°C – 5 min, [95°C – 15 sec, 64°C – 15 sec, 72°C – 30 sec, signal detection] x 43 cycles. Expression levels were calculated using the comparative Ct method.

### Flow cytometric analysis of the GFP-fusion protein abundance

Cytometric analysis was performed with 12 sets of three yeast strains (encoding a GFP-tagged protein (WT-GFP), the same strain containing a deletion of the adjacent gene (KO-GFP), and a strain harboring only the adjacent gene deletion (KO)), plus a wild-type strain as a control (WT). The strains were grown in complete synthetic medium (0.67% (w/v) Yeast Nitrogen Base (Difco), 2% (w/v) glucose, complete amino acid supplementation) overnight in 96-well flat bottom plates (Eppendorf), diluted into fresh medium with a dilution factor of 200, and grown for another 4 hours. After that they were analyzed using a Beckman Cytoflex S (488 nm laser, 525/40 bandpass emission filter) or a MACS Quant Analyzer, Miltenyi Biotech (488 nm laser, FITC emission filter), both equipped with an autosampler. Data on GFP fluorescence (FITC-A) were collected for ∼30 000 cells in 6 independent cultures of each sample, which were analyzed on two distinct days (3 in one day, 3 during another).

The raw data were extracted from *.fcs files with the FlowJo v.10 software. The FITC-A replicates of each strain were merged, the resulting distributions were compared between WT-GFP and KO-GFP strains with the Mann-Whitney *U* test. Additionally, we estimated the difference in medians Δmed_KO and Δmed_WT (by separately comparing *wt* and knockout fluorescence data against the matched autofluorescence data). Next, we computed the Δmed_KO/Δmed_WT ratio as the effect size estimate. To visualize distribution with the kernel density estimates we used the ggplot2 R package.

## RESULTS

### KanMX cassette expression is excessive in comparison to neighboring genes

Ribosome profiling combined with RNA-Seq provides expression data at the transcriptional and translational levels on a transcriptome-wide scale (27). Particular ribosome profiling studies were performed on yeast deletion strains. For our analysis we took the unique dataset produced by Chou et al. (12), where Ribo-Seq and RNA-Seq were performed for 57 yeast knockout strains, and our own ribosome profiling data (13). Both datasets were obtained from BY4741-based *S. cerevisiae* strains using the classical ribosome profiling protocol with cycloheximide (27). Although this drug may affect ribosome footprint distribution in the beginning of CDS (28) and at particular codons (29), it is nevertheless applicable for the analysis of overall ribosome coverage and differential expression (28).

First, we examined the transcriptional changes within a modified genomic locus, in which a target gene is replaced with the KanMX module (Figure 1A). To this end, we reconstructed the complete genome annotation for the knockout strains and mapped RNA-Seq reads to these genomes (*KO* datasets) or to the original genome of the *wt* BY4741 strain (*wt* dataset), respectively. Figures 1B-C illustrate the results of such mapping for *trm10*Δ and *wt* strains. We observed disproportionally high *kan* gene expression in the mutant strain, comparing to the transcription efficiency of other genes within the same locus (including at least a dozen upstream and downstream genes). Considering the whole transcriptome, *kan* is among the top 1% most highly expressed genes in the *trm10*Δ strain (Supplementary Figure S1A). The same effect is clearly exhibited in Ribo-Seq data (Supplementary Figures S1A-B), illustrating a high load of the *kan* transcript on the cell translational machinery.

The excessive transcription could skew the overall activity of the entire locus at the epigenetic level. Analysis of differential expression of the genes adjacent to *TRM10/YOL093W* showed that mRNA abundance of *RFC4/YOL094C* was reduced almost 4-fold, while the level of other mRNAs encoded in this locus did not experience observable changes (Figure 1D, Supplementary Figure S1B).

Similar analysis of RNA-Seq data for other 57 strains showed pervasive changes in transcription levels of neighboring genes (Supplementary Figure S2). Such alterations were unexpected and could potentially affect the abundance of encoded proteins and phenotype of the mutant strains.

### KanMX cassette shifts TSSs of the neighboring genes resulting in new uORFs and translational silencing

To further characterize unintended changes within mutated loci, we examined particular cases of the immediately adjacent genes in more detail. In the case of the *deg1Δ* strain, we observed a ∼3-fold decrease of the neighboring *SPB4* gene expression at the translational level (Figure 2A, cf. Ribo-Seq tracks of *wt* and *deg1Δ* strains, see also quantifications of Ribo-Seq and ribosome occupancy (RO) in Figure 2B; hereafter we use RO as Ribo-Seq coverage of a CDS normalized to its RNA-Seq coverage that serves as a proxy for translation efficiency of the transcript). The two genes were located in the 5’-to-5’ orientation (hereafter called “head-to-head”), with only 196 bp between their ATGs. Although overall *SPB4* mRNA level changed only slightly (Figure 2A, cf. RNA-Seq tracks, quantification is shown in Figure 2B), we observed a clear shift of the *SPB4* TSS toward the KanMX cassette in the case of the *deg1Δ* strain. Due to the shift, an upstream AUG codon was included in the new 5’ UTR, thus forming an inhibitory uORF (Figure 2C). This inhibited translation of the *SPB4* main coding region (Figure 2B, right panel), as uORFs usually dramatically decrease mRNA translation in yeast and are relatively rare in *S. cerevisiae* mRNAs (30-32). In the case of the altered *SPB4* transcript, the acquired uAUG was in the optimal nucleotide context (AggAUGA) and thus should be effectively recognized by the ribosome (33). Indeed, we observed ribosome coverage of a region corresponding to the uORF (Figure 2A). In this regard, the slightly decreased abundance of the *SPB4* mRNA (Figure 2A and B, RNA-Seq tracks) could be related to its reduced stability (34), as uORF-containing transcripts are subject to nonsense-mediated decay in yeast (35,36).

**Figure 2.**
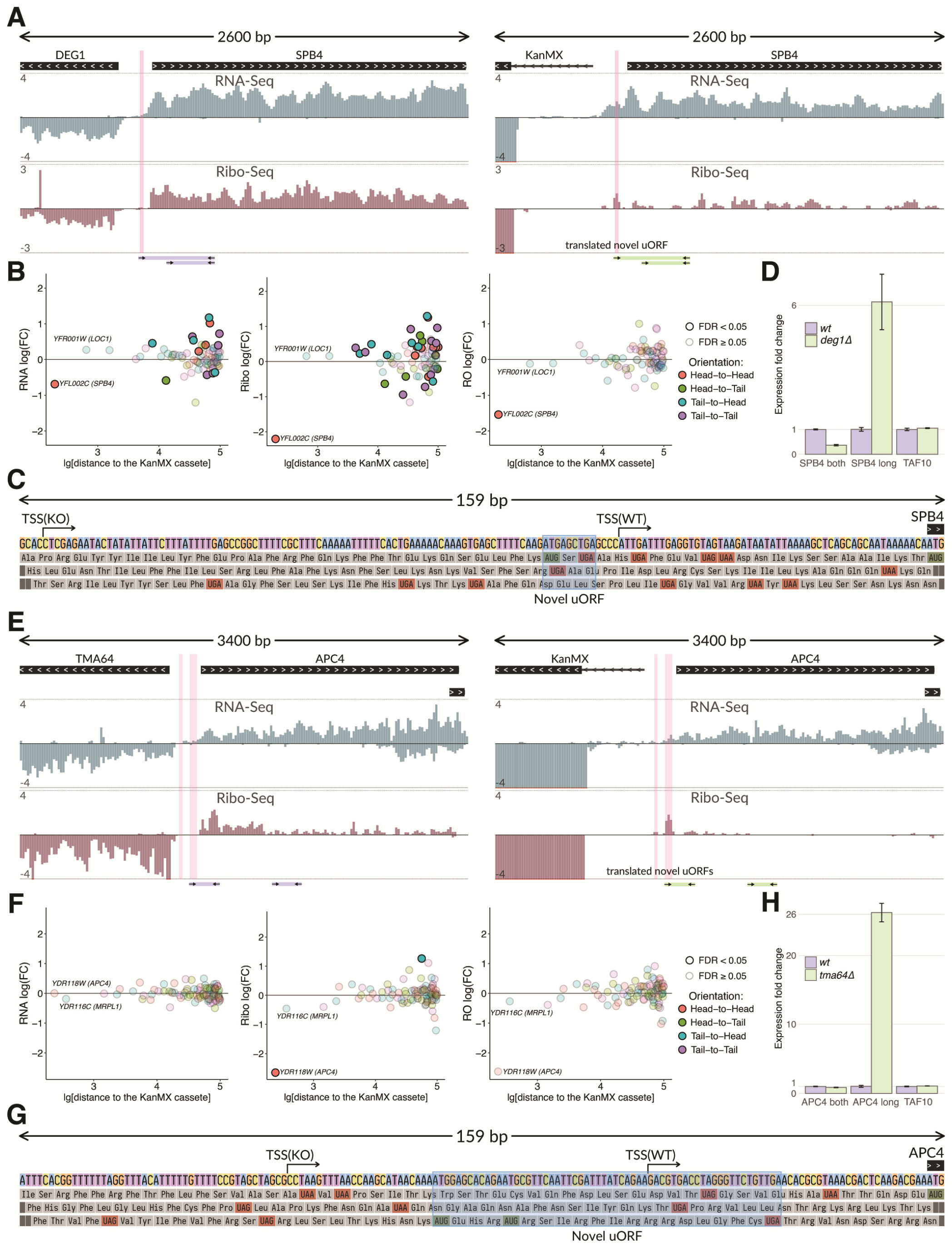
KanMX cassette shifts transcriptional start sites of neighboring genes generating novel uORFs and inducing translational silencing. (A) RNA-Seq and Ribo-Seq normalized coverage of the *DEG1-SPB4* locus in the *wt* (left) and *deg1*Δ (right) strains. The tracks show normalized coverage (positive and negative values correspond to the direct and reverse complementary DNA strands, respectively). Translation of the novel uORF (highlighted in pink) causes a decrease of the *SPB4* ribosome occupancy. Primer pairs and the corresponding PCR products used for validation of the TSS shift in (D) are schematically shown below the tracks. (B) Differential expression of genes within the surrounding chromosome region (±100 kb from the KanMX module) at the level of transcription (RNA-Seq, left panel), translation (Ribo-Seq, middle panel), and translation efficiency (RO, right panel), *deg1*Δ compared to *wt*. RO (ribosome occupancy) is the Ribo-Seq coverage of a CDS normalized to the RNA-Seq coverage. (C) Nucleotide sequence of the *SPB4* upstream region and its three-frame translation. The shift of the *SPB4* TSS (indicated by arrows) in the *deg1*Δ strain creates a novel uORF (shadowed). TSSs in both strains are determined as 5’ ends of the longest transcripts present in the RNA-Seq data. (D) RT-qPCR confirms 5’ UTR elongation of the *SPB4* mRNA in the *deg1*Δ strain. Primer pairs specific for the extended *SPB4* transcript (“SPB long”), for both shorter and longer isoforms (“SPB4 both”) or for *TAF10* mRNA (control) are used, as shown in (A). Product abundances are normalized to those in the *wt* strain. (E-H) The same as in (A-D) for the *TMA64-APC4* genetic locus. Shift of the *APC4* TSS in the *tma64*Δ strain creates a novel uORF, which decreases the *APC4* translation efficiency.

The TSS shift of SPB4 gene in the *deg1Δ* strain was confirmed experimentally by RT-qPCR with two primer pairs (Supplementary Table 2, schematically shown by arrows in Figure 2A). A forward primer from the first pair corresponded to the extended part of the presumptive extended *SPB4* transcript, while that from the other pair annealed to both the extended and original (normal length) mRNAs. In the strain with deleted *DEG1* we observed a substantially elevated amount of the extended variant, while the total abundance of both *SPB4* transcripts was reduced (Figure 2D).

An even more dramatic effect was observed in the case of *TMA64* gene knockout (13), which led to the deregulation of the neighboring essential gene *APC4*, located in the “head-to-head” orientation (Figure 2E). In *tma64Δ* we observed a ∼8-fold decrease of the *APC4* CDS expression at the translational level (Figure 2F). The region upstream of the original *APC4* TSS contained two ATG codons (Figure 2G). Replacing *TMA64* with the KanMX module shifted the *APC4* TSS upstream, thus forming two uORFs clearly covered by ribosome footprints in the Ribo-Seq track in Figure 2E. The RT-qPCR analysis confirmed that the abundance of the extended *APC4* transcript was dramatically increased in the *tma64Δ* strain (Figure 2H). *APC4* encodes a subunit of the Anaphase-Promoting Complex/Cyclosome (APC/C), involved in the control of cell cycle (37), while the *TMA64* gene product is a non-essential translation factor involved in ribosome recycling and reinitiation (38). Curiously, previous analyses of *tma64Δ* knockout strains revealed strong negative genetic interactions of *TMA64* with genes involved in APC/C-mediated anaphase checkpoint (39,40), as well as cell cycle abnormalities (41). Indeed, 7 out of 10 *TMA64* genetic interactions listed in SGD (https://www.yeastgenome.org, (42)) and documented in at least 2 data sources, encode either APC/C subunits (APC5, CDC20, CDC23, CDC27) or other components of the mitotic checkpoint (Supplementary Figure S3). Our data suggest that these interactions are false positives caused by the defect of the APC4 mRNA translation in the *tma64Δ* knockout strain that we found in our analysis.

### KanMX cassette alters 3’ end processing of the neighboring gene transcripts and deregulates their translation

The above cases exemplified the “head-to-head” arrangement of a target gene with its affected neighbor. However, the KanMX module could also disturb proper termination and correct formation of the 3’ end of the transcript in the case of the 3’-to-3’ (“tail-to-tail”) arrangement. To investigate this, we analyzed a number of such cases from the same study (12), and indeed found shifts of cleavage and polyadenylation sites (PASs).

Particularly, the PAS shift happens in the *TRM12* knockout, with the dramatic alteration in PAS position of the neighboring *GLO1* gene (Figure 3A). In this particular case, the alteration was probably unavoidable, as the original *GLO1* transcript extends into the *TRM12* coding region, where its PAS is located normally. In the *trm12Δ* strain, however, this region is replaced by the KanMX cassette. Mapping of the RNA-Seq reads onto the *trm12Δ* genome revealed the formation of an alternative 3’ terminus, which probably was the result of an activation of the cryptic premature PAS located in this locus (15) (Figures 3A and 3C). The resulting 3’ UTR is ∼2-fold shorter than the original one. Surprisingly, the dramatic changes in the *GLO1* 3’ UTR length did not lead to prominent perturbations in either abundance of the transcript or its RO (Figure 3B), at least under conditions tested. However, in other cases such alterations could affect gene expression, as 3’ UTRs play important roles in translation regulation and control of mRNA stability (43,44). Indeed, in the case of another “tail-to-tail” arranged gene pair, *SET3* and *SAP190*, we observed substantial (∼3-fold) reduction in RO of the *SET3* coding region in the strain where *SAP190* was replaced with the KanMX cassette (Figures 3D and 3E), as expected from the dramatic shortening of the *SET3* 3’ UTR (Figures 3D and 3F).

**Figure 3.**
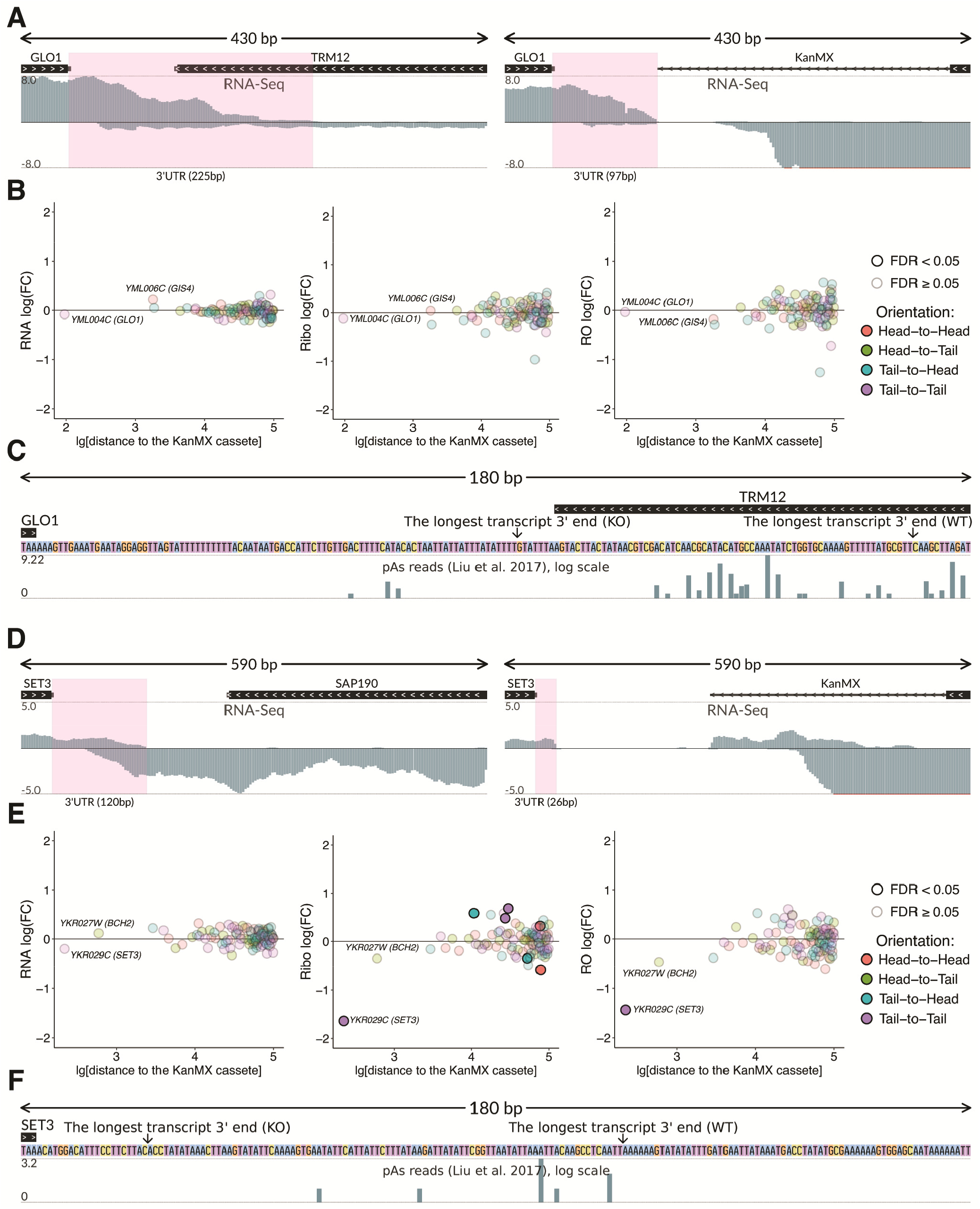
KanMX cassette deregulates expression of the neighboring gene transcripts at the translational level by altering 3’ end processing. (A) Normalized RNA-Seq coverage of the *GLO1-TRM12* locus in *wt* (left) and *trm12*Δ (right) strains. The *GLO1* 3’ UTR (highlighted in pink) is shorter in the mutant strain. (B) Differential expression of genes within the surrounding chromosome region (±100 kb from the KanMX module) at the level of transcription (RNA-Seq, left panel), translation (Ribo-Seq, middle panel), and translation efficiency (RO, right panel), *trm12*Δ compared to *wt*. (C) Nucleotide sequence of the *GLO1* downstream region. The putative major PASs of the *GLO1* gene in both strains are indicated by arrows. Putative PASs from (15) are shown in additional track, Y-axis in log-scale. (D-F) The same for the *SET3-SAP190* genetic locus, in (D) *SET3* 3’ UTR is highlighted in pink.

### Transcriptional and translational deregulation of neighboring genes caused by TSS shifts and alternative PAS activation are pervasive in yeast knockout strains

To systematize TSS shifting and alternative PAS activation, we estimated alterations of 5’ UTR and 3’ UTR lengths of neighboring genes observed in the analyzed datasets, depending on the mutual orientation of the knockout target and the nearest neighboring gene (Figures 4A). The analysis showed that “head-to-head” orientation often leads to TSS shifting: 5’ UTRs were extended by the TSS shifts in 11 of the 12 cases, when the distance between the ORFs of the neighboring genes was less than 1 kb. In the case of the “tail-to-tail” orientation, we found a trend towards 3’ UTRs shortening as a result of alternative PAS activation, but this effect was less exhibited in comparison with the TSS shift. In the case of “head-to-tail” and “tail-to-head” orientations, no significant alterations were observed, although particular changes are harder to track in such cases, as RNA sequences of the neighboring and knockout genes are located on the same DNA strand in close proximity.

**Figure 4.**
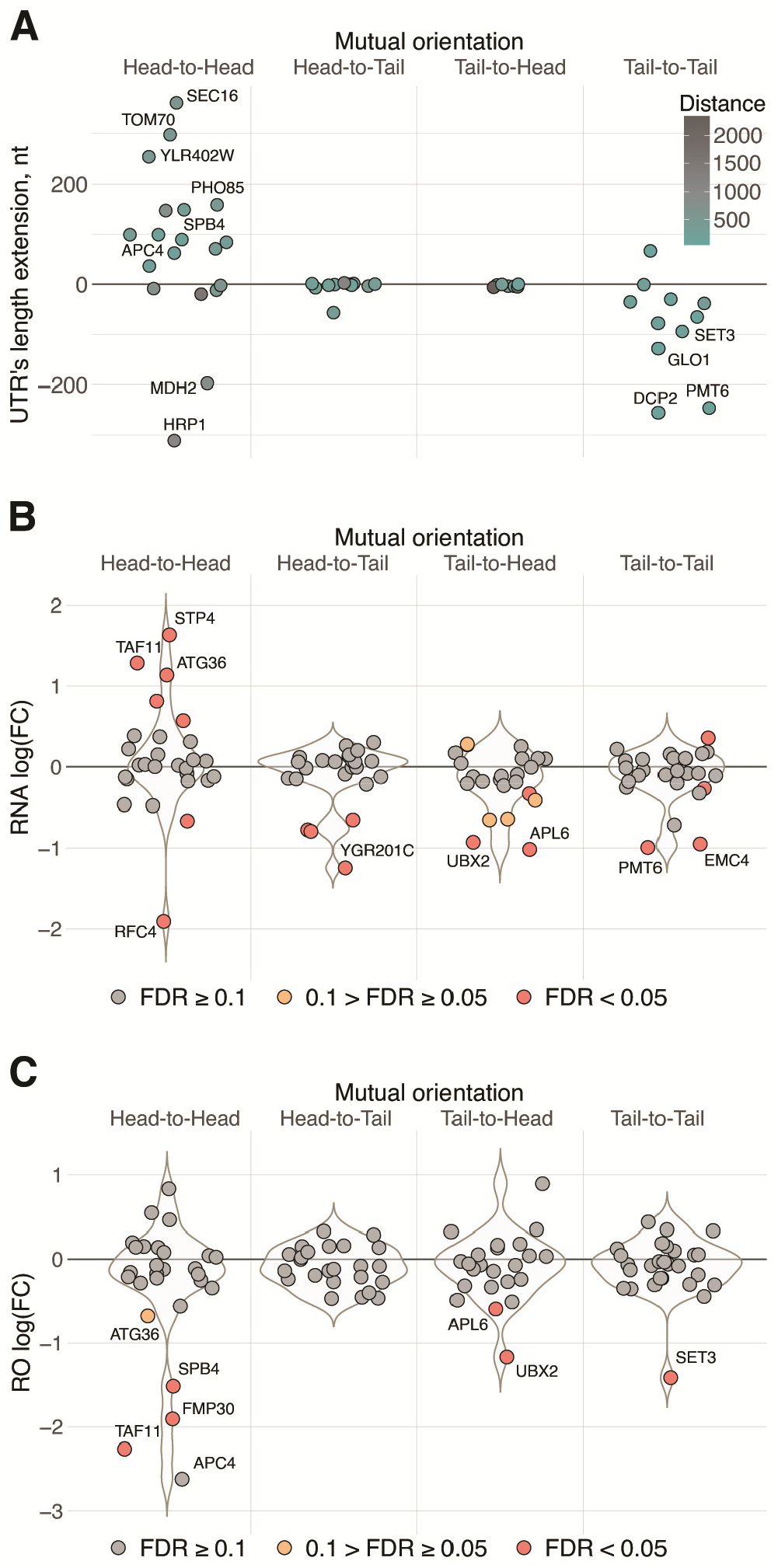
KanMX cassette alters 5’ and 3’ UTRs of neighboring genes, affecting transcript abundances and translation efficiencies. (A) Alterations of 5’ UTR and 3’ UTR lengths of neighboring genes induced by the KanMX module depending on their orientation to the cassette. (B) Differential expression of the neighboring genes in the 58 knockout strains analyzed in this study, at the level of transcription (RNA-Seq), as compared to *wt*. (C) Changes in the translation efficiency (RO) of mRNAs encoded by the neighboring genes in knockout strains compared to *wt*.

We then calculated expression changes of the closest neighboring genes in all datasets at both transcriptional and translational levels, and grouped the results with respect to the neighboring gene orientation to the deleted genes. Analysis of the RNA-Seq coverage (Figures 4B) showed strong transcriptional deregulation of many “head-to-head”-oriented neighboring genes, with either decreased or increased expression; for other mutual orientations, there was a tendency to observe the reduced mRNA level. Similar analysis of RO, which reflects translation efficiency (Figure 4C), revealed a prominent effect of translational downregulation for a substantial proportion of neighboring genes, leading to more than 2-fold drop (FDR<0.05) in at least 6 cases (Figure 4C), four of which can be explained by the above-described effect from novel uORFs (see Supplementary Figure S2).

### As much as 1/5 of knockout strains may have substantially (>2-fold) deregulated expression of neighboring genes that can result in changed protein abundance

Since ribosome coverage of a transcript can be used as a proxy for the gene product abundance, we analyzed Ribo-Seq data to calculate a proportion of cases in the analyzed datasets, where the protein synthesis from neighboring gene transcripts was deregulated (Figure 5A). Among the 58 analyzed strains, ribosome footprint coverage of neighboring gene transcripts was altered more than 2-fold (FDR<0.05) in 11 (19%), >3-fold in 7 (12%), and >4-fold in 5 cases (8%). Weaker effects (from 1.5- to 2-fold) were observed in 20 (34%) of the analyzed strains. This gives a rough estimate of the number of affected strains across the complete yeast deletion library.

**Figure 5.**
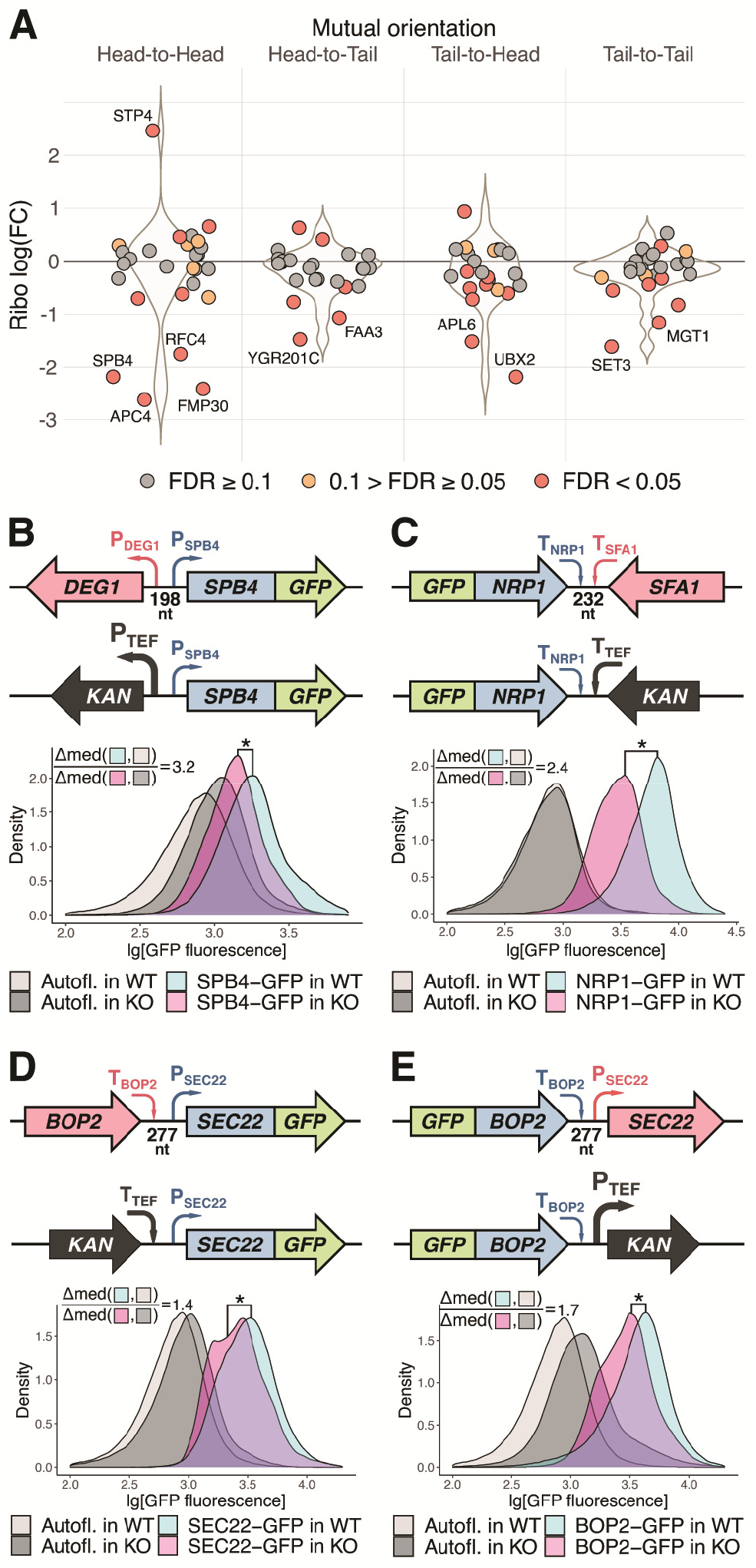
A substantial portion of analyzed knockout strains have deregulated expression of neighboring genes that can result in changes of the protein abundances. (A) Differential expression of the neighboring genes in the 58 knockout strains analyzed in this study at the level of translation (Ribo-Seq) depending on neighboring gene orientation to the *kan* gene (knockout strains compared to *wt*). (B-E) Gene replacement with the KanMX module induces changes in the abundance of the GFP-tagged proteins encoded by neighboring genes. Four cases with different mutual gene orientations (schematically shown in upper subpanels) were analyzed. For each case, 4 yeast strains were assayed by flow cytometry: 2 strains with the GFP-tagged neighboring gene (as shown in the schemes) and 2 corresponding strains without the GFP-tag (to take into account possible autofluorescence changes between the *KO* strains and *wt*). The density plots show distribution of the GFP fluorescence (log-scale, combined results from at least three independent experiments). Effect size estimates (Δmed ratios) are shown on the plots, see Materials and Methods for details. Combined results from at least three independent experiments are shown, *one-sided U-test p-value <10^−20^.

However, protein abundance is determined not only by synthesis but also degradation. Thus, it was important to experimentally verify the changes in the abundance of proteins encoded by the genes neighboring the KanMX module in knockout strains. We selected a number of strains expressing GFP-tagged proteins from the GFP-tagged systematic collections (24,25) and replaced the adjacent genes with the standard KanMX cassette. *Wt* and deletion strains having no GFP tag were used for autofluorescence background correction.

We assayed 5 strains with “head-to-head” orientation of genes, 4 strains with “tail-to-tail” orientation, and 3 with “head-to-tail” or “tail-to-head” orientation, in which we could reliably detect GFP fluorescence using flow cytometry. The set included GFP-tagged SPB4 protein corresponding to the *DEG1*-*SPB4* gene analyzed earlier (Figures 2A and 5B) and two complementing pairs of genes: *BOP2―SEC22-GFP* / *GFP-BOP2―SEC22*, and *RRD1―IMP2’-GFP* / IMP2’*―RRD1-GFP*). Importantly, half of the deleted genes (*IMP2’, IRC6, RRD1, SEC22, SFA1, VIP1*) were unrelated to translation, thus avoiding a bias of the Ribo-Seq analysis where most of the target genes were encoding tRNA modification enzymes (12).

A statistically significant decrease in the level of GFP fluorescence was observed in 6 out of the tested 12 cases (Figures 5B-C and Supplementary Figure S4), including the GFP-tagged SPB4. For 4 proteins we were unable to detect any significant changes of the protein level, while in one case the reporter level was increased (Supplementary Figure S4D). To sum up, deregulation of the expression of adjacent genes can affect their protein levels in the cell and, consequently, influence the phenotype of the knockout strains.

## DISCUSSION

The SGDP yeast deletion library is a powerful and indispensable instrument to study eukaryotic gene function (2,6). However, due to the compact nature of the yeast genome, replacement of every individual gene with the highly expressed KanMX cassette can potentially perturb the expression of the neighboring genes. Importantly, the undesirable effects of the cassette could extend beyond the transcriptional level, affecting also translation of mRNAs synthesized from neighboring genes. Such changes may cause inappropriate attribution of the side effects to the absence of the target gene, while in reality they are actually caused by perturbation of neighboring gene expression. This phenomenon is known as the neighboring gene effect, NGE (10,11). NGE should be taken into account during high-throughput studies of functional gene annotation and genetic interaction analyses. A few systematic studies devoted to this important issue revealed that NGE may be involved in as much as 7-15% of the identified gene-gene interactions (9-11). However, no mechanistic explanations of these effects have been proposed.

In this study, we took advantage of ribosome profiling (45,46) to investigate the molecular mechanisms behind NGE. Although this powerful method was originally developed for *S. cerevisiae* (27,47), to date only a limited number of Ribo-Seq studies have been performed in knockout yeast strains. We used one of the most comprehensive datasets containing ribosome profiling and RNA-Seq data for 57 mutant strains (12), as well as our own data (13), to characterize the transcriptional and translational changes that occur in genomic regions adjacent to the introduced KanMX cassette. We documented several types of changes, including (i) transcriptional repression of the neighboring genes (Figures 1 and 4B); (ii) shifts of TSSs toward the cassette, causing the appearance of uAUG(s) in the extended 5’ UTRs, reduction of translation efficiency and likely mRNA stability (Figures 2 and 4A); and (iii) triggering alternative pre-mRNA cleavage and polyadenylation events, in some cases also leading to deregulation of mRNA expression at the translational level (Figures 3 and 4A). Finally, using GFP reporters, we showed that such perturbations can result in changes of protein abundance (Figures 5B and 5C).

Although our study was restrained by the limited amount of published yeast Ribo-Seq datasets, the available examples clearly show that replacement of a gene by the KanMX cassette often significantly affects expression of the neighboring genes via both transcriptional and translational mechanisms.

One could argue that all datasets that we initially analyzed were obtained for strains with mutations of translation machinery components. Indeed, ribosome profiling is usually applied to mutants or conditions where translation is expected to be altered. However, in a large portion of these strains no phenotype associated with compromised overall translation was observed (12). We also showed that only the adjacent genes were typically exhibiting differential expression at the level of transcription or translation (see Supplementary Figure S2). Moreover, our GFP-based validation assay was also applied to genes not involved in translation. Thus, the observed phenomenon is likely general and not limited to translation-related genes. Extrapolating our results, we estimate that changes in the levels of transcription and translation of neighboring genes of 2-fold and higher may be observed in up to one fifth of knockout strains in the yeast deletion library.

The revealed TSS shifts (in the case of “head-to-head” gene arrangement) and alternative PAS activation (in “tail-to-tail” orientation of the genes), leading to mRNA translation decline, were the most striking observations of our study (Figure 4A, overviewed in Figure 6). The effects of the TSS shifts and PAS changes in knockout strains may be even more important under different growth conditions, cell stress or sporulation, when numerous alternative TSSs and PASs are activated (48,49).

**Figure 6.**
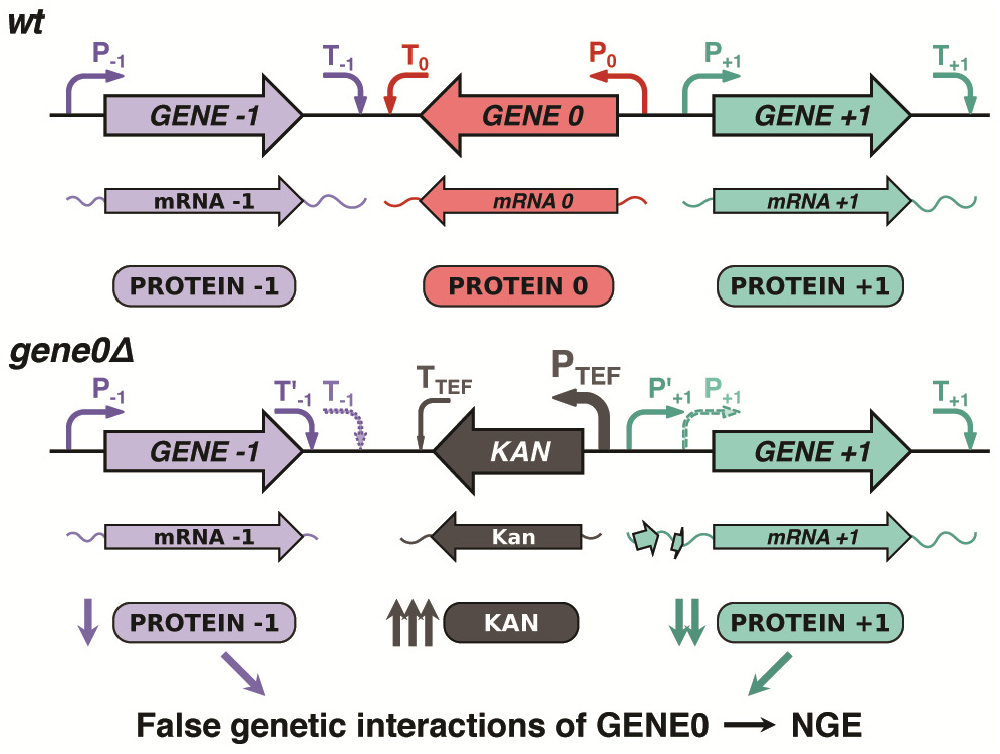
A model explaining NGE in knockout strains arising from perturbation of neighboring gene expression at the translational level. As *GENE0* is replaced by the KanMX cassette, its strong regulatory elements (P-TEF and T-TEF) perturb transcription of the neighboring genes, altering PAS and TSS of *GENE-1* and *GENE+1*, respectively. This decreases their mRNA translation due to the GENE-1 3’ UTR shortening and uORF(s) appearance in the *GENE+1* 5’ UTR.

Taking into account the pervasiveness of such effects, we compiled a complete list of potentially affected genes (Supplementary Table 3) where one can find a qualitative estimation of an expected effect, which replacement by the KanMX module may induce on the nearest adjacent gene. The estimation is based on a set of empirical rules revealed in this study: mutual gene orientation, distance between the genes, and, in the case of “head-to-head” orientation, the presence of AUG codon in the spacer located near the TSS of the neighboring gene.

The identified patterns of expression changes of neighboring genes could explain the false genetic interactions of the deleted genes, so-called NGE phenomenon (Figure 6), both in the cases we examined here and in many other cases. For example, almost a half of the neighboring gene pairs with the distance between the genes less than 200 bp and with a genetic interaction profile similarity of PCC > 0.1 (according to the TheCellMap.org (9,40)) have at least one ATG codon within 50 bp upstream of a TSS of the neighboring gene, while the same is true for only a quarter of pairs with such arrangement of genes and with PCC < 0.01 (see Supplementary Table 3). This probably could explain the observed NGE in some of the former gene pairs, with clearer criteria to be formulated when more high-throughput data become available for systematic analysis.

All in all, we propose that in many cases NGE in knockout strains is caused by KanMX-induced defects at the translational level, as revealed in this study. This should be taken into account when using the yeast knockout collection for functional gene annotation and genetic interaction analysis, as well as when interpreting results based on the similar methods of targeted gene manipulation in any other model eukaryotic organism.

## CONFLICT OF INTEREST

The authors have no conflicts of interest to disclose.

### ABBREVIATIONS

SGDP: Saccharomyces Genome Deletion Project,
NGE: neighboring gene effect; 5’
UTR: 5’ untranslated region; 3’
UTR: 3’ untranslated region;
uAUG: upstream AUG codon;
ORF: upstream open reading frame;
TSS: transcriptional start site;
PAS: pre-mRNA cleavage and polyadenylation site;
GFP: green fluorescent protein;
RO: ribosome occupancy.

## SUPPLEMENTARY DATA

Supplementary Data.pdf contains Supplementary Tables 1, 2, and Supplementary Figures S1, S2, and S4.

Supplementary Figure 2.pdf

Supplementary Table 3.xlsx

## ACKNOWLEDGMENTS

We are grateful to Pavel Baranov (University College Cork, Ireland) for helpful discussion and the Moscow State University Development Program PNR5 for providing access to the MACS Quant Analyzer flow cytometer, as well as the Shared-Access Equipment Centre “Industrial Biotechnology” of Federal Research Center “Fundamentals of Biotechnology” of Russian Academy of Sciences for providing access to the Beckman Cytoflex S flow cytometer. We also thank Dr. Maya Schuldiner for sharing N-terminally GFP-tagged yeast strains. A.A.E., R.O.E., A.S.K., and S.E.D. are members of Interdisciplinary Scientific and Educational School of Moscow University ≪Molecular Technologies of the Living Systems and Synthetic Biology≫.

