## Supplementary Tables 1, 2; Supplementary Figures S1, S3, and S4 for "A standard knockout procedure alters expression of adjacent loci at the translational level"

### SUPPLEMENTARY DATA

**Supplementary Table 1**. Yeast strains used in this study.

| Strain name | Strain background | Genotype | Origin |
|---|---|---|---|
| *wt* | BY4741 | *MATa his3Δ1 leu2Δ0 met15Δ0 ura3Δ0* | Yeast deletion collection |
| *tma64Δ* | BY4741 | *MATa his3Δ1 leu2Δ0 met15Δ0 ura3Δ0 tma64::KanMX* | Yeast deletion collection |
| *spb4-gfp* | BY4741 | *MATa his3Δ1 leu2Δ0 met15Δ0 ura3Δ0 SPB4-GFP::HIS3* | Huh et al. 2003 |
| *spb4-gfp deg1Δ* | BY4741 | *MATa his3Δ1 leu2Δ0 met15Δ0 ura3Δ0 SPB4-GFP::HIS3 deg1::KanMX* | This study |
| *deg1Δ* | BY4741 | *MATa his3Δ1 leu2Δ0 met15Δ0 ura3Δ0 deg1::KanMX* | Yeast deletion collection |
| *sec22-gfp* | BY4741 | *MATa his3Δ1 leu2Δ0 met15Δ0 ura3Δ0 SEC22-GFP::HIS3* | Huh et al. 2003 |
| *sec22-gfp bop2Δ* | BY4741 | *MATa his3Δ1 leu2Δ0 met15Δ0 ura3Δ0 SEC22-GFP::HIS3 bop2::KanMX* | This study |
| *bop2Δ* | BY4741 | *MATa his3Δ1 leu2Δ0 met15Δ0 ura3Δ0 bop2::KanMX* | Yeast deletion collection |
| *bop2-gfp* | BY4741 | *MATa his3Δ1 leu2Δ0 met15Δ0 ura3Δ0 BOP2-GFP::HIS3* | Huh et al. 2003 |
| *bop2-gfp sec22Δ* | BY4741 | *MATa his3Δ1 leu2Δ0 met15Δ0 ura3Δ0 BOP2-GFP::HIS3 sec22::KanMX* | This study |
| *sec22Δ* | BY4741 | *MATa his3Δ1 leu2Δ0 met15Δ0 ura3Δ0 sec22::KanMX* | Yeast deletion collection |
| *nrp1-gfp* | BY4741 | *MATa his3Δ1 leu2Δ0 met15Δ0 ura3Δ0 NRP1-GFP::HIS3* | Huh et al. 2003 |
| *nrp1-gfp sfa1Δ* | BY4741 | *MATa his3Δ1 leu2Δ0 met15Δ0 ura3Δ0 NRP1-GFP::HIS3 sfa1::KanMX* | This study |
| *sfa1Δ* | BY4741 | *MATa his3Δ1 leu2Δ0 met15Δ0 ura3Δ0 sfa1::KanMX* | Yeast deletion collection |
| *imp2'-gfp* | BY4741 | *MATa his3Δ1 leu2Δ0 met15Δ0 ura3Δ0 IMP2'-GFP::HIS3* | Huh et al. 2003 |
| *imp2'-gfp rrd1Δ* | BY4741 | *MATa his3Δ1 leu2Δ0 met15Δ0 ura3Δ0 IMP2'-GFP::HIS3 rrd1::KanMX* | This study |
| *rrd1Δ* | BY4741 | *MATa his3Δ1 leu2Δ0 met15Δ0 ura3Δ0 rrd1::KanMX* | Yeast deletion collection |
| *rrd1-gfp* | BY4741 | *MATa his3Δ1 leu2Δ0 met15Δ0 ura3Δ0 RRD1-GFP::HIS3* | Huh et al. 2003 |
| *rrd1-gfp imp2'Δ* | BY4741 | *MATa his3Δ1 leu2Δ0 met15Δ0 ura3Δ0 RRD1-GFP::HIS3 imp2'::KanMX* | This study |
| *imp2'Δ* | BY4741 | *MATa his3Δ1 leu2Δ0 met15Δ0 ura3Δ0 imp2'::KanMX* | Yeast deletion collection |
| *utp21-gfp* | BY4741 | *MATa his3Δ1 leu2Δ0 met15Δ0 ura3Δ0 UTP21-GFP::HIS3* | Huh et al. 2003 |
| *utp21-gfp vip1Δ* | BY4741 | *MATa his3Δ1 leu2Δ0 met15Δ0 ura3Δ0 UTP21-GFP::HIS3 vip1::KanMX* | This study |
| *vip1Δ* | BY4741 | *MATa his3Δ1 leu2Δ0 met15Δ0 ura3Δ0 vip1::KanMX* | Yeast deletion collection |
| *atg20-gfp* | BY4741 | *MATa his3Δ1 leu2Δ0 met15Δ0 ura3Δ0 ATG20-GFP::HIS3* | Huh et al. 2003 |
| *atg20-gfp* | BY4741 | *MATa his3Δ1 leu2Δ0 met15Δ0 ura3Δ0 ATG20-GFP::HIS3* | This study |

| | | | |
|---|---|---|---|
| *trm3Δ* | | *trm3::KanMX* | |
| *trm3Δ* | BY4741 | *MATa his3Δ1 leu2Δ0 met15Δ0 ura3Δ0 trm3::KanMX* | Yeast deletion collection |
| *gfp-sec5* | BY4741 | *MATa his3Δ1 leu2Δ0 met15Δ0 ura3Δ0 pNOP1-GFP-SEC5::URA3* | Weill et al. 2018 |
| *gfp-sec5 trm82Δ* | BY4741 | *MATa his3Δ1 leu2Δ0 met15Δ0 ura3Δ0 pNOP1-GFP-SEC5::URA3 trm82::KanMX* | This study |
| *trm82Δ* | BY4741 | *MATa his3Δ1 leu2Δ0 met15Δ0 ura3Δ0 trm82::KanMX* | Yeast deletion collection |
| *gfp-keg1* | BY4741 | *MATa his3Δ1 leu2Δ0 met15Δ0 ura3Δ0 pNOP1-GFP-KEG1::URA3* | Weill et al. 2018 |
| *gfp-keg1 irc6Δ* | BY4741 | *MATa his3Δ1 leu2Δ0 met15Δ0 ura3Δ0 pNOP1-GFP-KEG1::URA3 irc6::KanMX* | This study |
| *irc6Δ* | BY4741 | *MATa his3Δ1 leu2Δ0 met15Δ0 ura3Δ0 irc6::KanMX* | Yeast deletion collection |
| *gfp-enp2* | BY4741 | *MATa his3Δ1 leu2Δ0 met15Δ0 ura3Δ0 pNOP1-GFP-ENP2::URA3* | Weill et al. 2018 |
| *gfp-enp2 ecl1Δ* | BY4741 | *MATa his3Δ1 leu2Δ0 met15Δ0 ura3Δ0 pNOP1-GFP-ENP2::URA3 ecl1::KanMX* | This study |
| *ecl1Δ* | BY4741 | *MATa his3Δ1 leu2Δ0 met15Δ0 ura3Δ0 ecl1::KanMX* | Yeast deletion collection |
| *ubx2-gfp* | BY4741 | *MATa his3Δ1 leu2Δ0 met15Δ0 ura3Δ0 UBX2-GFP::HIS3* | Huh et al. 2003 |
| *ubx2-gfp trm9Δ* | BY4741 | *MATa his3Δ1 leu2Δ0 met15Δ0 ura3Δ0 UBX2-GFP::HIS3 trm9::KanMX* | This study |
| *trm9Δ* | BY4741 | *MATa his3Δ1 leu2Δ0 met15Δ0 ura3Δ0 trm9::KanMX* | Yeast deletion collection |

**Supplementary Table 2**. Primers used in this study.

**I. Primers for yeast strain construction**

| GFP-tagged gene | Deleted gene | Primer sequence | Primer type (F – forward, R – reverse) |
|---|---|---|---|
| *ENP2* | *ECL1* | CCCTCCACTCCTTTCAGGTATTCTTTACCGATTTGCATATCAATCATAT AATGGATGTCCACGAGGTCTCTTCCGATCTACCAAGACAGGACGTAC GCTGCAGGTCGACG | Deletion, F |
| *ENP2* | *ECL1* | CATGAGTAGGTTAAAGGCAAGTCGAAGAAGATACGAAGCAAGTGAA GCCGTTTACGGTGTCGGTCTCGTAGGGCCACGTACTTACAACATAAT CGATGAATTCGAGCTCGT | Deletion, R |
| *ENP2* | *ECL1* | TACATTCACATCCATACTTGGACAC | Verification, F |
| *UTP21* | *VIP1* | CATACAAATTCAAAAGCATCTCGTAGCATATTAATATATTGCAGAAG GTCATGGATGTCCACGAGGTCTCTATGGAGACGCACATTCCTGCCGT ACGCTGCAGGTCGACG | Deletion, F |
| *UTP21* | *VIP1* | GAGTAAATACTTATTTAGTTTTGGGTTACTAAATTAAAAATTGGGTGT GATCACTACGGTGTCGGTCTCGTAGTGAGATACATCACGCACATCAT CGATGAATTCGAGCTCGT | Deletion, R |
| *UTP21* | *VIP1* | CTAACTCGCTATAGCCTTTTCAGTG | Verification, F |
| *SEC22* | *BOP2* | GGCTGTGATTTCGAAAGTTATATATTCTCCGGGTAGAAGTGAAAAGG ATG GATGTCCACGAGGTCTCTCCATCGAGATAACGCTAAGCCGTACGCTG CAGGTCGACG | Deletion, F |
| *SEC22* | *BOP2* | GAAATTTATTTAATTATATATGCTAGTACAACACGTTTGGTTGAAAAT TAGACCTCACGGTGTCGGTCTCGTAGAGTAATCATACATACCGCGCA TCGATGAATTCGAGCTCGT | Deletion, R |
| *SEC22* | *BOP2* | CTGTTAATGCCTCTTCATGATTTCT | Verification, F |
| *BOP2* | *SEC22* | CCAACAAAAACCCTGACAGTGACACCCCGTTACACACTCACAATTAA | Deletion, F |

| | | | |
|---|---|---|---|
| | | GTAGGGATGGATGTCCACGAGGTCTCTTCCAGAGCTAGGAAGTAAG CCGTACGCTGCAGGTCGACG | |
| *BOP2* | *SEC22* | CACTTGGACCAAATTGATCGGGATTTGTGATGTGGGATGATGGGGT GACGTCTACGGTGTCGGTCTCGTAGTGAGGATAAAGAACTCGCGCAT CGATGAATTCGAGCTCGT | Deletion, R |
| *BOP2* | *SEC22* | AATTTTCAACCAAACGTGTTGTACT | Verification, F |
| *SPB4* | *DEG1* | CTCGAGGTGCCCACATGCAATCTTTACTGCCCTACTATAACCTCCCTT GAATGGATGTCCACGAGGTCTCTTTTAGCCAGAGCACACGCAGCGTA CGCTGCAGGTCGACG | Deletion, F |
| *SPB4* | *DEG1* | GAAAAGAAATATAGTCTTCAAGGTTATATTATACAGGTTTATATATTA TTTTACGGTGTCGGTCTCGTAGTACGGTAGACCATTGCCGAGATCGA TGAATTCGAGCTCGT | Deletion, R |
| *SPB4* | *DEG1* | CCTTATCCAGGGAAGTAAAGAAAAC | Verification, F |
| *SEC5* | *TRM82* | GCTAAAGTATAGCAAGGTTTGTCAATTTGAAGCAGTCCGCGTGTTTT ACAATGGATGTCCACGAGGTCTCTGGAGAGCGACCCTTCATTCTCGT ACGCTGCAGGTCGACG | Deletion, F |
| *SEC5* | *TRM82* | CGGGTACGTATCTTTTTTTATATTAGTTATAAAGATGTACTTACTACCT TTTTATTTCACGGTGTCGGTCTCGTAGTGTATCAACAGGGTGGAAAC ATCGATGAATTCGAGCTCGT | Deletion, R |
| *SEC5* | *TRM82* | TTACTGATATTTTGATCCAGGAGGA | Verification, F |
| *ATG20* | *TRM3* | GAATTGTTTAACGAACGTTACAAACTCTATACCTTTTTTTACCAGCAA AATGGATGTCCACGAGGTCTCTCCGATTAGAGGTTGACAGATCGTAC GCTGCAGGTCGACG | Deletion, F |
| *ATG20* | *TRM3* | GTTTACATACTAAAAACTTCATGGTTAACAAAAGTGGGAATGAAACC TGATTCTTACGGTGTCGGTCTCGTAGCACTGACTTCGAGGTCGTGTAT CGATGAATTCGAGCTCGT | Deletion, R |
| *ATG20* | *TRM3* | ATCTCTTCATTTTGGAGATGACTTG | Verification, F |
| *IMP2'* | *RRD1* | GCAAAGTGCAAAAGAACGCACATATGAACAAGCATTAAACGAGCAA AGAAATGGATGTCCACGAGGTCTCTTACGATTGCAGAGATTGCTGCG TACGCTGCAGGTCGACG | Deletion, F |
| *IMP2'* | *RRD1* | CGAAAATAGAAGTCATAATGCTTGTCATACACATTTATATGTTTAATT AATATTACGGTGTCGGTCTCGTAGACATGGACGACATTGTGCTCATC GATGAATTCGAGCTCGT | Deletion, R |
| *IMP2'* | *RRD1* | CTTTTCTTCTTGCTACGACCTCTTT | Verification, F |
| *RRD1* | *IMP2'* | GGAAAGGGTGAGTACCAAAAGAACCAACAAGAGAAACAACCAAGTA CGCAATGGATGTCCACGAGGTCTCTTGATCTCTACGCACTTGCTGCGT ACGCTGCAGGTCGACG | Deletion, F |
| *RRD1* | *IMP2'* | GCTGTATATAAGTATGTGTTGCTAAAAAGGAATTAG TGCAGTGATTATTGGTCACGGTGTCGGTCTCGTAGCATATCCACTAA GGTTGCTCATCGATGAATTCGAGCTCGT | Deletion, R |
| *RRD1* | *IMP2'* | TGCGATATCAACGTAGTGTACTTGT | Verification, F |
| *UBX2* | *TRM9* | GAACAGAGATGAGGTCTCGAAGAGCCAAGAAATAAAAGGTTAAGAA CCAACATGGATGTCCACGAGGTCTCTATATGTGACCGCACCTCTGGC GTACGCTGCAGGTCGACG | Deletion, F |
| *UBX2* | *TRM9* | CTATCGTAATACCTGCTGCTACAAAATACACTGTCTACCTATATATCAC CTTCACGGTGTCGGTCTCGTAGTGTATCAACAGGGTGGAAACATCGA TGAATTCGAGCTCGT | Deletion, R |
| *UBX2* | *TRM9* | GGATGGTGTCCAAAGGACCTTGA | Verification, F |
| *KEG1* | *IRC6* | CTGATGCAGCAAGATAGCAAGTATATATACGCAAAAATACCAATCTA CCATGGATGTCCACGAGGTCTCTACACAGGCATGGAATGTATCCGTA CGCTGCAGGTCGACG | Deletion, F |
| *KEG1* | *IRC6* | GCAATCTTATTGGTTTAATACATATGTATACACATATACATATCTGTAC ATACTCACGGTGTCGGTCTCGTAGAATCGACAGGGAGACCAGTCATC GATGAATTCGAGCTCGT | Deletion, R |
| *KEG1* | *IRC6* | TTCCATTCCTATCACTTTGACTTTC | Verification, F |

| | | | |
|---|---|---|---|
| *NRP1* | *SFA1* | CTTCTACAAAATCTCCAAGTAAAGAAGGAATATAAGTAATATAAGTACAATGGATGTCCACGAGGTCTCTCTTTCGGACGTATGTGCAGTCGTACGCTGCAGGTCGACG | Deletion, F |
| *NRP1* | *SFA1* | GTATTCCAGAAAATTTGAGTCATGCTTACTTAGTTTAATTAAGTACTCCTACGGTGTCGGTCTCGTAGCCTTGATGATAGAGGGCTTTATCGATGAATTCGAGCTCGT | Deletion, R |
| *NRP1* | *SFA1* | CAGAATTTGTTGGCCTATTTTCTTA | Verification, F |
| | | CTGCAGCGAGGAGCCGTAAT | Verification, Universal R (KanMX) |

**II. Primers for RT-qPCR**

| Primer sequence | Description |
|---|---|
| GCTTTTCAAGATGAGCTGAGCC | Forward primer to the extended 5' UTR of the SPB4 |
| TCCAAGCATCGACAATTCCCA | Forward primer to the CDS of SPB4 |
| AGTACCAACCAGGAGCTGAC | Reverse primer to the CDS of SPB4 |
| AAAATGGAGCACAGAATGCGT | Forward primer to the extended 5' UTR of the SPB4 |
| CCCAATCGTAATTCGCCACC | Reverse primer to the CDS of APC4 (paired with previous one) |
| CGCTGTTTTCTTGTCGGTGG | Forward primer to the CDS of the APC4 |
| TAAGGTGTTGCCAGTTCCCG | Reverse primer to the CDS of APC4 (paired with previous one) |

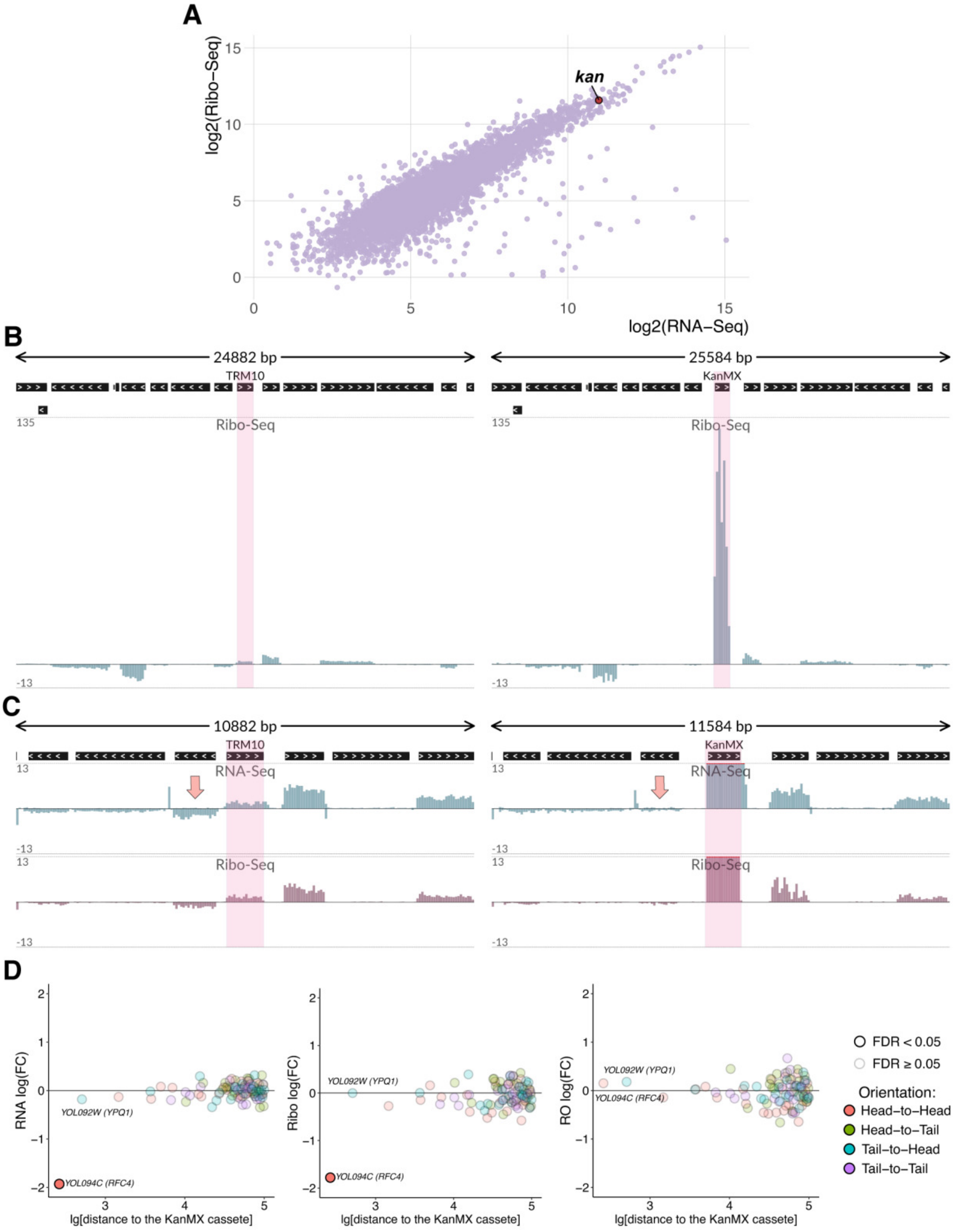


**Figure S1**. KanMX cassette used to perform gene deletions is highly expressed within the mutated locus and affects the expression of the adjacent gene. (A) *kan* is among the top 1% most highly expressed genes in *trm10Δ* yeast strain, both at the transcriptional and translational levels. RNA-Seq (X-axis, log-scale) and Ribo-Seq (Y-axis, log-scale) read coverage of *S. cerevisiae* CDSs, obtained for a reconstructed genome annotation of the *trm10Δ* strain. (B) Excessive *kan* gene expression at the translational level in the mutant strain, as compared to other genes in the vicinity. Ribo-Seq data in *TRM10* locus (including 15 genes in the vicinity) in *wt* and *trm10Δ* strains are shown as normalized genomic tracks, positive and negative values correspond to the direct and reverse complementary DNA strands, respectively. The highlighted frames indicate the *TRM10* CDS (*wt* strain) and the KanMX module (*trm10Δ* strain). (C) RNA-Seq and Ribo-Seq data in *TRM10* locus (including 7 genes in the vicinity) in *wt* and *trm10Δ* strains are shown as normalized genomic tracks at an enlarged scale. Note that the KanMX coverage is truncated to maintain the same scale of the Y-axis as for the deleted gene. (D) Differential expression of genes in the mutated locus (±100 kb from the KanMX module) at the level of transcription (RNA-Seq, left subpanel), translation (Ribo-Seq, middle subpanel), and translation efficiency (RO, right subpanel), in *trm10*Δ strain compared to *wt*. FDR: false discovery rate.

**Figure S2** – see separate file.

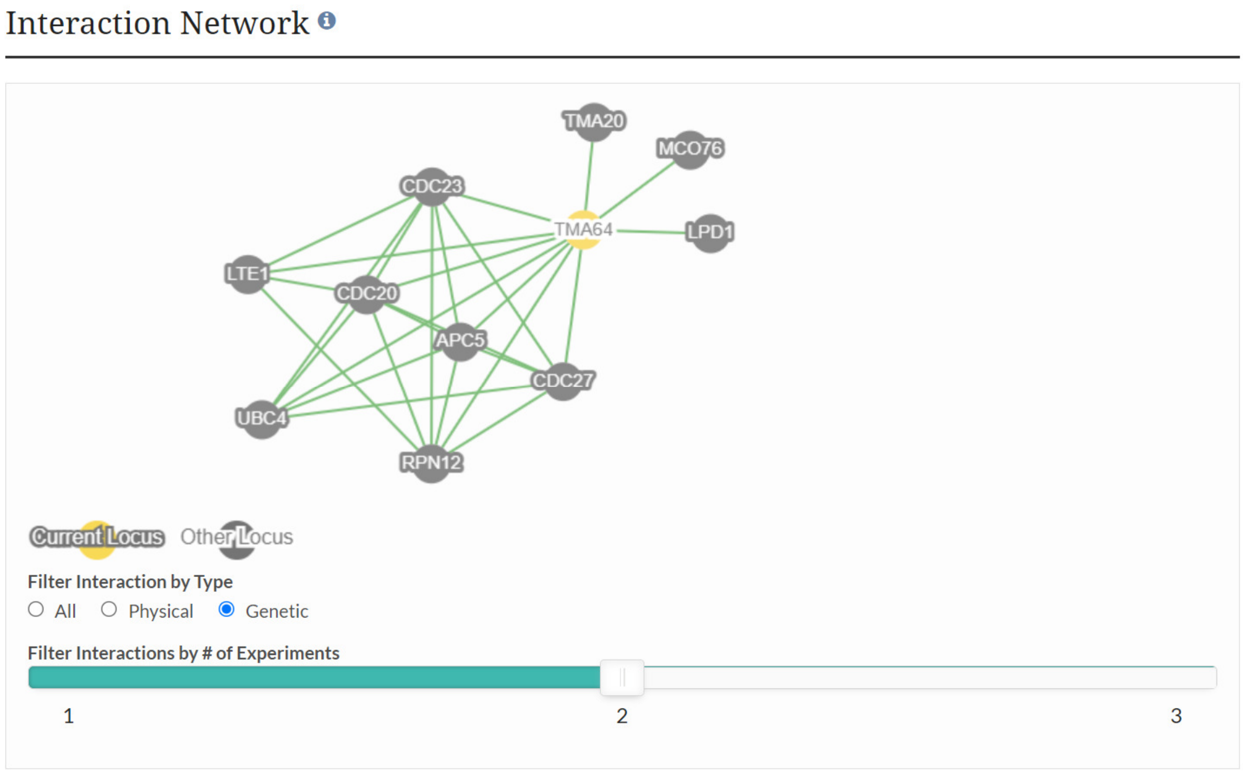


**Figure S3**. *TMA64* genetic interactions listed in SGD (https://www.yeastgenome.org, Cherry et al., 2012) and documented in at least 2 data sources. 7 out of 10 interacting genes encode either APC/C subunits (APC5, CDC20, CDC23, CDC27) or other components of the mitotic checkpoint (UBI4 – key ubiquitin-conjugating enzyme E2 for the APC/C; RPN12 – a subunit of the 19S regulatory particle of the 26S proteasome lid; LTE1 – a GDP/GTP exchange factor required for mitotic exit at low temperatures).

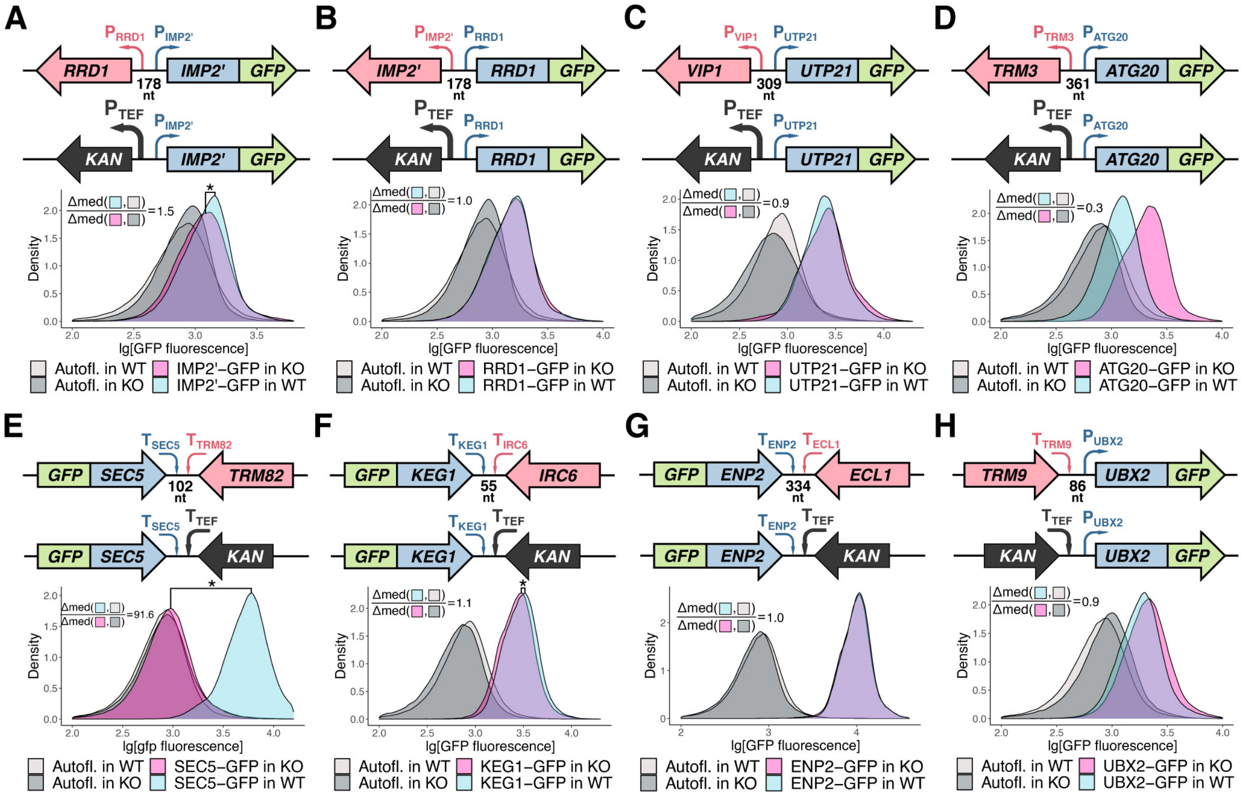


**Figure S4**. Gene replacement with the KanMX module induces changes in the abundance of GFP-tagged proteins encoded by neighboring genes. (A-H) Eight cases with different mutual gene orientations (schematically shown in upper subpanels) were analyzed. For each case, 4 yeast strains were assayed by flow cytometry: 2 strains with the GFP-tagged neighboring gene (as shown in the schemes) and 2 corresponding strains without the GFP-tag (to take into account possible autofluorescence changes between the *KO* strains and *wt*). The density plots show distribution of the GFP fluorescence (log-scale, combined results from at least three independent experiments). Effect size estimates (Δmed ratios) are shown on the plots, see Materials and Methods for details.*one-sided U-test p-value $<10^{-20}$.
