## Supplementary Figure 2 for "A standard knockout procedure alters expression of adjacent loci at the translational level"

### SUPPLEMENTARY DATA

**Figure S2**. KanMX cassette affects the expression of adjacent genes. (A) Scatterplots illustrating differential expression of genes in the mutated locus (±100 kb from the KanMX module) in the knockout strain as compared to *wt*. Subpanels: RNA-Seq (left), Ribo-Seq (middle), ribosome occupancy (right). X-axis: log-distance, base pairs; Y-axis: fold change. FDR: false discovery rate. (B) RNA-Seq and Ribo-Seq normalized read coverage in the locus including entire CDSs within a region of approximately ±100 kb from the target gene. Subpanels: *wt* (left), *KO* strain (right). Positive and negative values correspond to the direct and reverse complementary DNA strands, respectively. Note that the KanMX coverage track is truncated to maintain the same Y-axis scale as for the target gene.

# S2-1

**A**

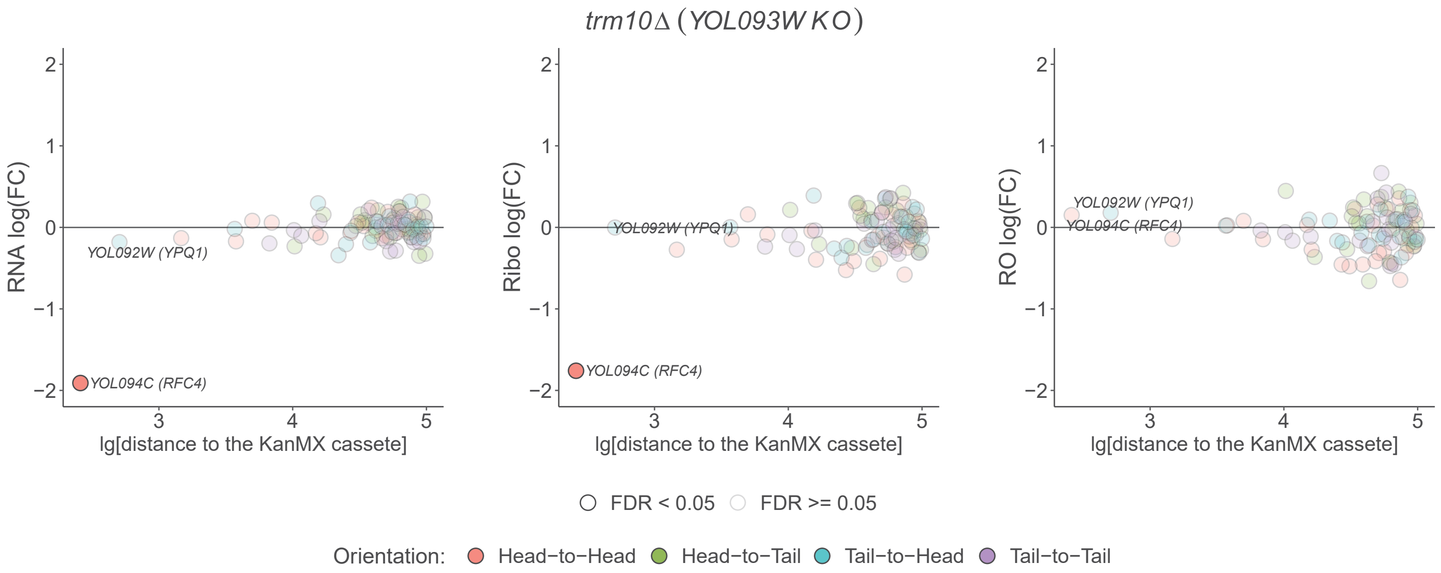

trm10Δ (YOL093W KO)
RNA log(FC)
Ribo log(FC)
RO log(FC)
lg[distance to the KanMX cassete]
YOL092W (YPQ1)
YOL094C (RFC4)
FDR < 0.05
FDR >= 0.05
Orientation:
Head−to−Head
Head−to−Tail
Tail−to−Head
Tail−to−Tail


**B**

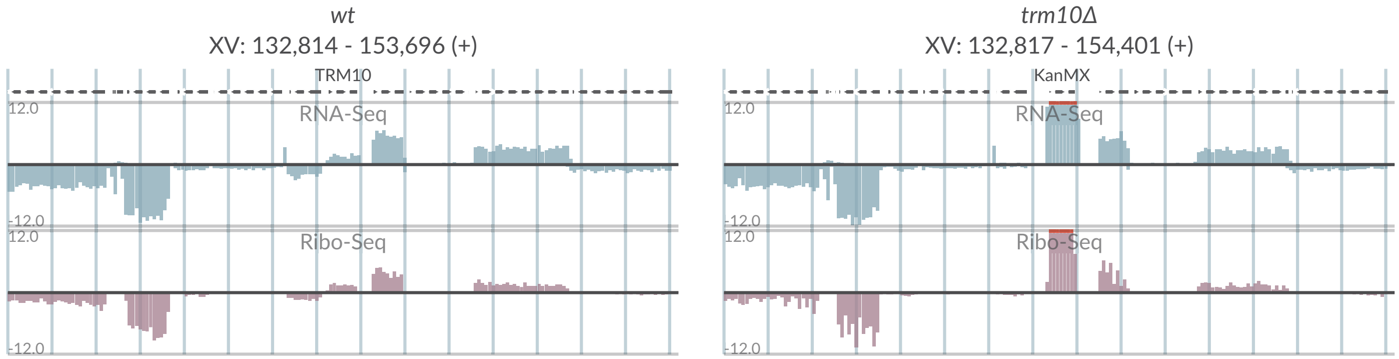

wt
XV: 132,814 - 153,696 (+)
TRM10
trm10Δ
XV: 132,817 - 154,401 (+)
KanMX
RNA-Seq
Ribo-Seq
12.0
-12.0

# S2-2

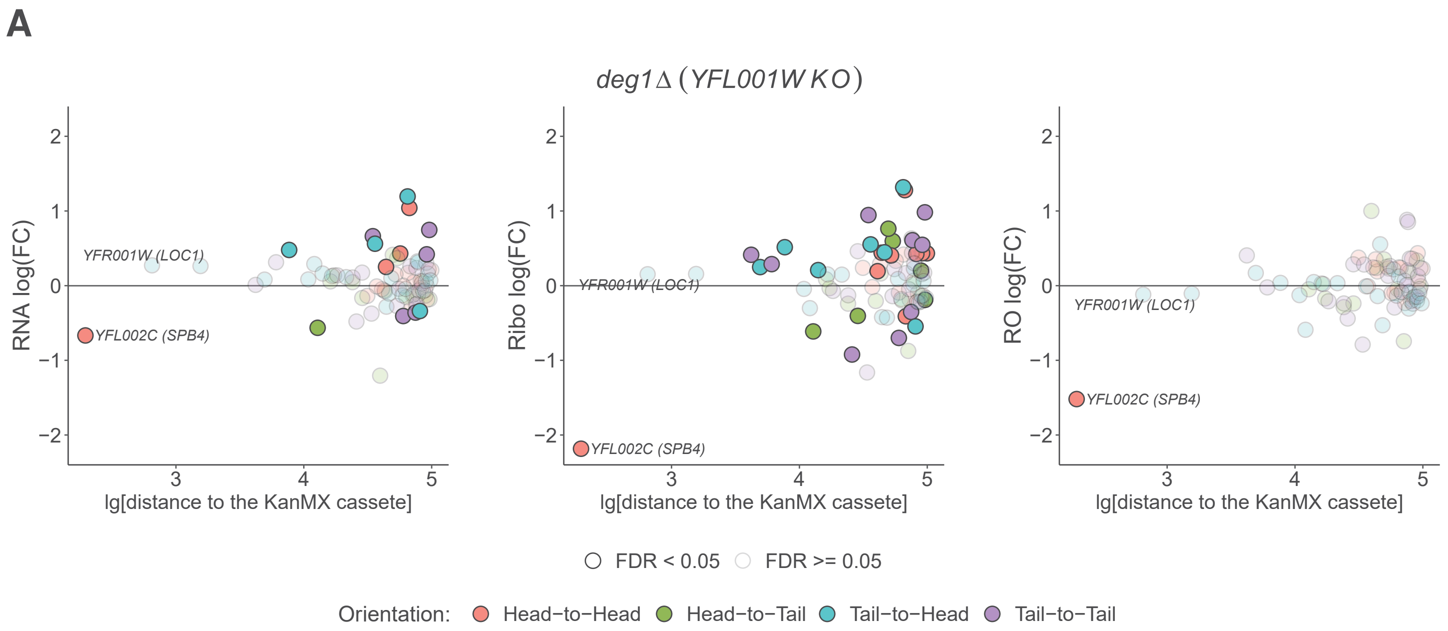
A
*deg1∆ ( YFL001W KO )*
RNA log(FC)
Ribo log(FC)
RO log(FC)
YFR001W (LOC1)
YFL002C (SPB4)
lg[distance to the KanMX cassete]
FDR < 0.05
FDR >= 0.05
Orientation:
Head−to−Head
Head−to−Tail
Tail−to−Head
Tail−to−Tail

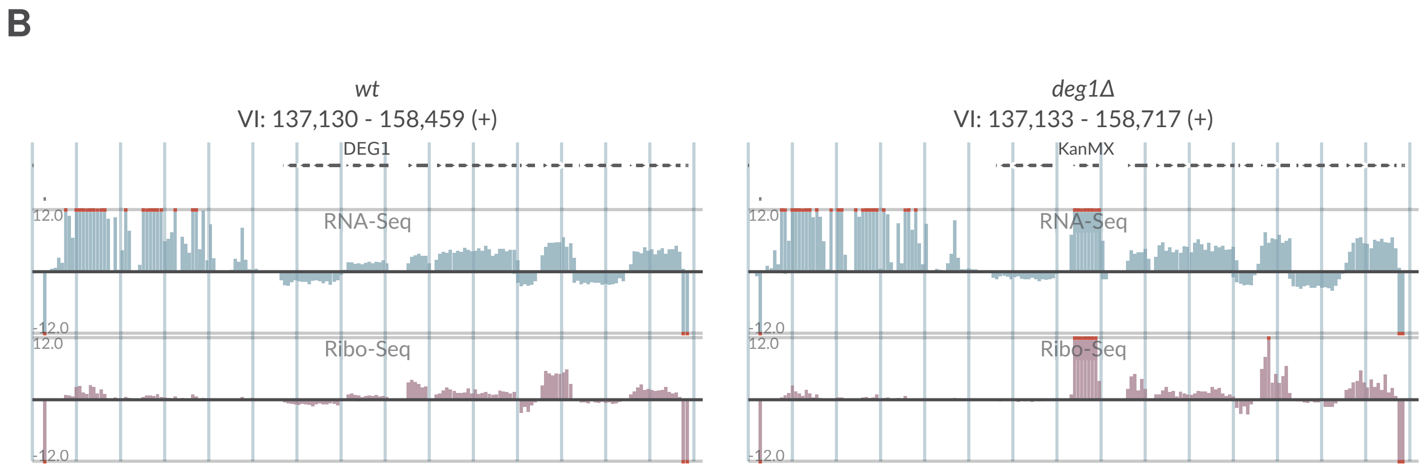
B
*wt*
VI: 137,130 - 158,459 (+)
DEG1
RNA-Seq
Ribo-Seq
*deg1Δ*
VI: 137,133 - 158,717 (+)
KanMX
RNA-Seq
Ribo-Seq

# S2-3

## A

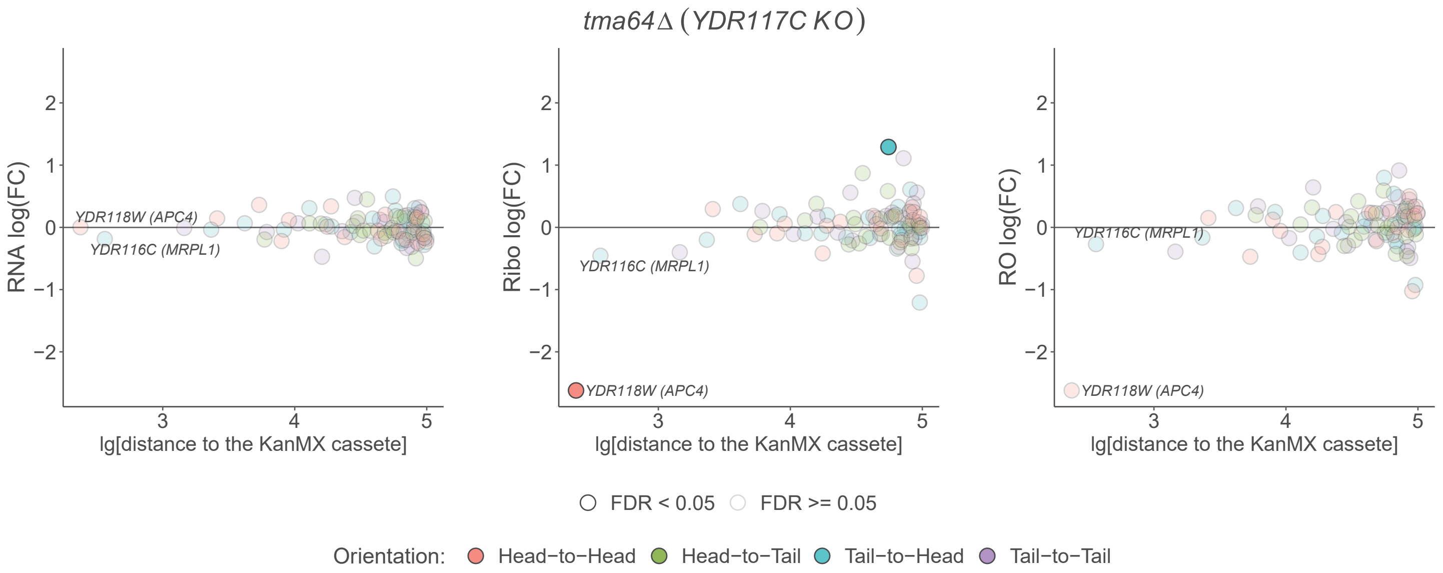

tma64Δ (YDR117C KO)
RNA log(FC)
Ribo log(FC)
RO log(FC)
YDR118W (APC4)
YDR116C (MRPL1)
lg[distance to the KanMX cassete]
FDR < 0.05
FDR >= 0.05
Orientation: Head−to−Head Head−to−Tail Tail−to−Head Tail−to−Tail


## B

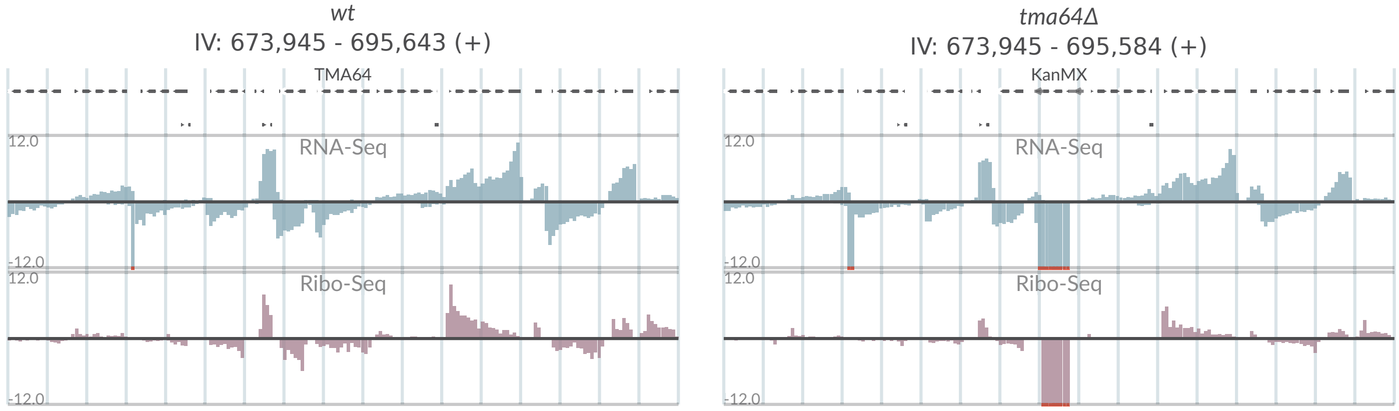

wt
IV: 673,945 - 695,643 (+)
TMA64
tma64Δ
IV: 673,945 - 695,584 (+)
KanMX
RNA-Seq
Ribo-Seq
12.0
-12.0

# S2-4

## A

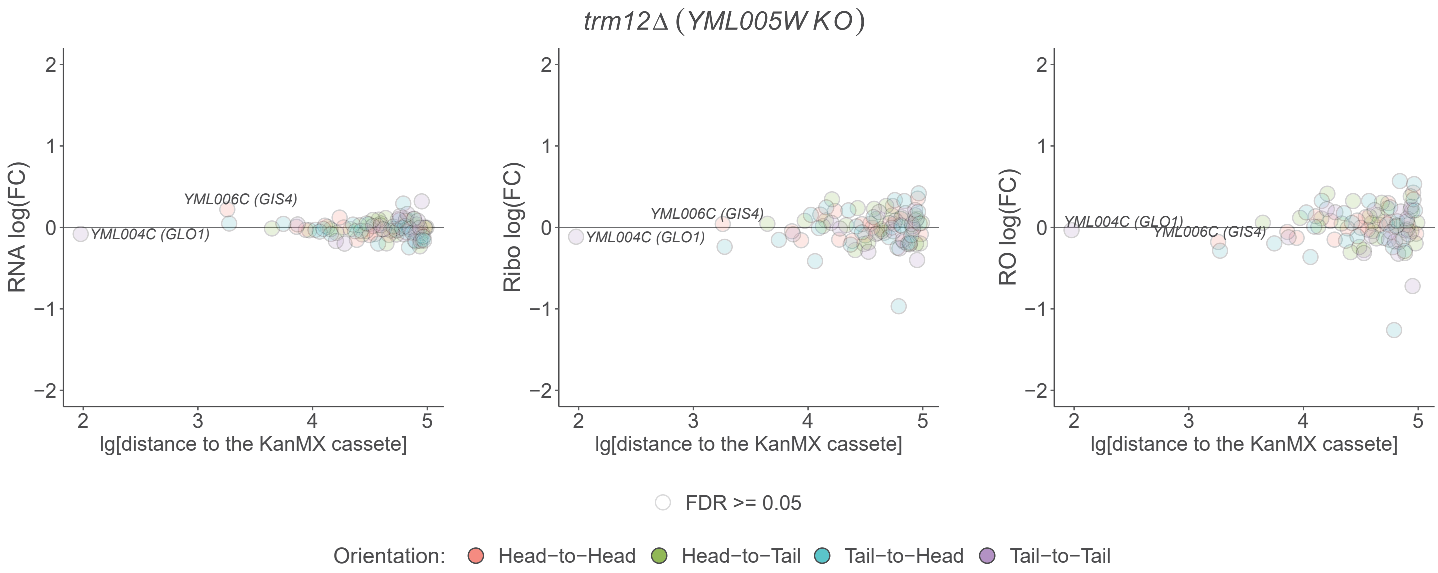

*trm12Δ (YML005W KO)*
RNA log(FC)
Ribo log(FC)
RO log(FC)
YML006C (GIS4)
YML004C (GLO1)
lg[distance to the KanMX cassete]
FDR >= 0.05
Orientation: Head−to−Head Head−to−Tail Tail−to−Head Tail−to−Tail


## B

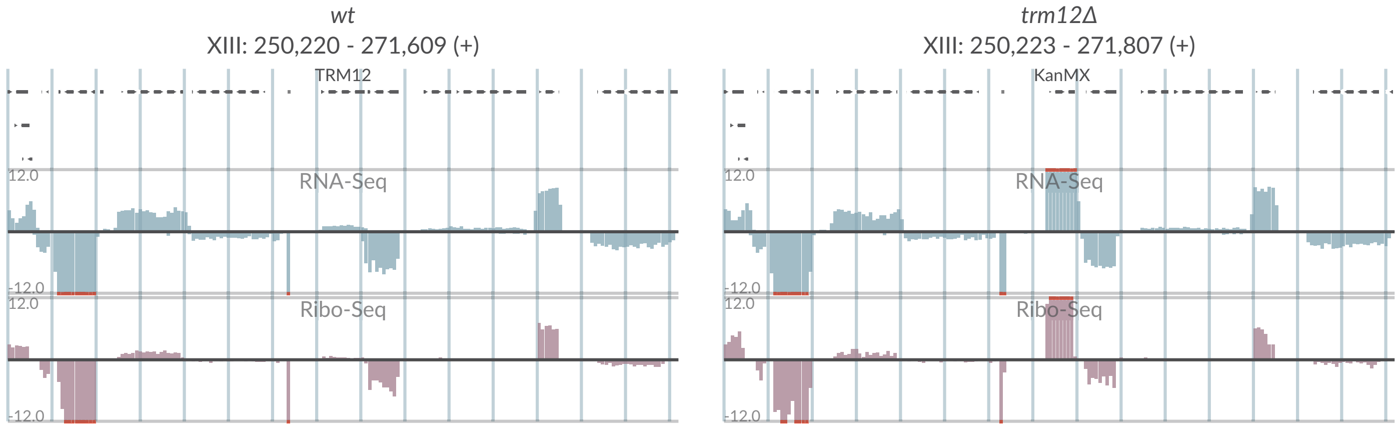

*wt*
XIII: 250,220 - 271,609 (+)
TRM12
*trm12Δ*
XIII: 250,223 - 271,807 (+)
KanMX
RNA-Seq
Ribo-Seq
12.0
-12.0

# S2-5

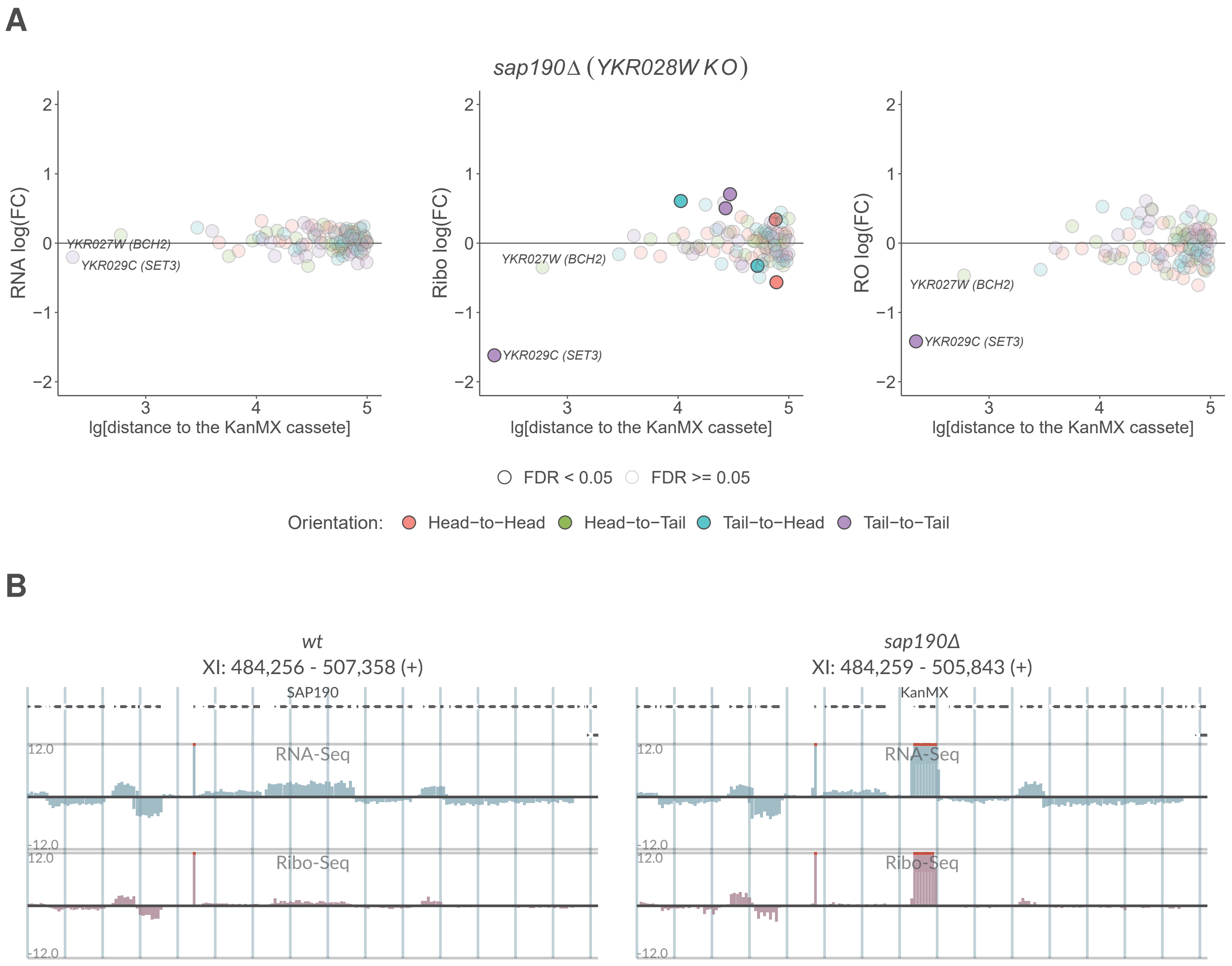
A
sap190Δ (YKR028W KO)
RNA log(FC)
Ribo log(FC)
RO log(FC)
lg[distance to the KanMX cassete]
YKR027W (BCH2)
YKR029C (SET3)
FDR < 0.05
FDR >= 0.05
Orientation:
Head−to−Head
Head−to−Tail
Tail−to−Head
Tail−to−Tail
B
wt
XI: 484,256 - 507,358 (+)
SAP190
sap190Δ
XI: 484,259 - 505,843 (+)
KanMX
RNA-Seq
Ribo-Seq
12.0
-12.0

# S2-6

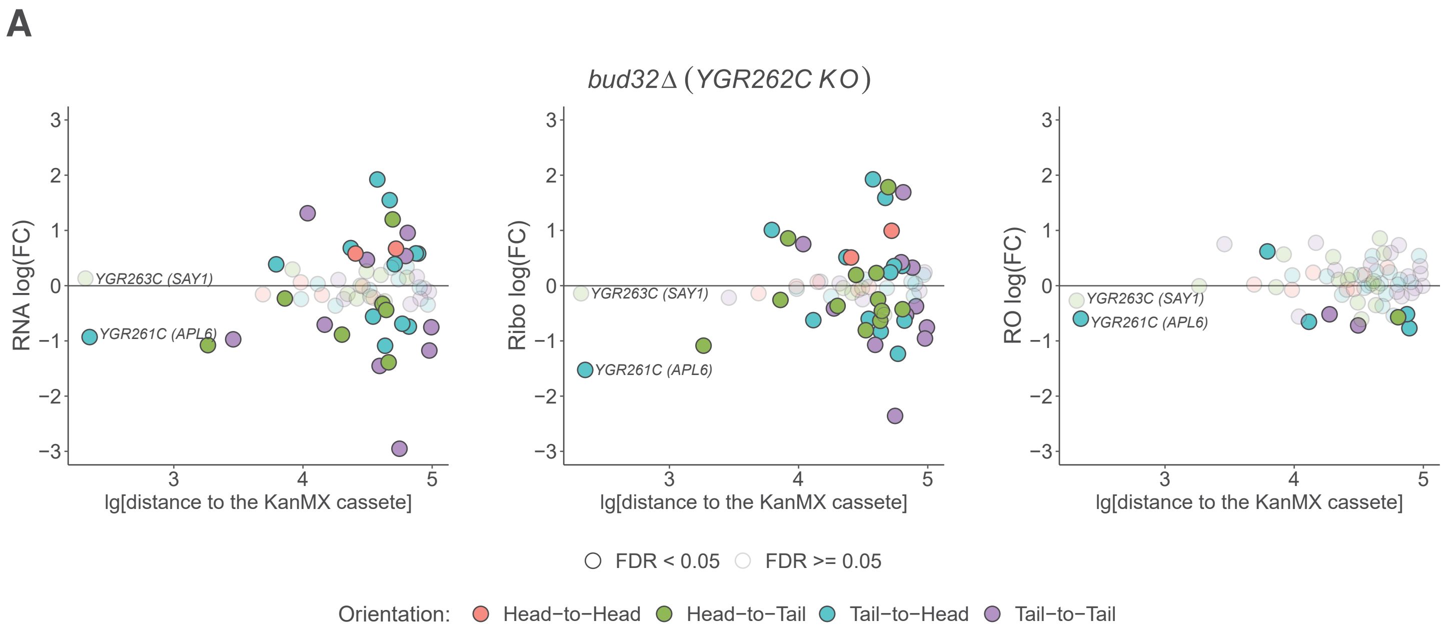
A
bud32Δ (YGR262C KO)
RNA log(FC)
Ribo log(FC)
RO log(FC)
YGR263C (SAY1)
YGR261C (APL6)
lg[distance to the KanMX cassete]
FDR < 0.05
FDR >= 0.05
Orientation:
Head−to−Head
Head−to−Tail
Tail−to−Head
Tail−to−Tail

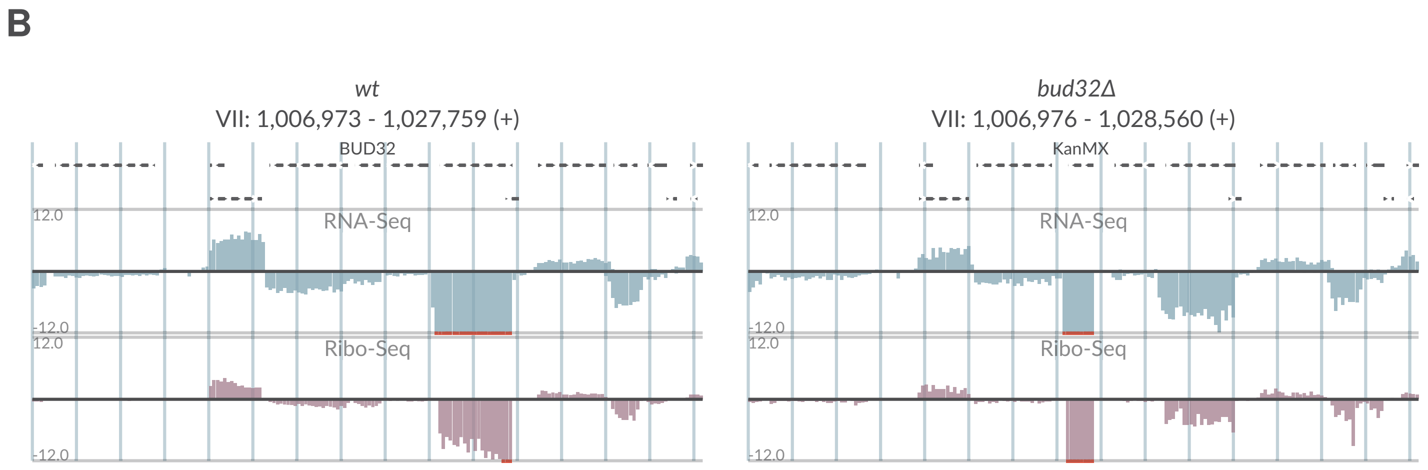
B
wt
VII: 1,006,973 - 1,027,759 (+)
BUD32
bud32Δ
VII: 1,006,976 - 1,028,560 (+)
KanMX
RNA-Seq
Ribo-Seq
12.0
-12.0

# S2-7

**A**

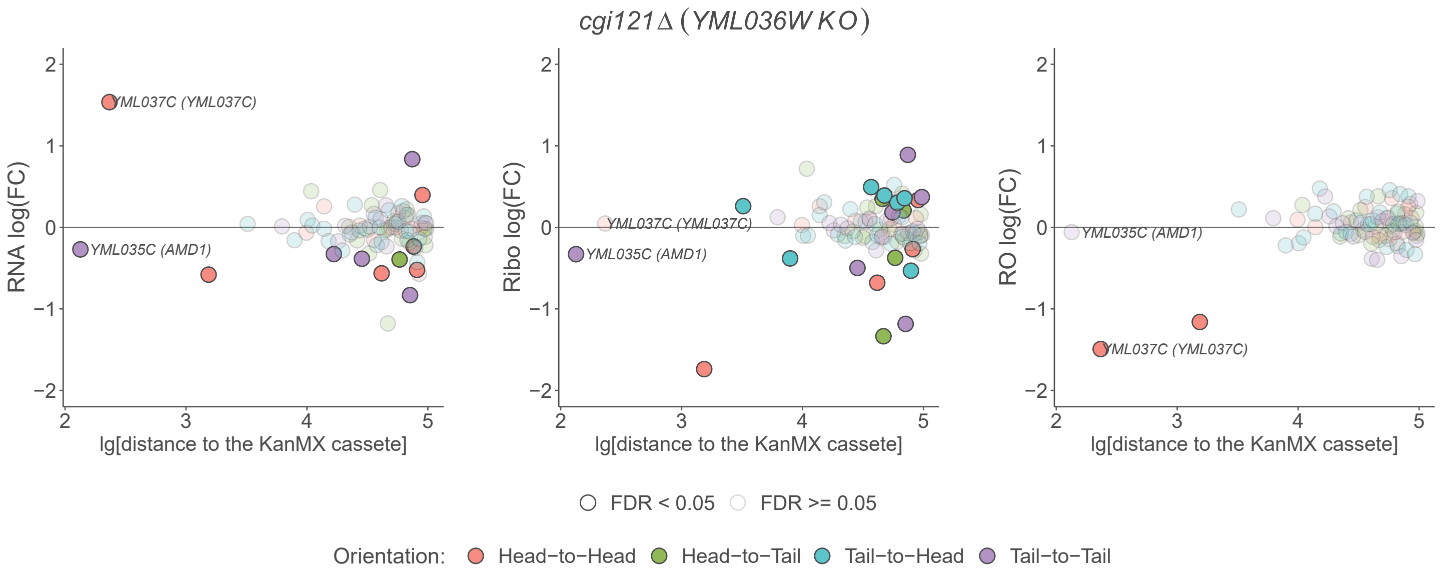

cgi121Δ (YML036W KO)
RNA log(FC)
Ribo log(FC)
RO log(FC)
lg[distance to the KanMX cassete]
YML037C (YML037C)
YML035C (AMD1)
FDR < 0.05
FDR >= 0.05
Orientation:
Head−to−Head
Head−to−Tail
Tail−to−Head
Tail−to−Tail


**B**

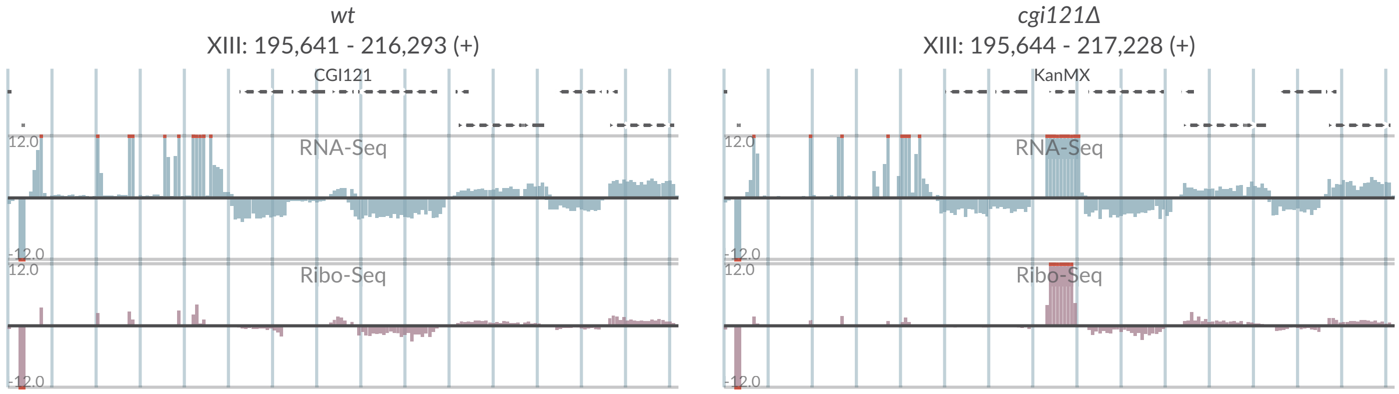

wt
XIII: 195,641 - 216,293 (+)
CGI121
RNA-Seq
Ribo-Seq
cgi121Δ
XIII: 195,644 - 217,228 (+)
KanMX
RNA-Seq
Ribo-Seq

# S2-8

## A

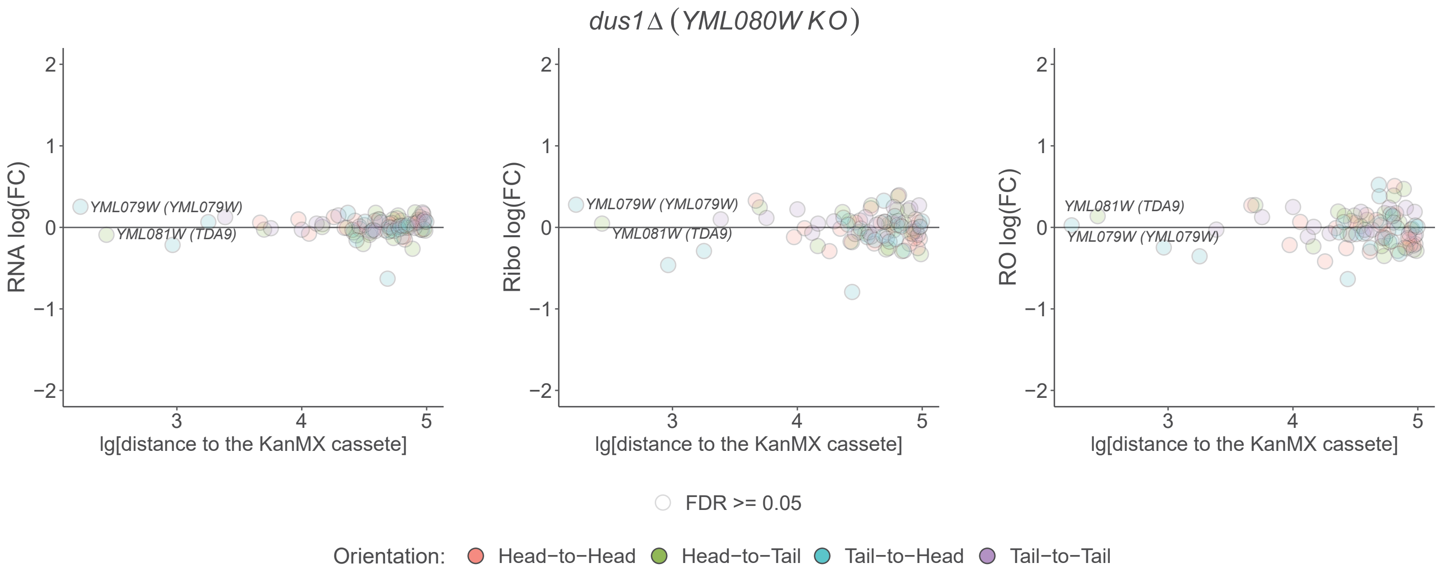
*dus1Δ (YML080W KO)*
RNA log(FC)
Ribo log(FC)
RO log(FC)
*YML079W (YML079W)*
*YML081W (TDA9)*
lg[distance to the KanMX cassete]
FDR >= 0.05
Orientation: Head−to−Head Head−to−Tail Tail−to−Head Tail−to−Tail

## B

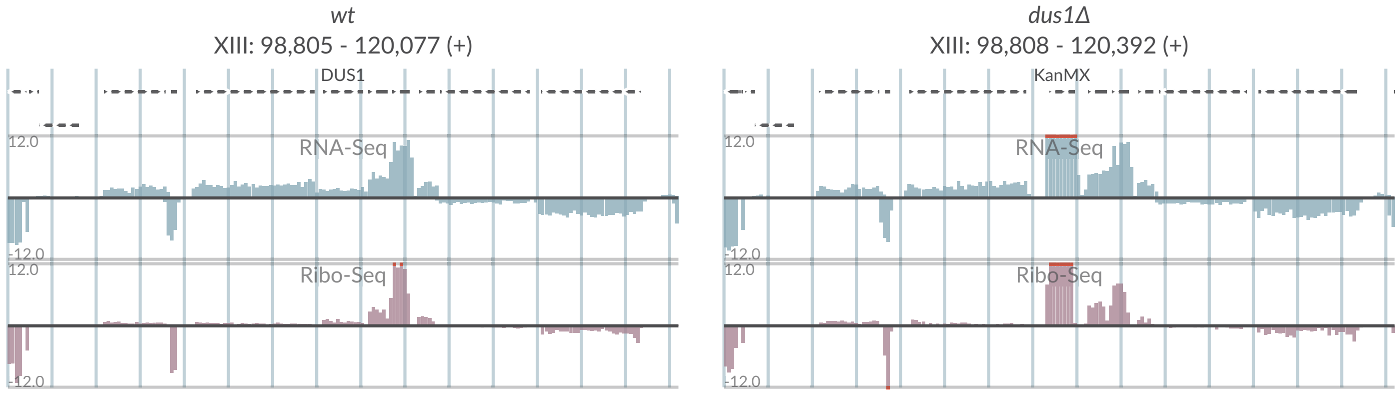
*wt*
XIII: 98,805 - 120,077 (+)
DUS1
RNA-Seq
Ribo-Seq
*dus1Δ*
XIII: 98,808 - 120,392 (+)
KanMX
RNA-Seq
Ribo-Seq

# S2-9

## A

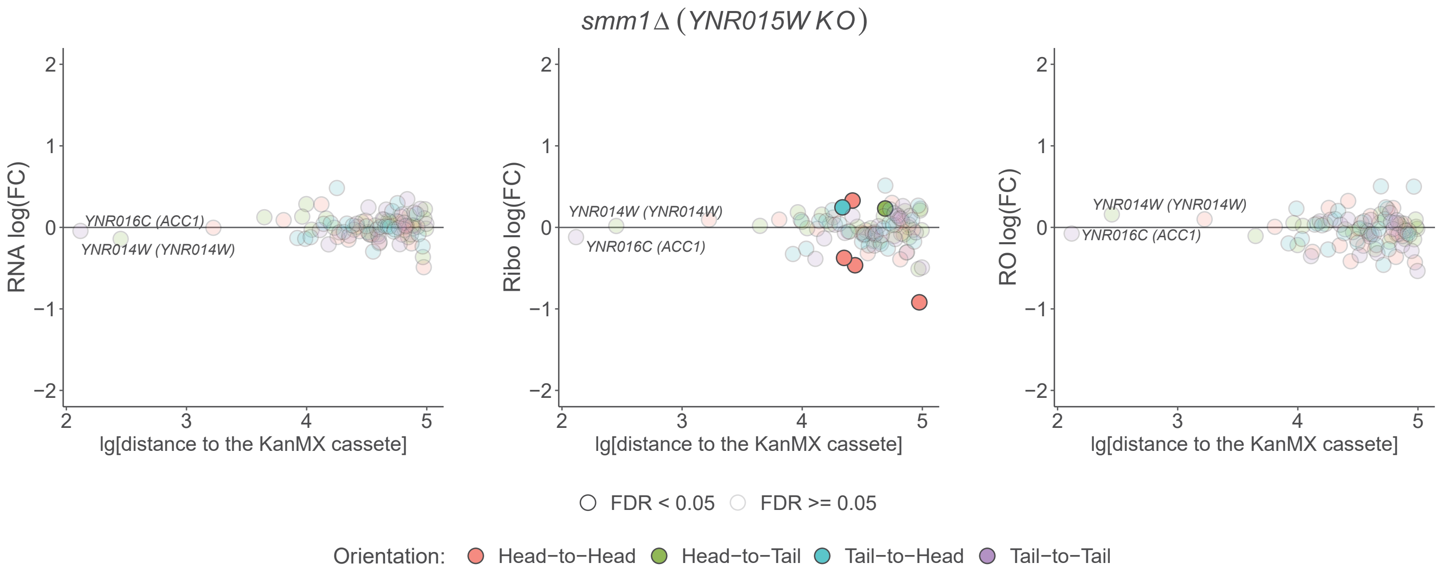

smm1Δ (YNR015W KO)
RNA log(FC)
Ribo log(FC)
RO log(FC)
YNR016C (ACC1)
YNR014W (YNR014W)
lg[distance to the KanMX cassete]
FDR < 0.05
FDR >= 0.05
Orientation:
Head−to−Head
Head−to−Tail
Tail−to−Head
Tail−to−Tail


## B

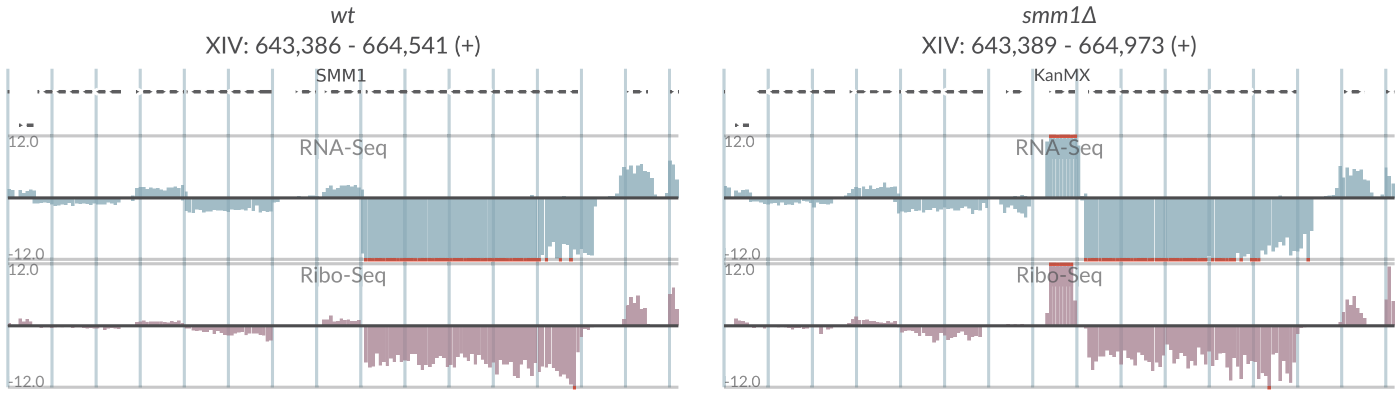

wt
XIV: 643,386 - 664,541 (+)
SMM1
smm1Δ
XIV: 643,389 - 664,973 (+)
KanMX
RNA-Seq
Ribo-Seq
12.0
-12.0

# S2-10

**A**

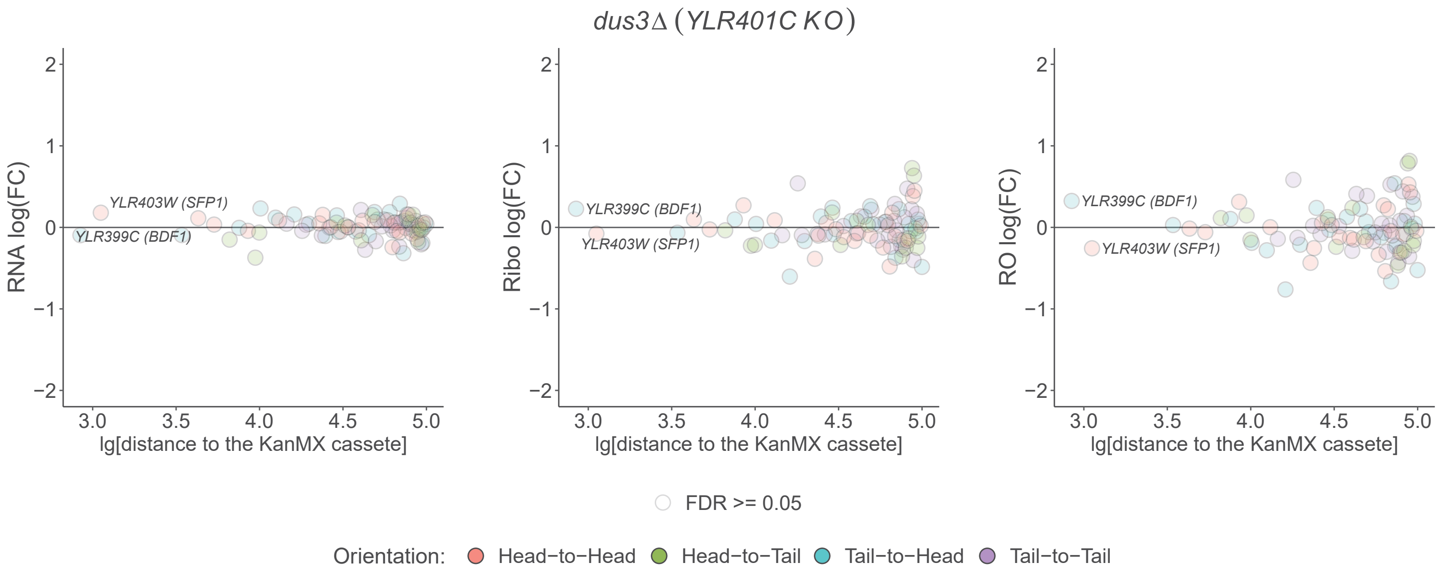
dus3Δ (YLR401C KO)
RNA log(FC)
Ribo log(FC)
RO log(FC)
YLR403W (SFP1)
YLR399C (BDF1)
lg[distance to the KanMX cassete]
FDR >= 0.05
Orientation: Head−to−Head Head−to−Tail Tail−to−Head Tail−to−Tail

**B**

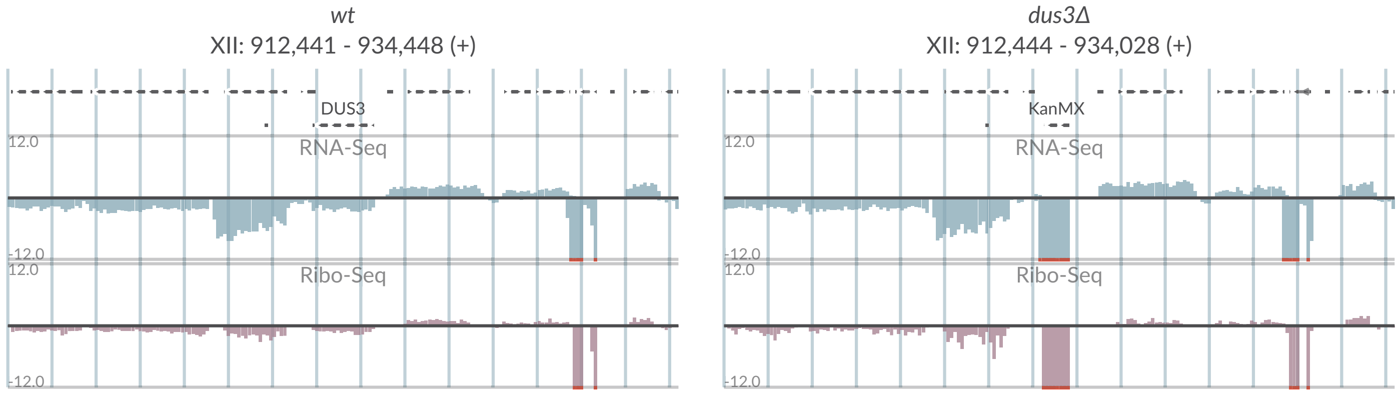
wt
XII: 912,441 - 934,448 (+)
DUS3
RNA-Seq
Ribo-Seq
dus3Δ
XII: 912,444 - 934,028 (+)
KanMX
RNA-Seq
Ribo-Seq
12.0
-12.0

# S2-11

**A**

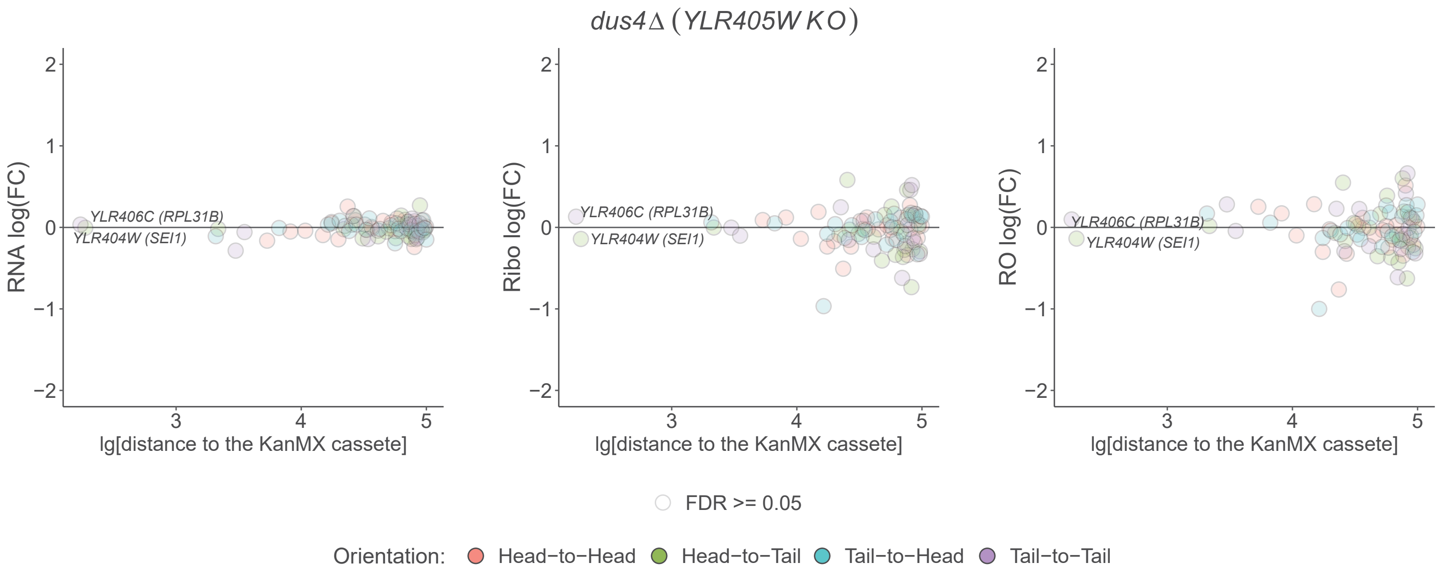

dus4Δ (YLR405W KO)
RNA log(FC)
Ribo log(FC)
RO log(FC)
YLR406C (RPL31B)
YLR404W (SEI1)
lg[distance to the KanMX cassete]
FDR >= 0.05
Orientation: Head−to−Head Head−to−Tail Tail−to−Head Tail−to−Tail


**B**

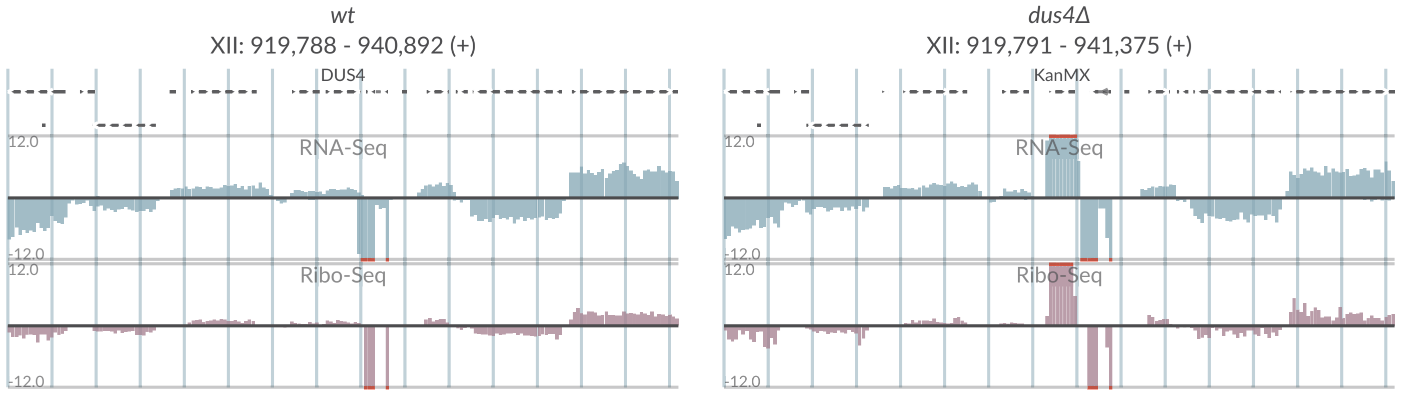

wt
XII: 919,788 - 940,892 (+)
DUS4
dus4Δ
XII: 919,791 - 941,375 (+)
KanMX
RNA-Seq
Ribo-Seq
12.0
-12.0

# S2-12

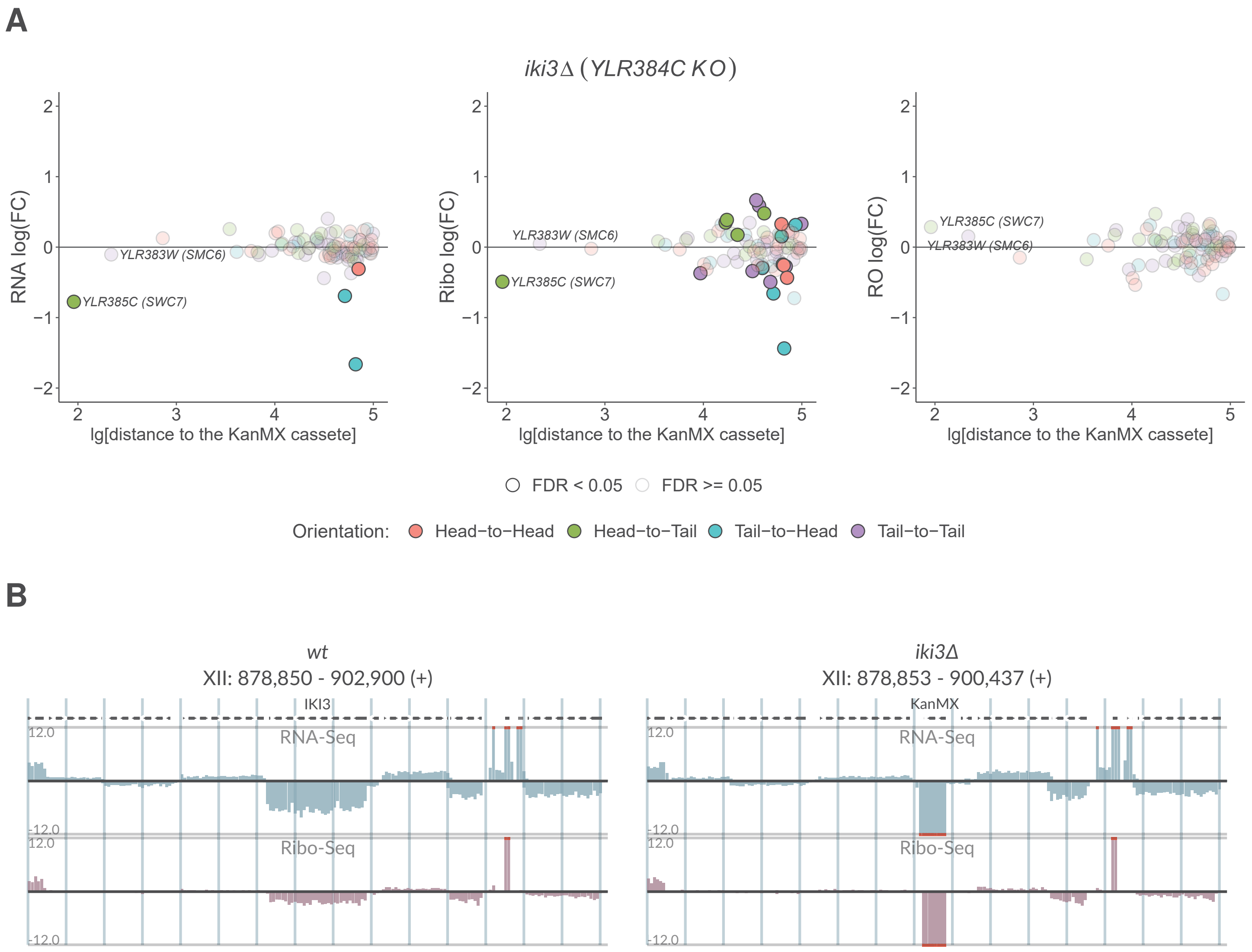
A
*iki3Δ (YLR384C KO)*
RNA log(FC)
Ribo log(FC)
RO log(FC)
lg[distance to the KanMX cassete]
YLR383W (SMC6)
YLR385C (SWC7)
FDR < 0.05
FDR >= 0.05
Orientation:
Head−to−Head
Head−to−Tail
Tail−to−Head
Tail−to−Tail
B
*wt*
XII: 878,850 - 902,900 (+)
IKI3
RNA-Seq
Ribo-Seq
*iki3Δ*
XII: 878,853 - 900,437 (+)
KanMX

## S2-13

**A**

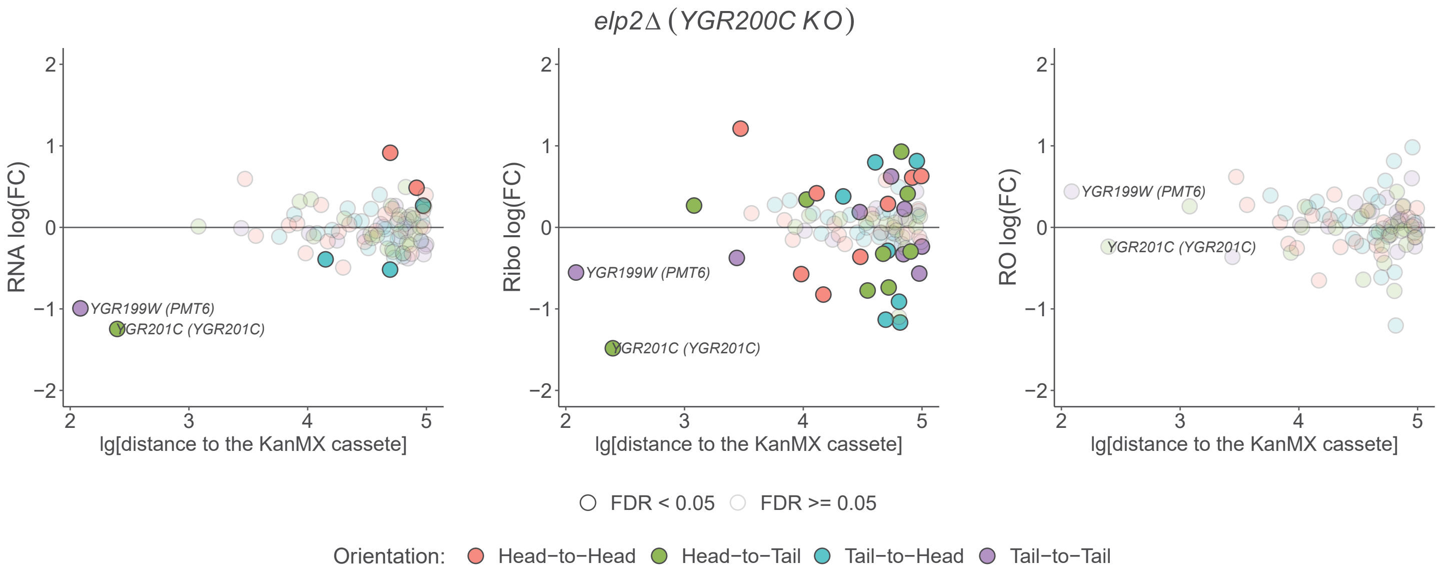

elp2Δ (YGR200C KO)
RNA log(FC)
Ribo log(FC)
RO log(FC)
lg[distance to the KanMX cassete]
YGR199W (PMT6)
YGR201C (YGR201C)
FDR < 0.05
FDR >= 0.05
Orientation:
Head−to−Head
Head−to−Tail
Tail−to−Head
Tail−to−Tail


**B**

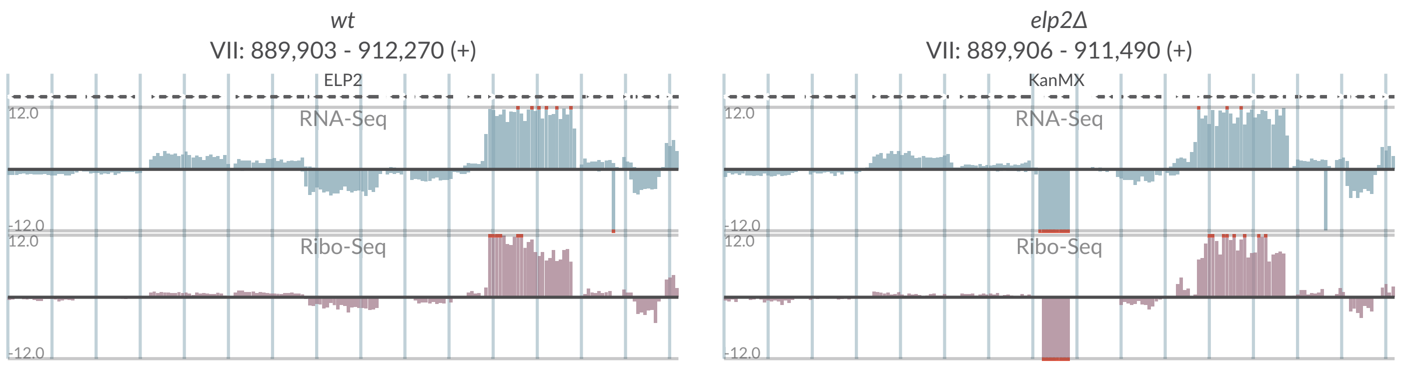

wt
VII: 889,903 - 912,270 (+)
ELP2
elp2Δ
VII: 889,906 - 911,490 (+)
KanMX
RNA-Seq
Ribo-Seq
12.0
-12.0

# S2-14

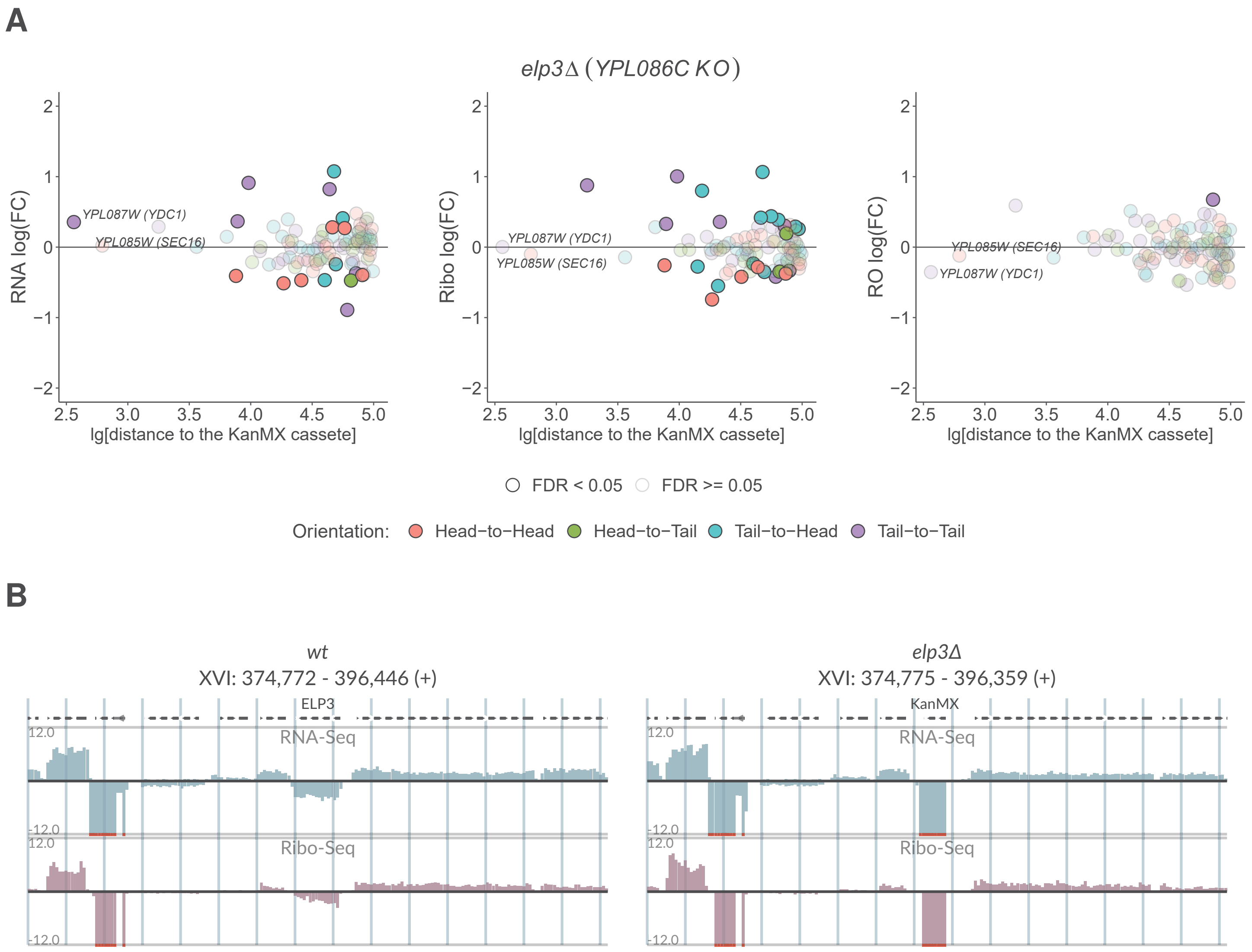
A
elp3Δ (YPL086C KO)
RNA log(FC)
Ribo log(FC)
RO log(FC)
YPL087W (YDC1)
YPL085W (SEC16)
lg[distance to the KanMX cassete]
FDR < 0.05
FDR >= 0.05
Orientation: Head−to−Head Head−to−Tail Tail−to−Head Tail−to−Tail
B
wt
XVI: 374,772 - 396,446 (+)
ELP3
elp3Δ
XVI: 374,775 - 396,359 (+)
KanMX
RNA-Seq
Ribo-Seq
12.0
-12.0

# S2-15

**A**

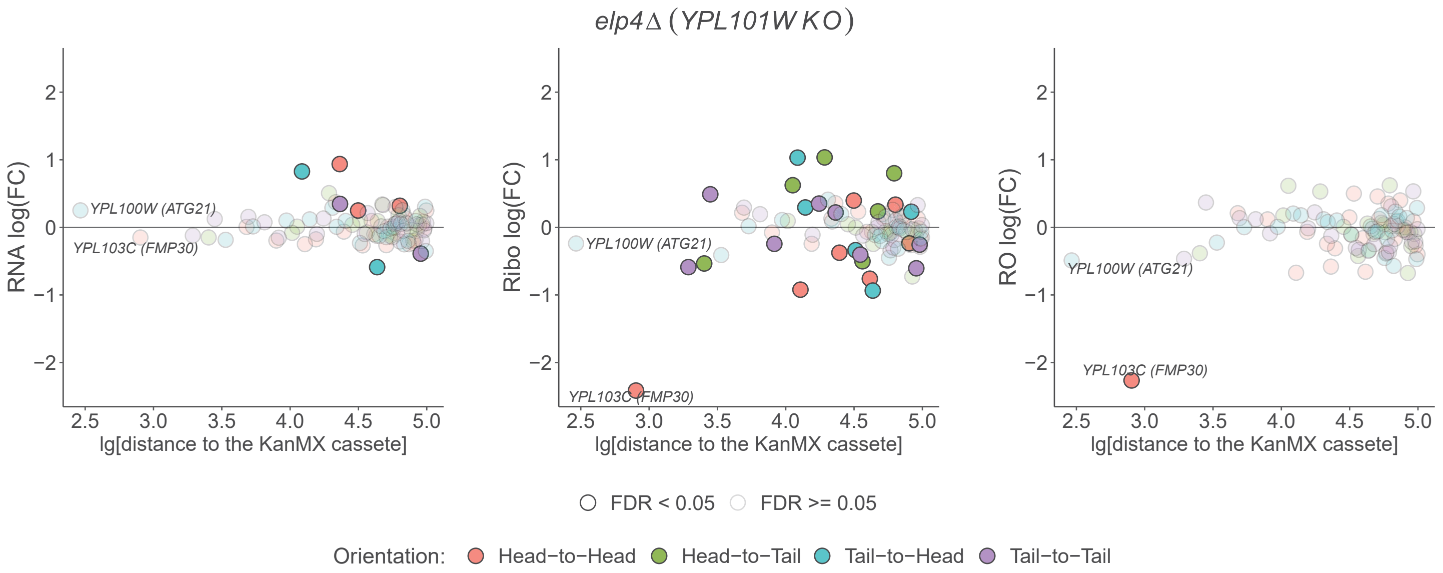
elp4Δ (YPL101W KO)
RNA log(FC)
Ribo log(FC)
RO log(FC)
YPL100W (ATG21)
YPL103C (FMP30)
lg[distance to the KanMX cassete]
FDR < 0.05
FDR >= 0.05
Orientation:
Head−to−Head
Head−to−Tail
Tail−to−Head
Tail−to−Tail

**B**

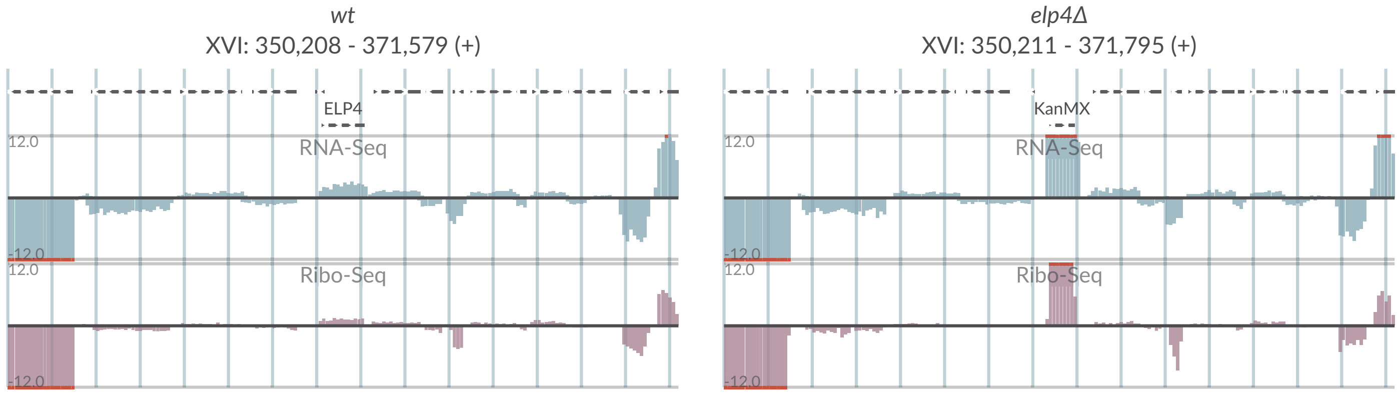
wt
XVI: 350,208 - 371,579 (+)
ELP4
RNA-Seq
Ribo-Seq
elp4Δ
XVI: 350,211 - 371,795 (+)
KanMX
RNA-Seq
Ribo-Seq

# S2-16

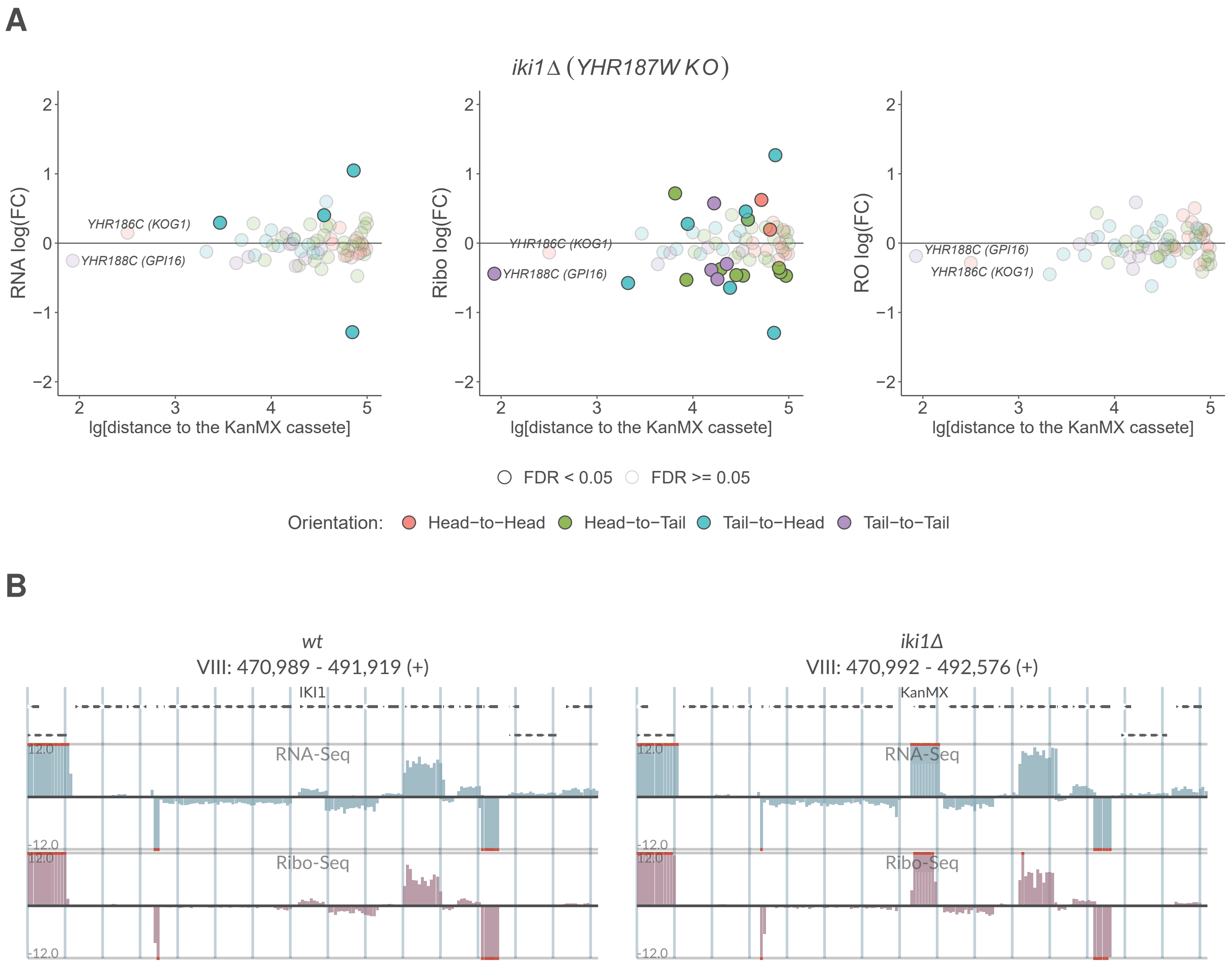
A
iki1Δ (YHR187W KO)
RNA log(FC)
Ribo log(FC)
RO log(FC)
YHR186C (KOG1)
YHR188C (GPI16)
lg[distance to the KanMX cassete]
FDR < 0.05
FDR >= 0.05
Orientation:
Head−to−Head
Head−to−Tail
Tail−to−Head
Tail−to−Tail
B
wt
VIII: 470,989 - 491,919 (+)
IKI1
iki1Δ
VIII: 470,992 - 492,576 (+)
KanMX
RNA-Seq
Ribo-Seq
12.0
-12.0

# S2-17

## A

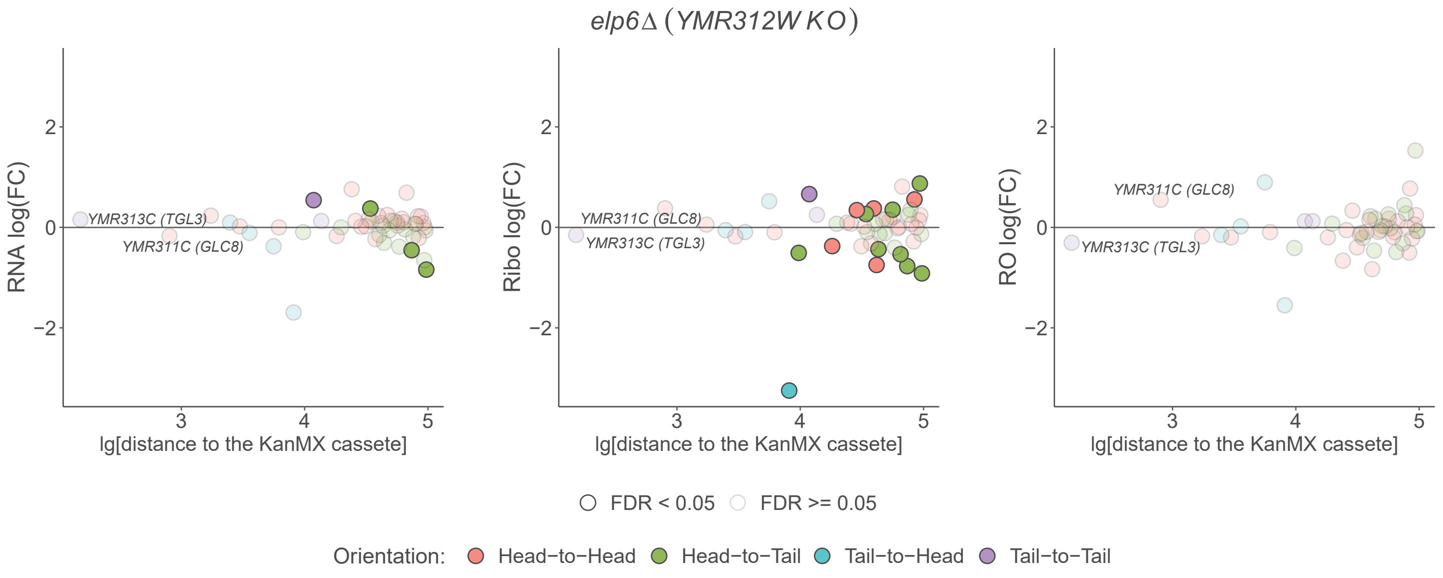

elp6Δ (YMR312W KO)
RNA log(FC)
Ribo log(FC)
RO log(FC)
YMR313C (TGL3)
YMR311C (GLC8)
lg[distance to the KanMX cassete]
FDR < 0.05
FDR >= 0.05
Orientation:
Head−to−Head
Head−to−Tail
Tail−to−Head
Tail−to−Tail


## B

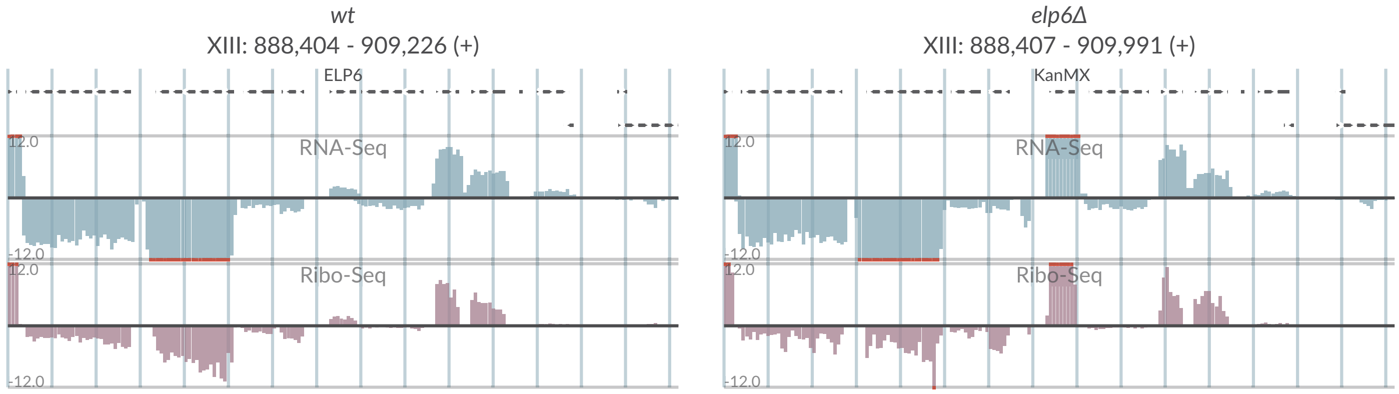

wt
XIII: 888,404 - 909,226 (+)
ELP6
elp6Δ
XIII: 888,407 - 909,991 (+)
KanMX
RNA-Seq
Ribo-Seq
12.0
-12.0

# S2-18

## A

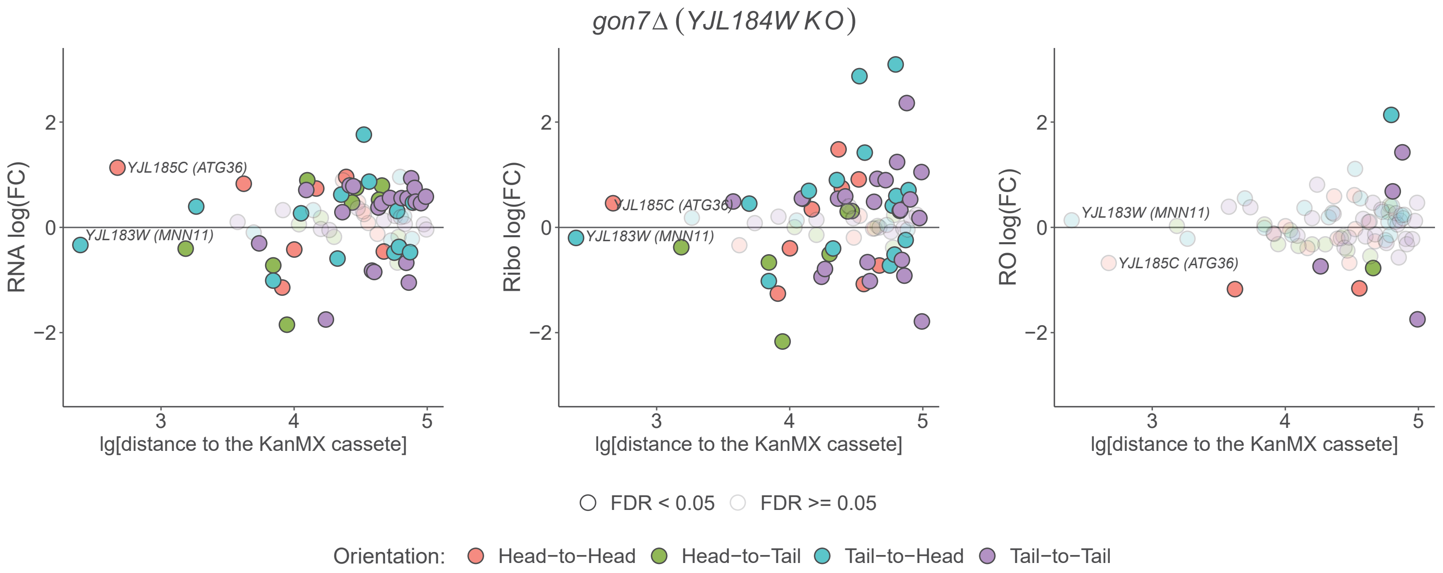

gon7Δ (YJL184W KO)
RNA log(FC)
Ribo log(FC)
RO log(FC)
YJL185C (ATG36)
YJL183W (MNN11)
lg[distance to the KanMX cassete]
FDR < 0.05
FDR >= 0.05
Orientation:
Head−to−Head
Head−to−Tail
Tail−to−Head
Tail−to−Tail


## B

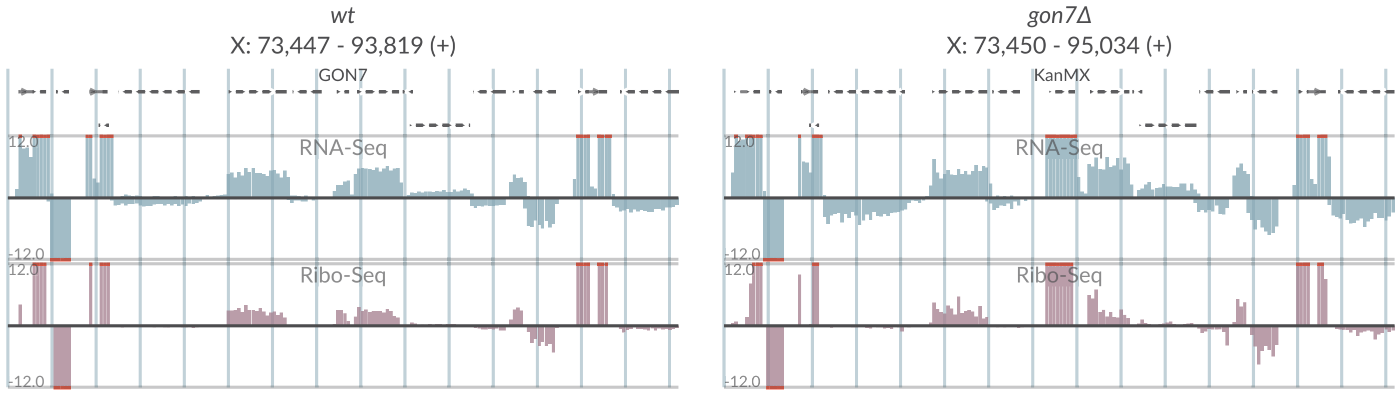

wt
X: 73,447 - 93,819 (+)
GON7
RNA-Seq
Ribo-Seq
gon7Δ
X: 73,450 - 95,034 (+)
KanMX
RNA-Seq
Ribo-Seq

## S2-19

**A**

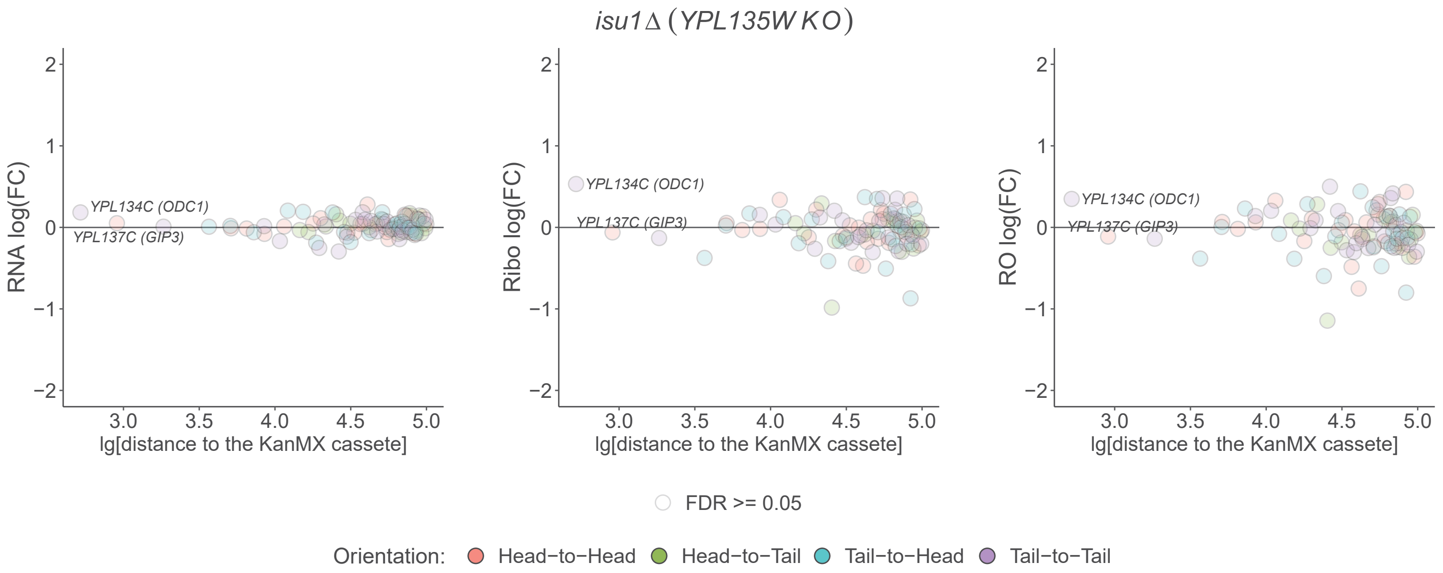

isu1Δ (YPL135W KO)
RNA log(FC)
Ribo log(FC)
RO log(FC)
YPL134C (ODC1)
YPL137C (GIP3)
lg[distance to the KanMX cassete]
FDR >= 0.05
Orientation: Head−to−Head Head−to−Tail Tail−to−Head Tail−to−Tail


**B**

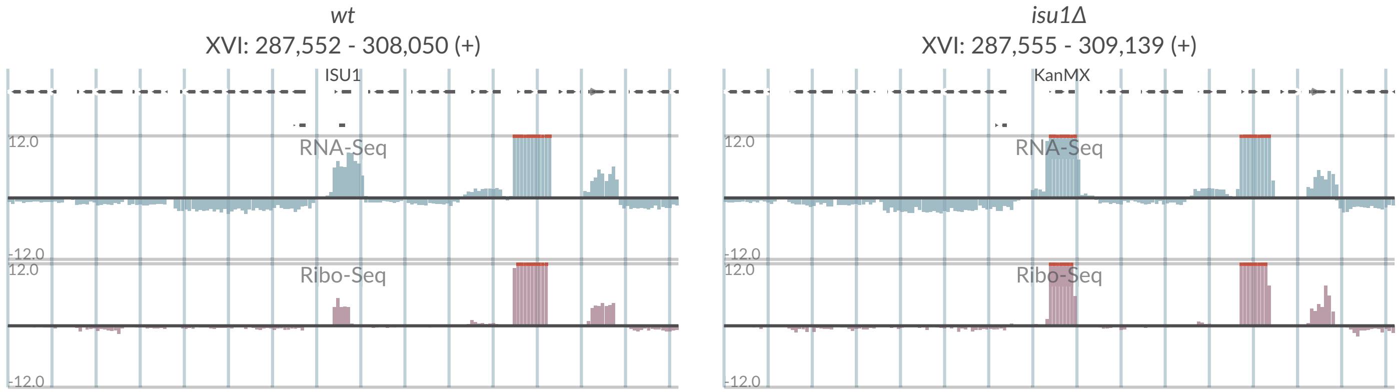

wt
XVI: 287,552 - 308,050 (+)
ISU1
isu1Δ
XVI: 287,555 - 309,139 (+)
KanMX
RNA-Seq
Ribo-Seq
12.0
-12.0

# S2-20

## A

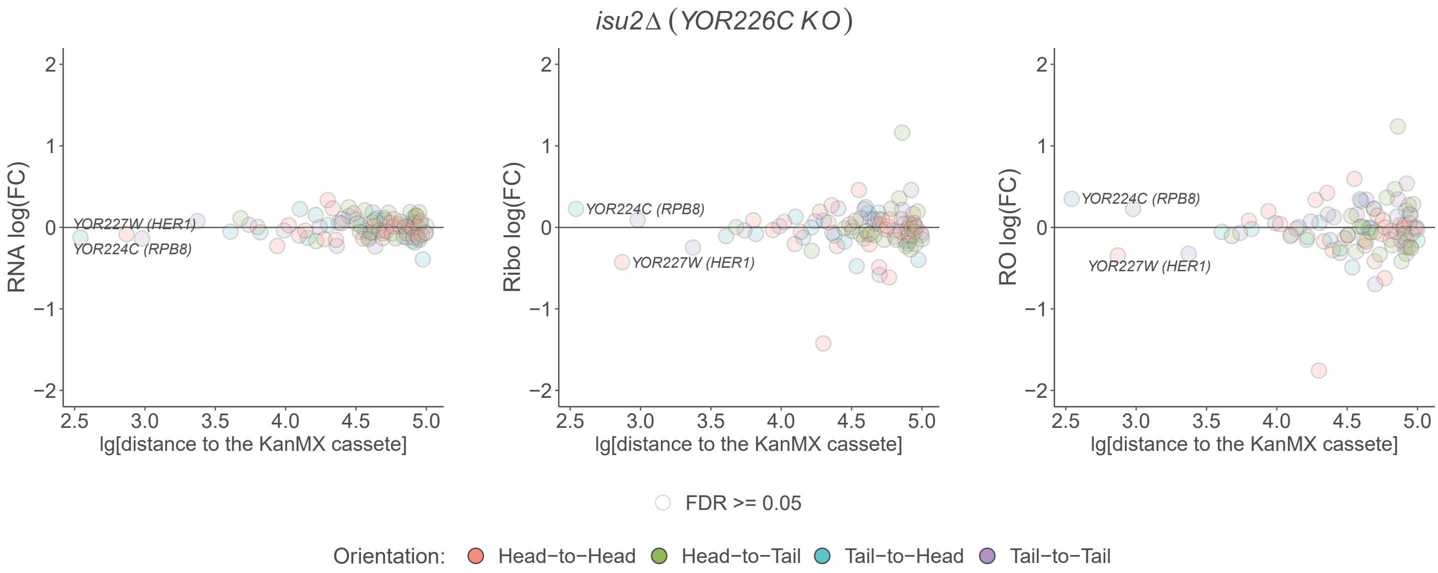

isu2Δ (YOR226C KO)
RNA log(FC)
Ribo log(FC)
RO log(FC)
YOR227W (HER1)
YOR224C (RPB8)
lg[distance to the KanMX cassete]
FDR >= 0.05
Orientation: Head−to−Head Head−to−Tail Tail−to−Head Tail−to−Tail


## B

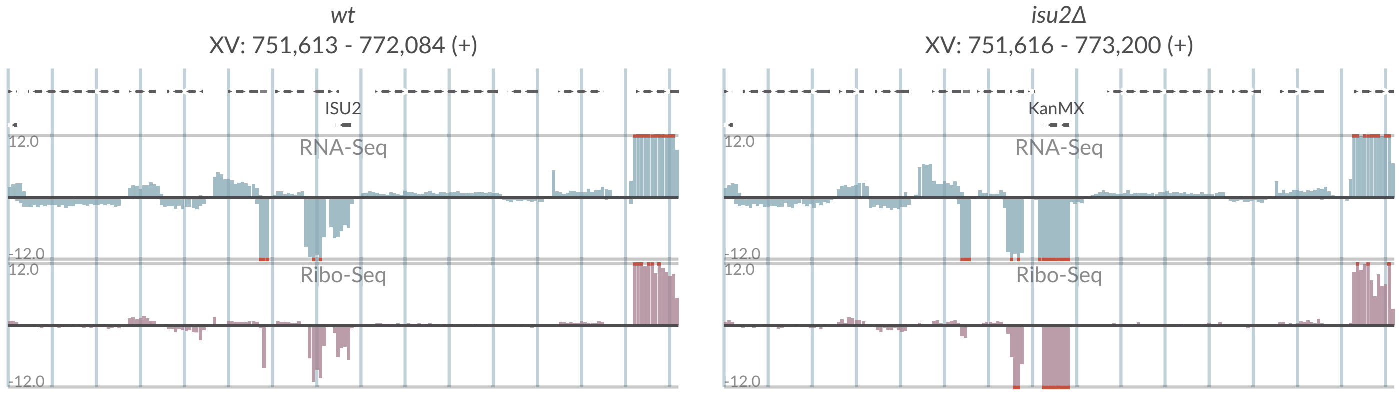

wt
XV: 751,613 - 772,084 (+)
ISU2
RNA-Seq
Ribo-Seq
isu2Δ
XV: 751,616 - 773,200 (+)
KanMX

# S2-21

**A**

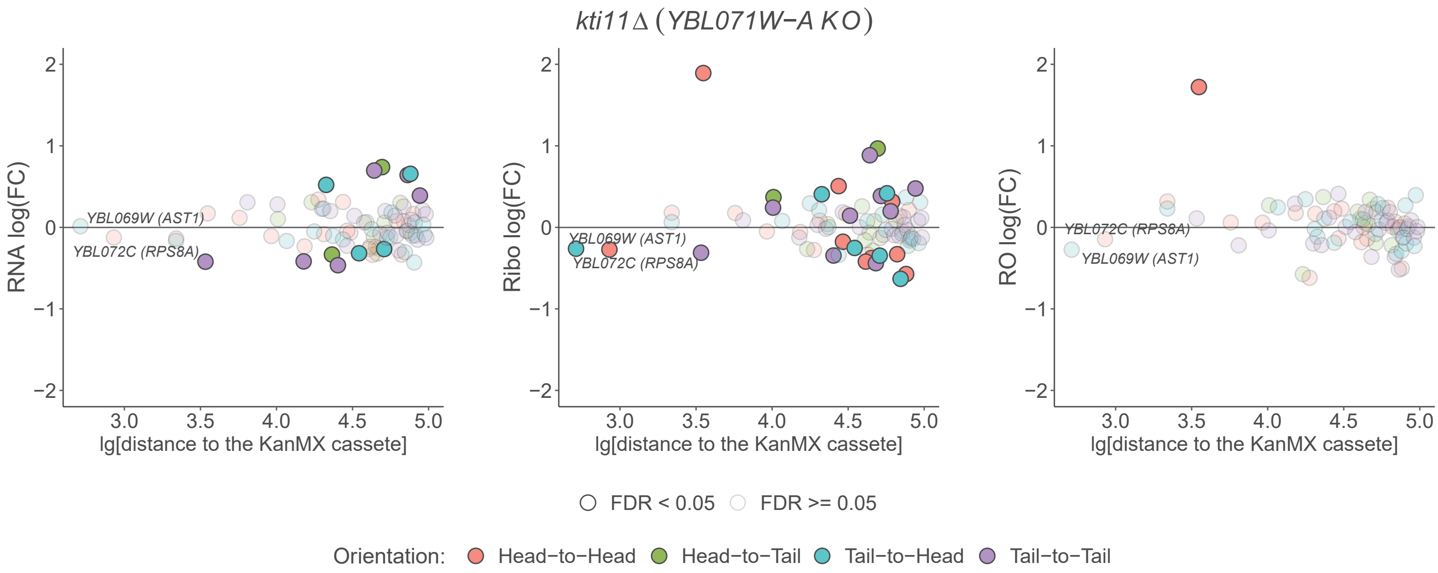

kti11Δ (YBL071W−A KO)
RNA log(FC)
Ribo log(FC)
RO log(FC)
YBL069W (AST1)
YBL072C (RPS8A)
lg[distance to the KanMX cassete]
FDR < 0.05
FDR >= 0.05
Orientation:
Head−to−Head
Head−to−Tail
Tail−to−Head
Tail−to−Tail


**B**

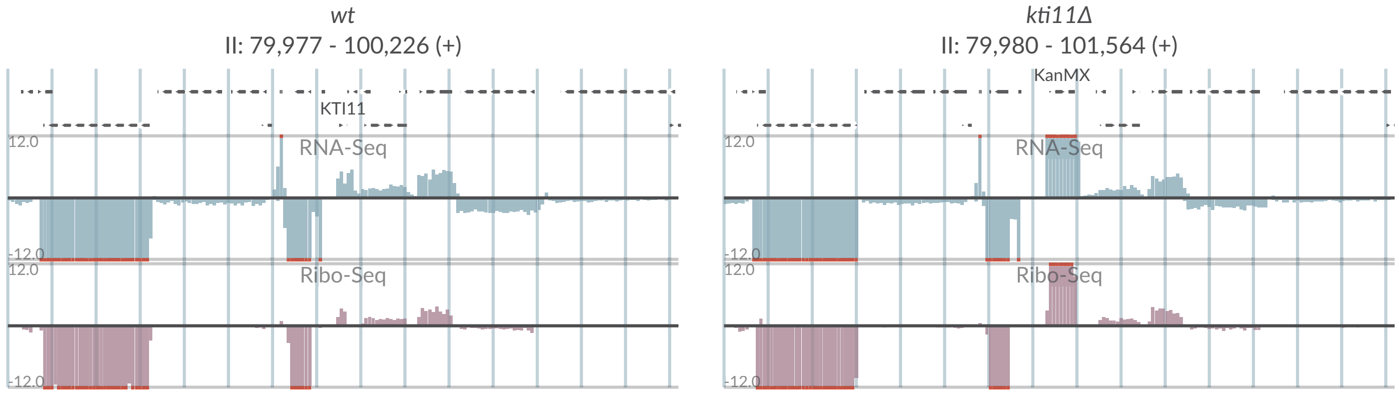

wt
II: 79,977 - 100,226 (+)
kti11Δ
II: 79,980 - 101,564 (+)
KTI11
KanMX
RNA-Seq
Ribo-Seq
12.0
-12.0

# S2-22

A

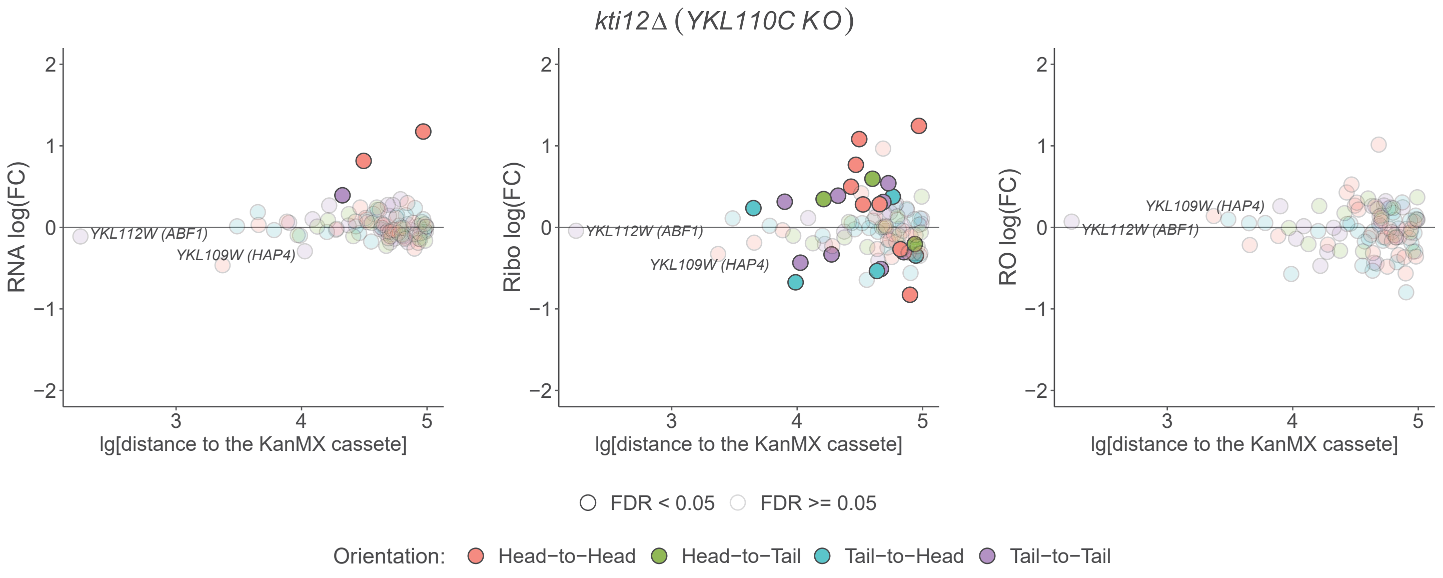
kti12Δ (YKL110C KO)
RNA log(FC)
Ribo log(FC)
RO log(FC)
YKL112W (ABF1)
YKL109W (HAP4)
lg[distance to the KanMX cassete]
FDR < 0.05
FDR >= 0.05
Orientation:
Head−to−Head
Head−to−Tail
Tail−to−Head
Tail−to−Tail

B

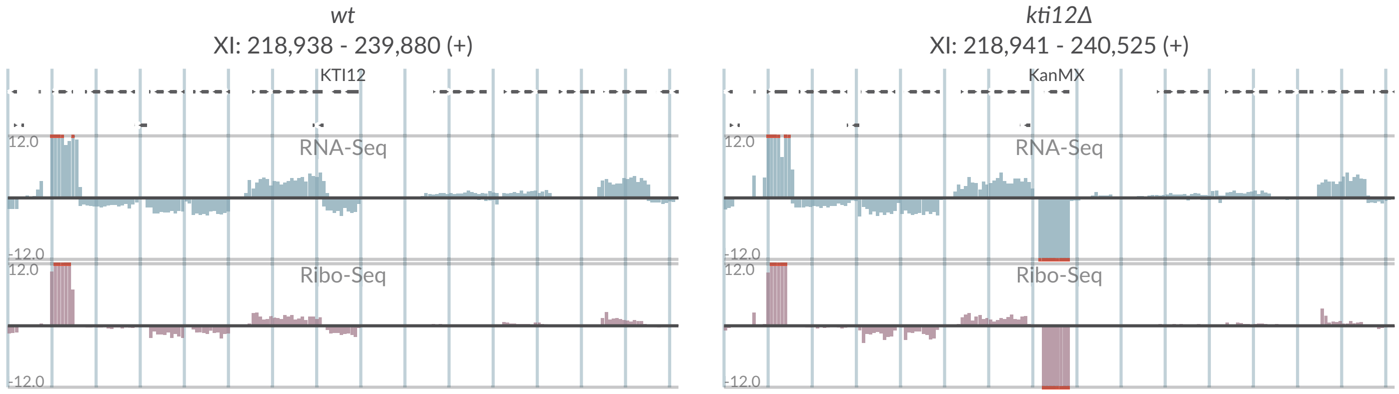
wt
XI: 218,938 - 239,880 (+)
KTI12
kti12Δ
XI: 218,941 - 240,525 (+)
KanMX
RNA-Seq
Ribo-Seq
12.0
-12.0

# S2-23

**A**

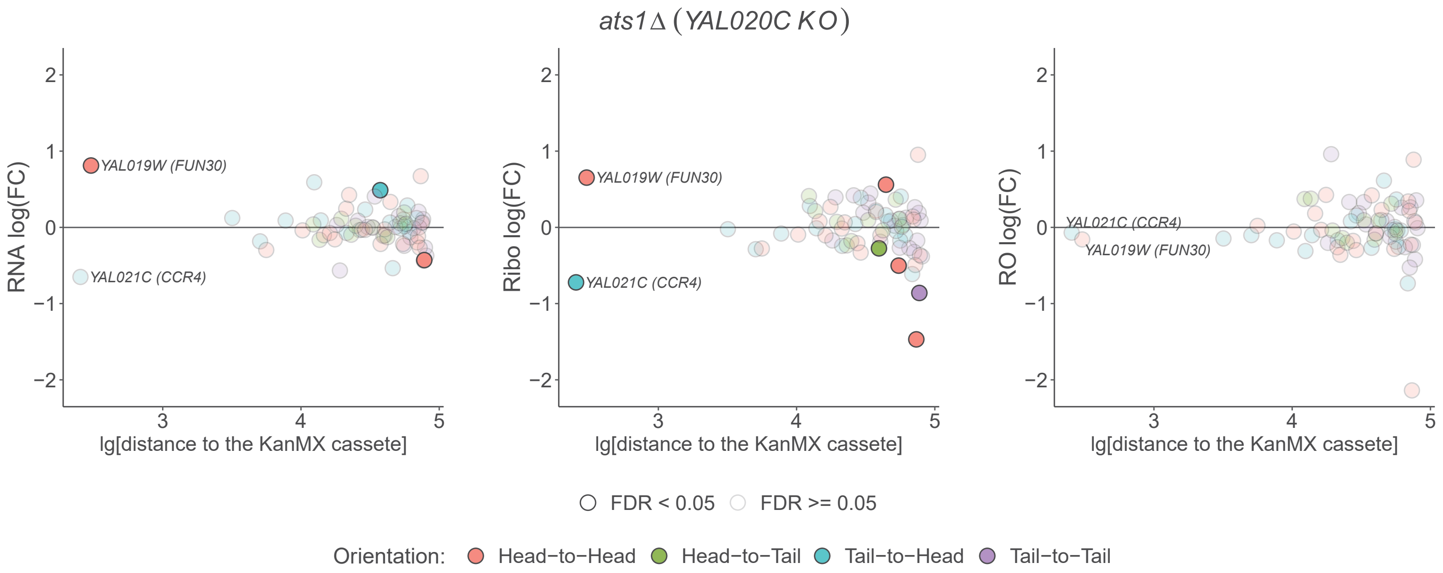
ats1Δ (YAL020C KO)
RNA log(FC)
Ribo log(FC)
RO log(FC)
YAL019W (FUN30)
YAL021C (CCR4)
lg[distance to the KanMX cassete]
FDR < 0.05
FDR >= 0.05
Orientation:
Head−to−Head
Head−to−Tail
Tail−to−Head
Tail−to−Tail

**B**

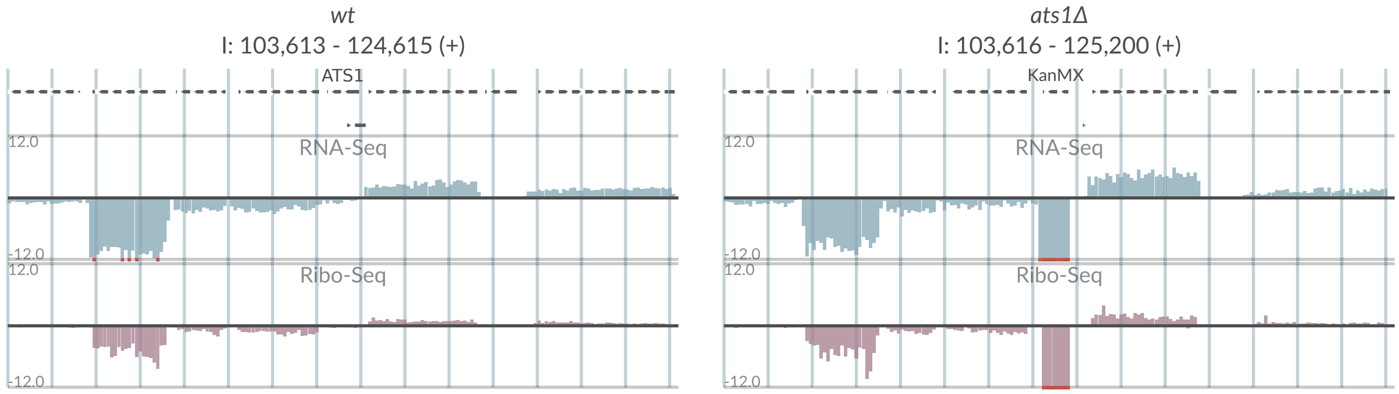
wt
I: 103,613 - 124,615 (+)
ATS1
ats1Δ
I: 103,616 - 125,200 (+)
KanMX
RNA-Seq
Ribo-Seq
12.0
-12.0

# S2-24

## A

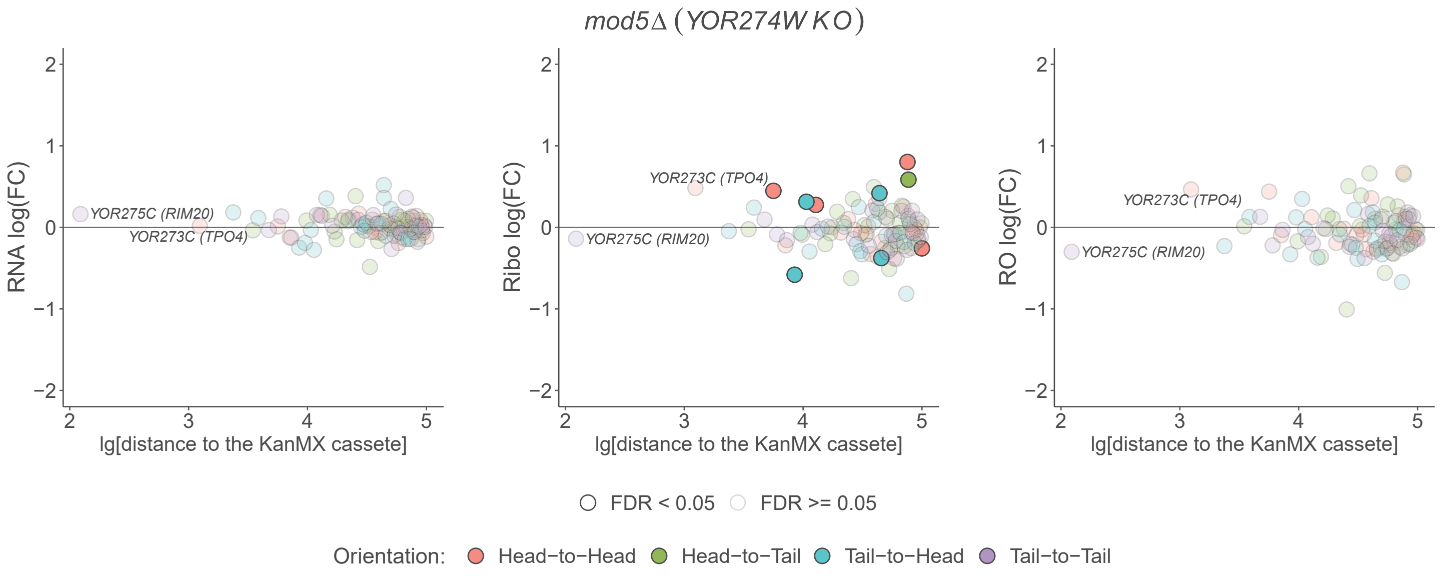

mod5Δ (YOR274W KO)
RNA log(FC)
Ribo log(FC)
RO log(FC)
YOR275C (RIM20)
YOR273C (TPO4)
lg[distance to the KanMX cassete]
FDR < 0.05
FDR >= 0.05
Orientation:
Head−to−Head
Head−to−Tail
Tail−to−Head
Tail−to−Tail


## B

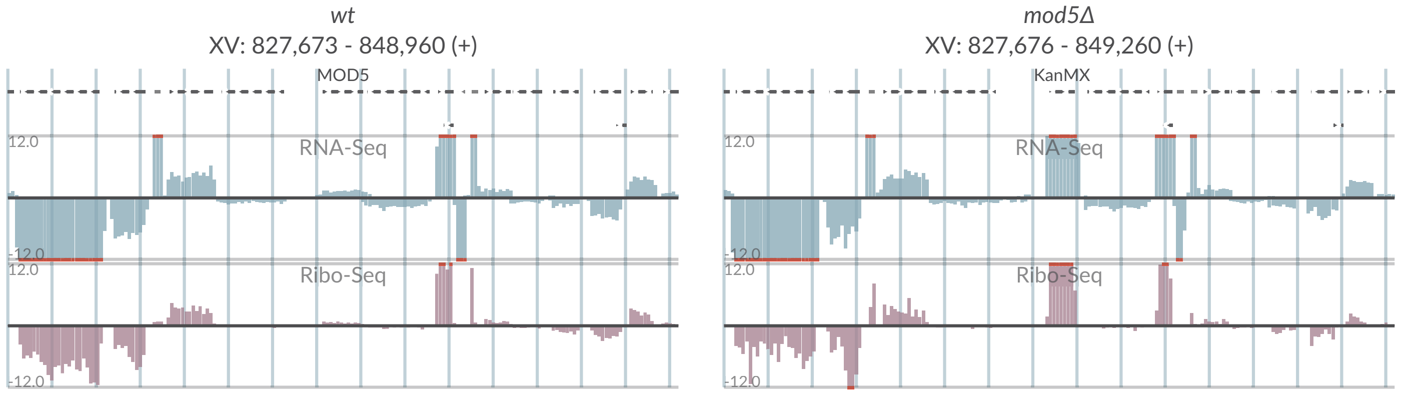

wt
XV: 827,673 - 848,960 (+)
MOD5
RNA-Seq
Ribo-Seq
12.0
-12.0
mod5Δ
XV: 827,676 - 849,260 (+)
KanMX

# S2-25

## A

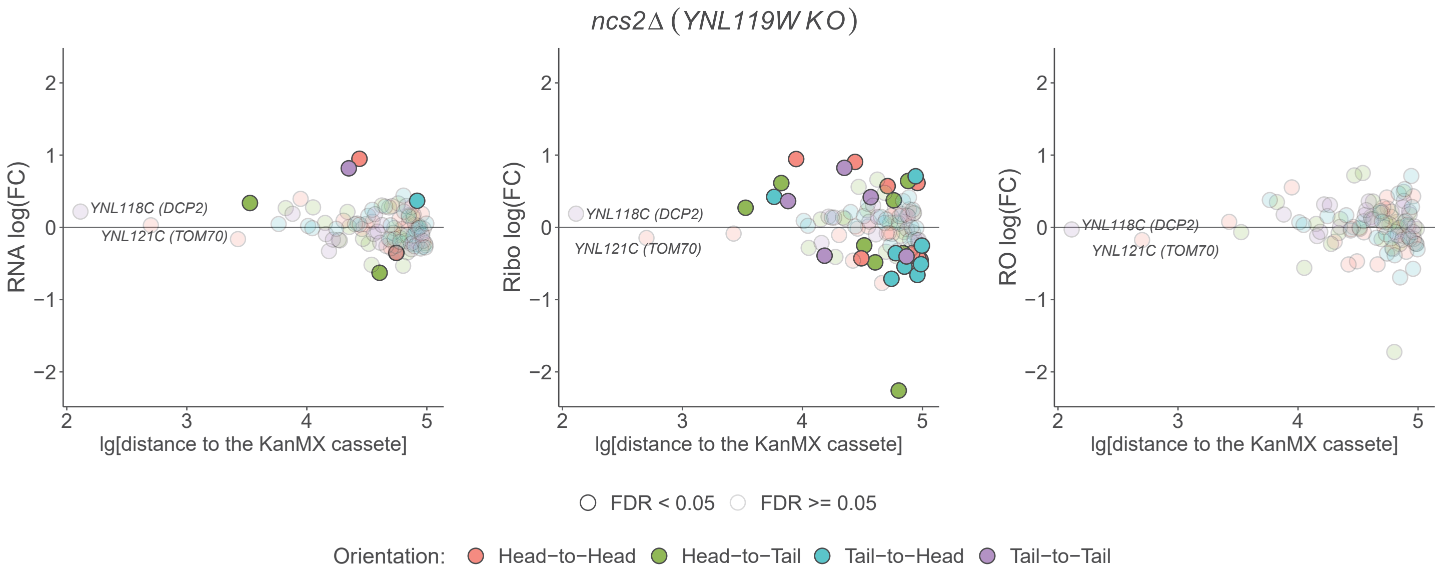

ncs2Δ (YNL119W KO)
RNA log(FC)
Ribo log(FC)
RO log(FC)
YNL118C (DCP2)
YNL121C (TOM70)
lg[distance to the KanMX cassete]
FDR < 0.05
FDR >= 0.05
Orientation:
Head−to−Head
Head−to−Tail
Tail−to−Head
Tail−to−Tail


## B

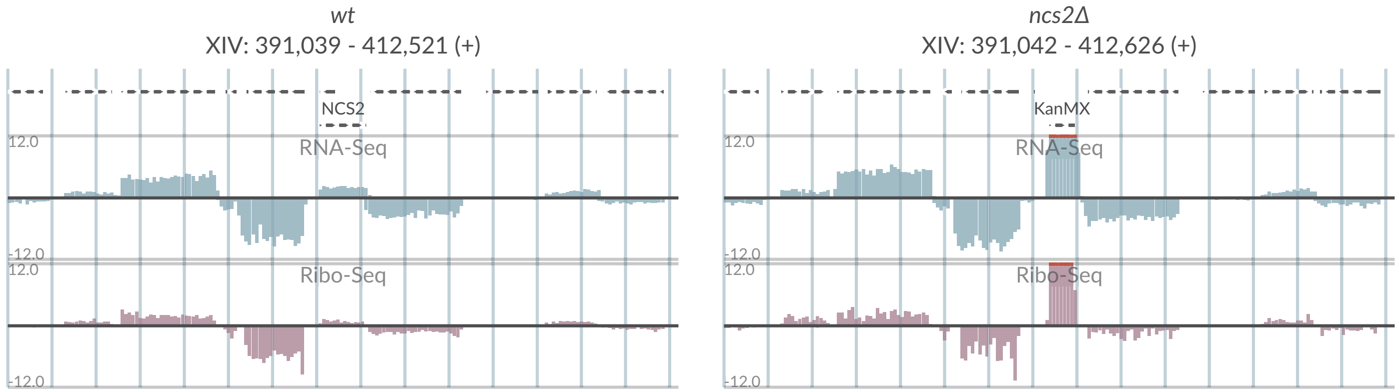

wt
XIV: 391,039 - 412,521 (+)
ncs2Δ
XIV: 391,042 - 412,626 (+)
NCS2
KanMX
RNA-Seq
Ribo-Seq
12.0
-12.0

# S2-26

**A**

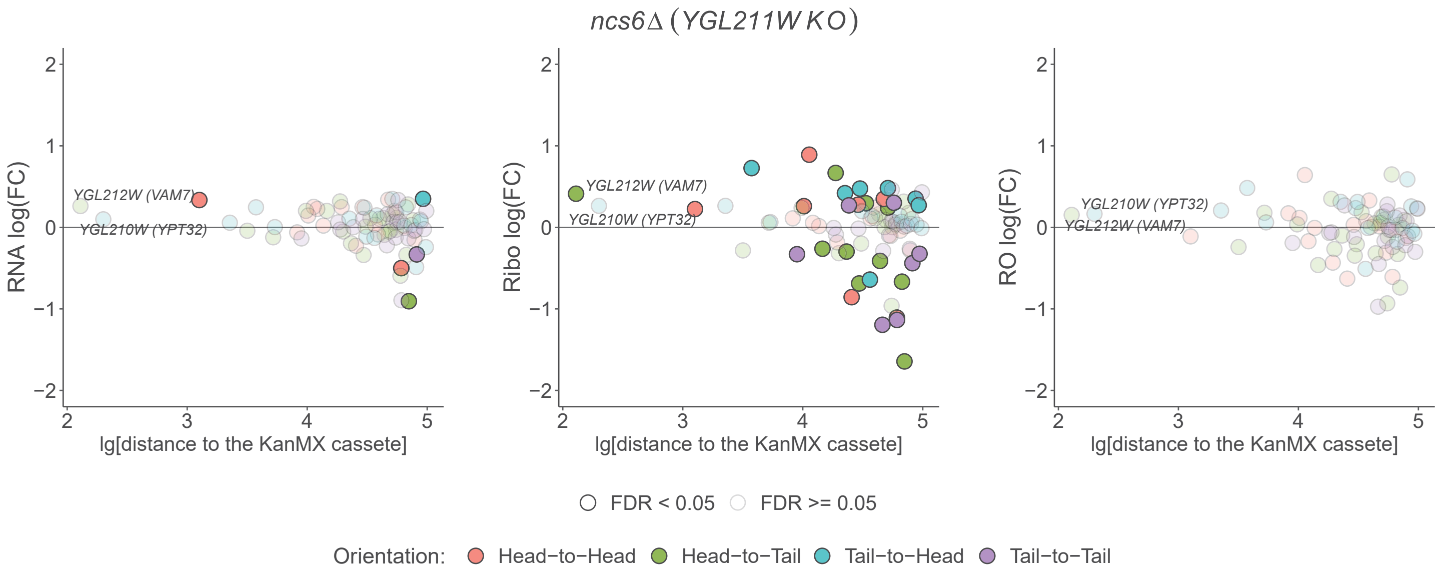

ncs6Δ (YGL211W KO)
RNA log(FC)
Ribo log(FC)
RO log(FC)
lg[distance to the KanMX cassete]
YGL212W (VAM7)
YGL210W (YPT32)
FDR < 0.05
FDR >= 0.05
Orientation:
Head−to−Head
Head−to−Tail
Tail−to−Head
Tail−to−Tail


**B**

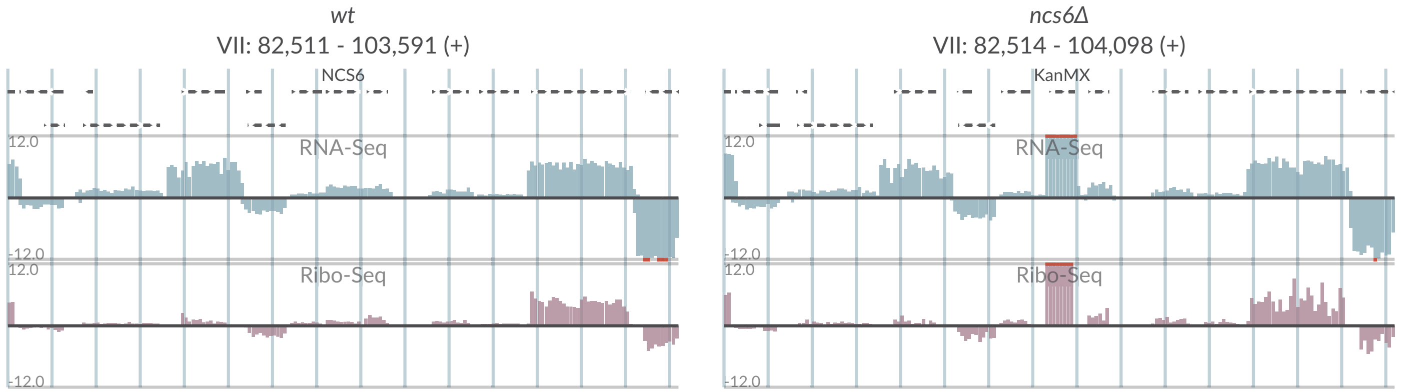

wt
VII: 82,511 - 103,591 (+)
ncs6Δ
VII: 82,514 - 104,098 (+)
NCS6
KanMX
RNA-Seq
Ribo-Seq
12.0
-12.0

# S2-27

## A

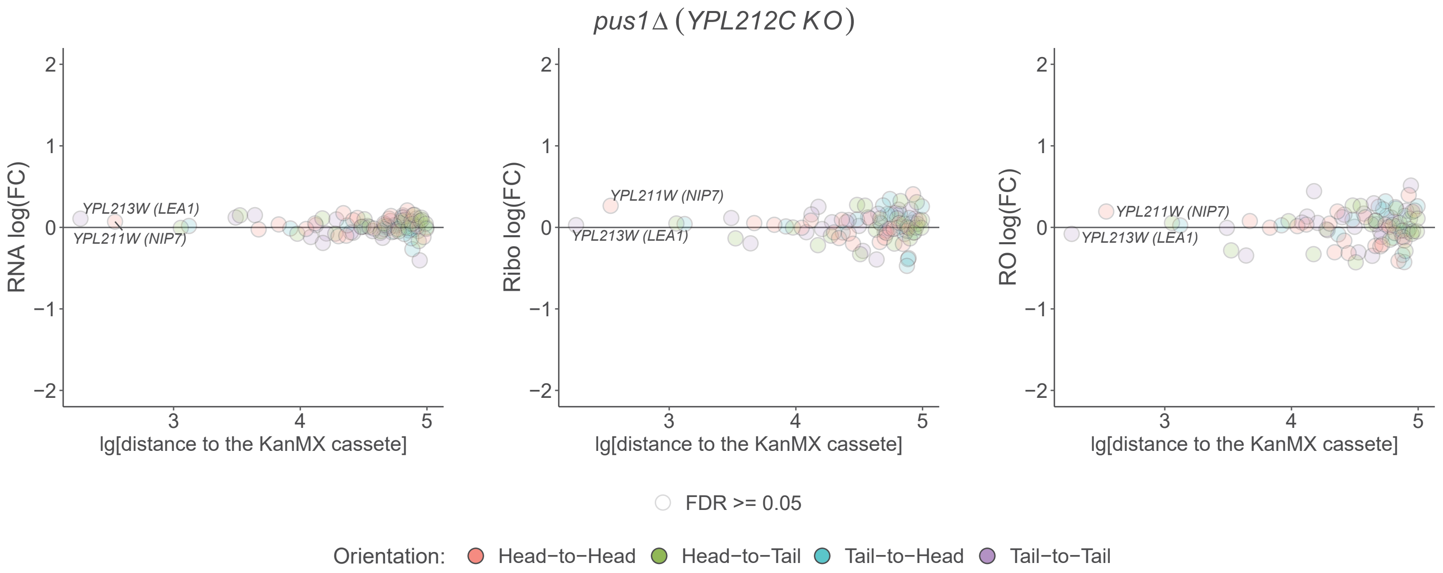

pus1Δ (YPL212C KO)
RNA log(FC)
Ribo log(FC)
RO log(FC)
YPL213W (LEA1)
YPL211W (NIP7)
lg[distance to the KanMX cassete]
FDR >= 0.05
Orientation: Head−to−Head Head−to−Tail Tail−to−Head Tail−to−Tail


## B

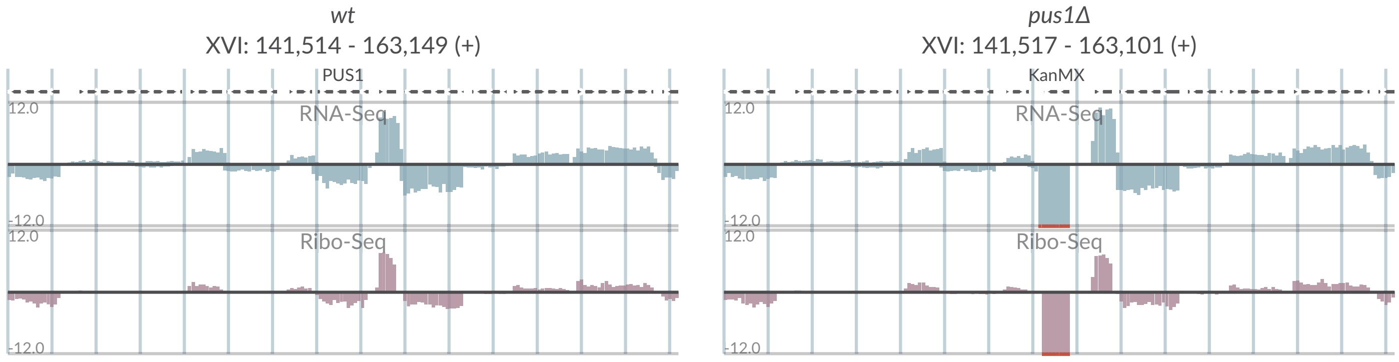

wt
XVI: 141,514 - 163,149 (+)
PUS1
pus1Δ
XVI: 141,517 - 163,101 (+)
KanMX
RNA-Seq
Ribo-Seq
12.0
-12.0

# S2-28

## A

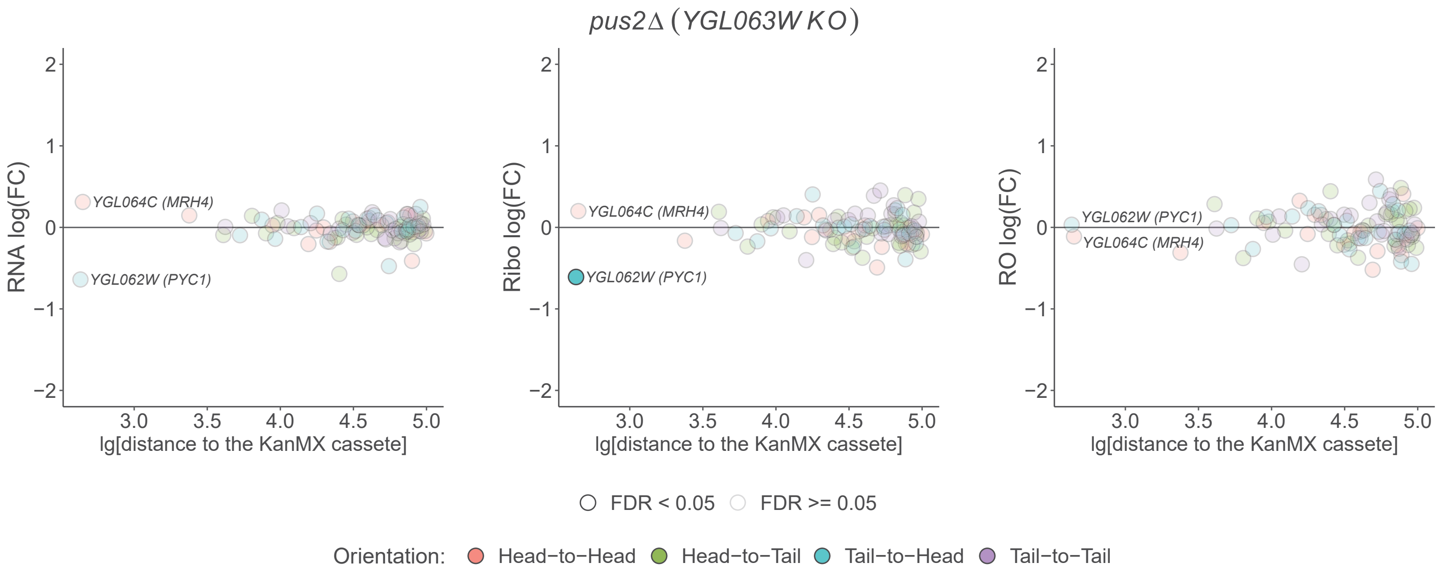

pus2Δ (YGL063W KO)
RNA log(FC)
Ribo log(FC)
RO log(FC)
YGL064C (MRH4)
YGL062W (PYC1)
lg[distance to the KanMX cassete]
FDR < 0.05
FDR >= 0.05
Orientation:
Head−to−Head
Head−to−Tail
Tail−to−Head
Tail−to−Tail


## B

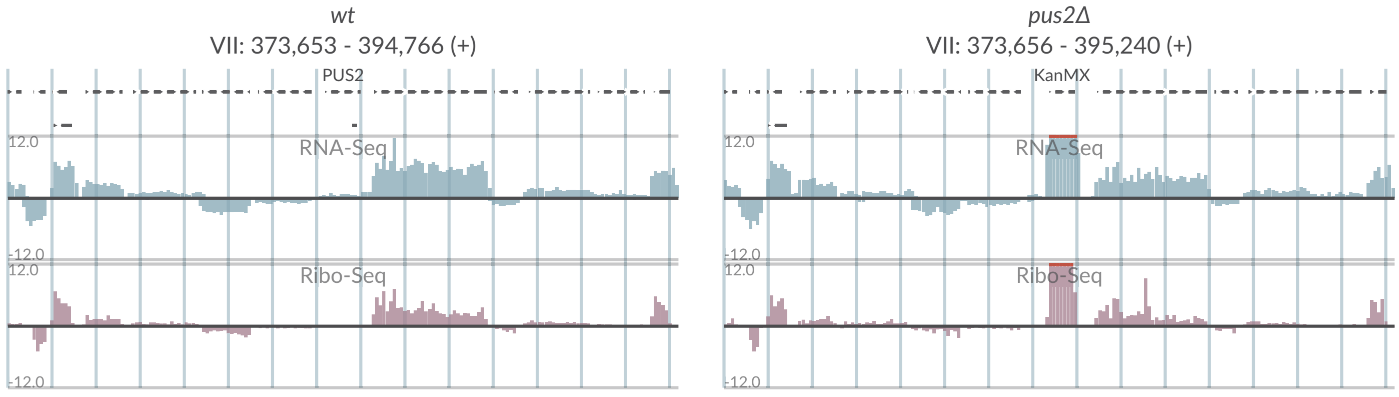

wt
VII: 373,653 - 394,766 (+)
pus2Δ
VII: 373,656 - 395,240 (+)
PUS2
KanMX
RNA-Seq
Ribo-Seq
12.0
-12.0

# S2-29

## A

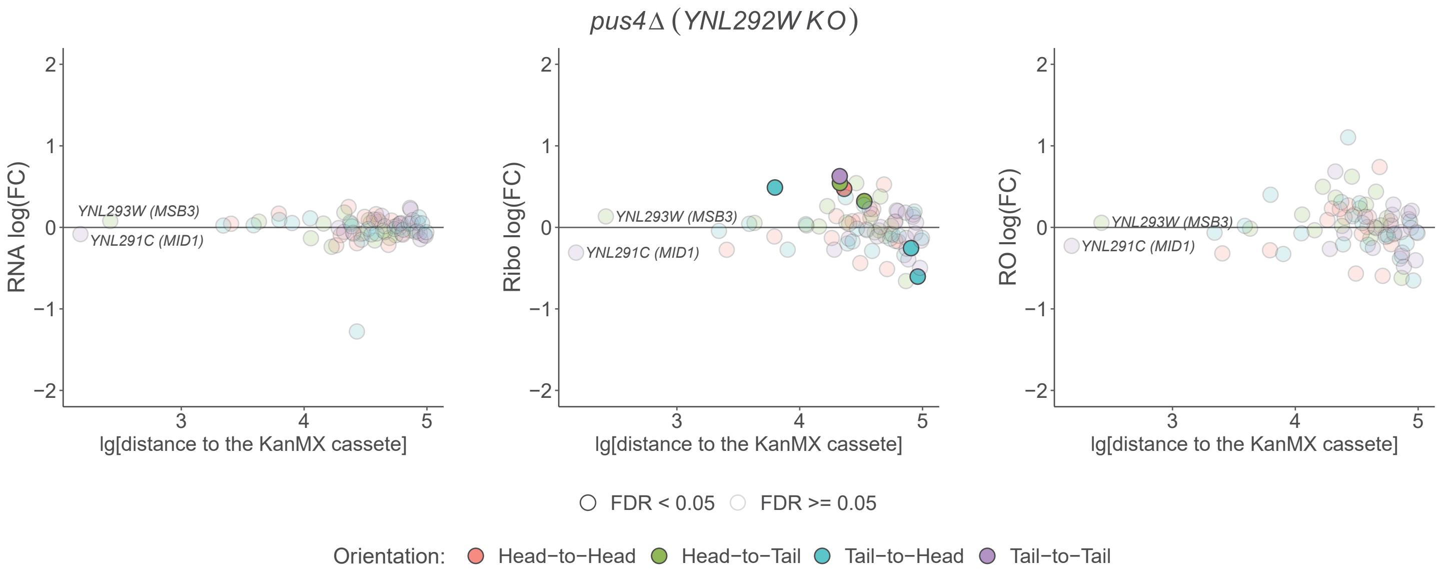

pus4Δ (YNL292W KO)
RNA log(FC)
Ribo log(FC)
RO log(FC)
YNL293W (MSB3)
YNL291C (MID1)
lg[distance to the KanMX cassete]
FDR < 0.05
FDR >= 0.05
Orientation:
Head−to−Head
Head−to−Tail
Tail−to−Head
Tail−to−Tail


## B

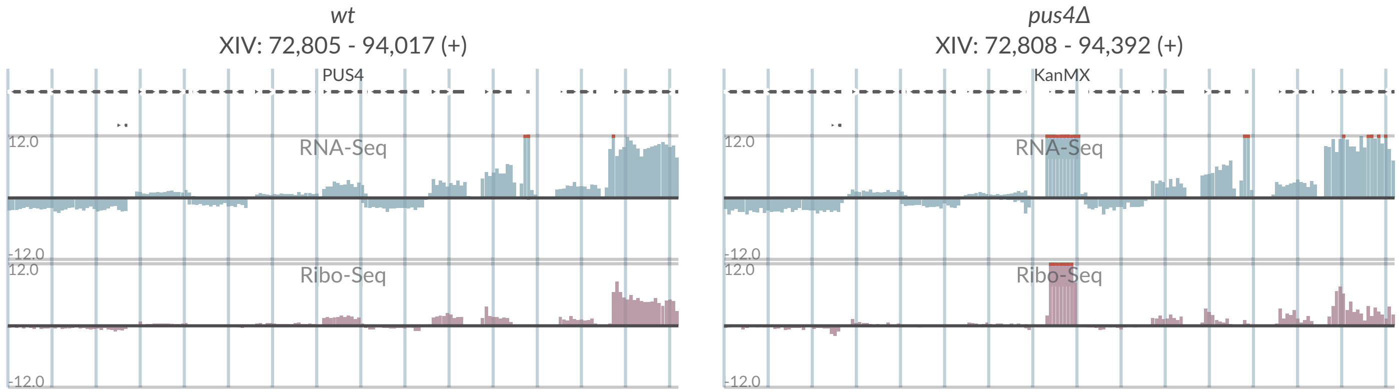

wt
XIV: 72,805 - 94,017 (+)
PUS4
RNA-Seq
Ribo-Seq
pus4Δ
XIV: 72,808 - 94,392 (+)
KanMX

# S2-30

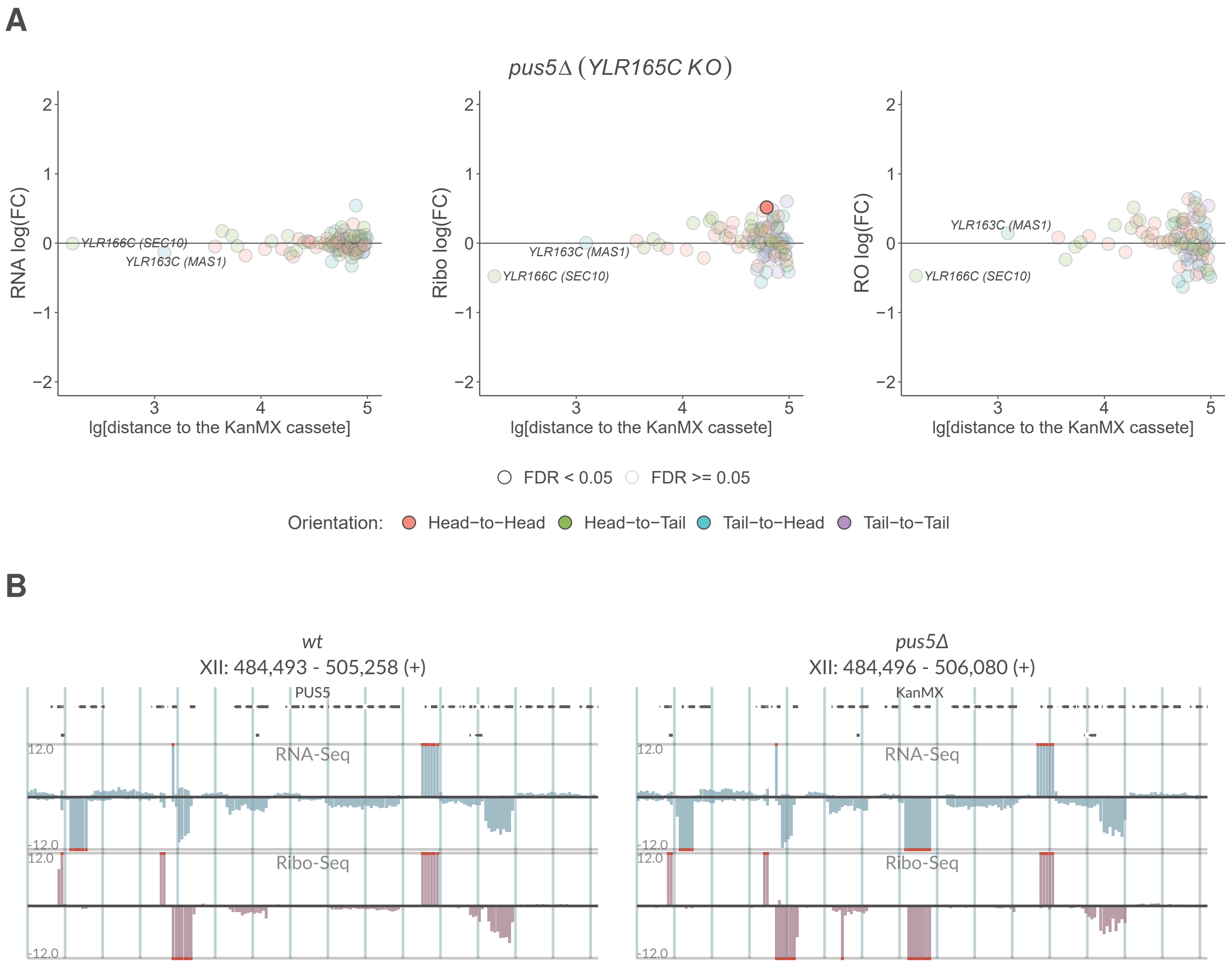
A
pus5Δ (YLR165C KO)
RNA log(FC)
Ribo log(FC)
RO log(FC)
lg[distance to the KanMX cassete]
YLR166C (SEC10)
YLR163C (MAS1)
FDR < 0.05
FDR >= 0.05
Orientation:
Head−to−Head
Head−to−Tail
Tail−to−Head
Tail−to−Tail
B
wt
XII: 484,493 - 505,258 (+)
PUS5
pus5Δ
XII: 484,496 - 506,080 (+)
KanMX
RNA-Seq
Ribo-Seq
12.0
-12.0

# S2-31

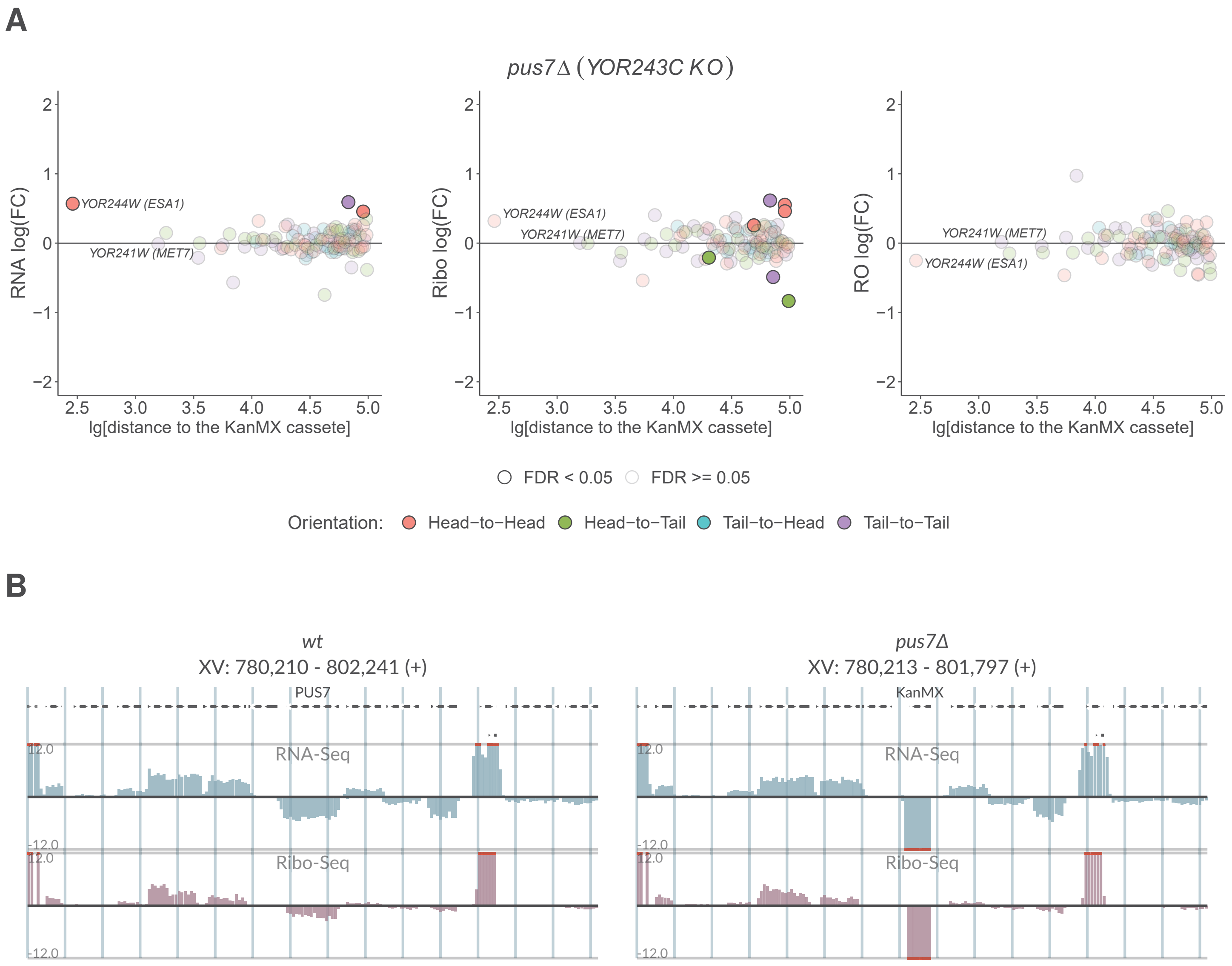
A
pus7Δ (YOR243C KO)
RNA log(FC)
Ribo log(FC)
RO log(FC)
lg[distance to the KanMX cassete]
YOR244W (ESA1)
YOR241W (MET7)
FDR < 0.05
FDR >= 0.05
Orientation:
Head−to−Head
Head−to−Tail
Tail−to−Head
Tail−to−Tail
B
wt
XV: 780,210 - 802,241 (+)
PUS7
pus7Δ
XV: 780,213 - 801,797 (+)
KanMX
RNA-Seq
Ribo-Seq
12.0
-12.0

# S2-32

## A

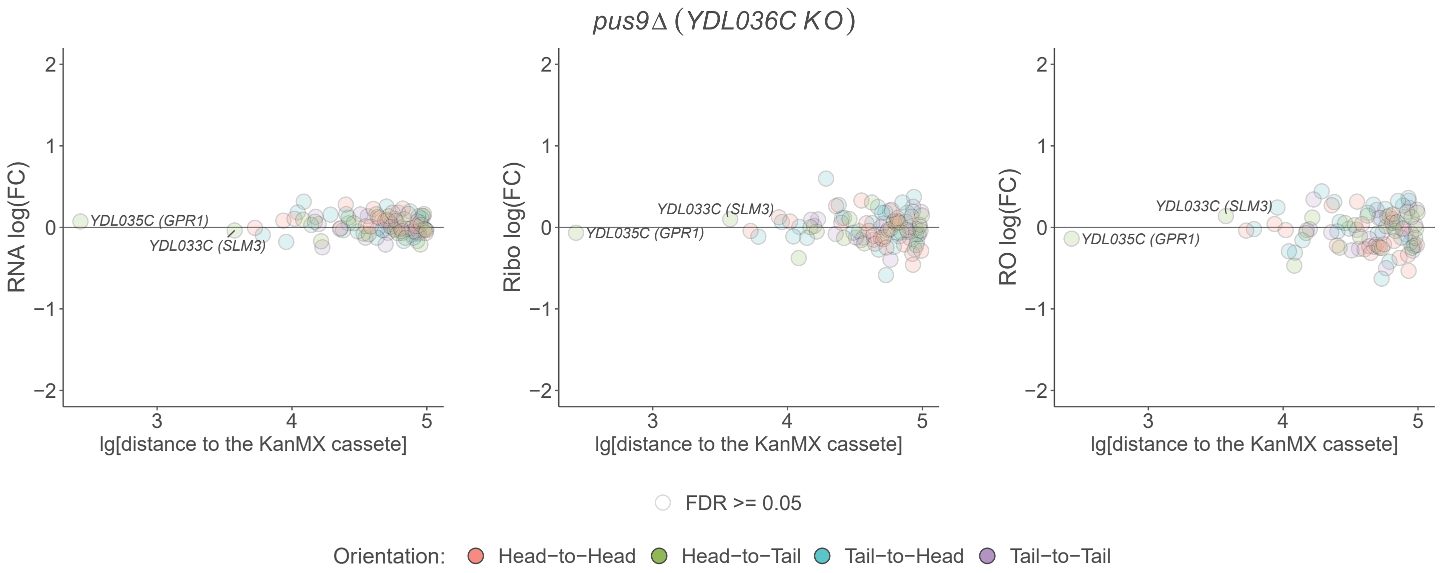

pus9Δ (YDL036C KO)
RNA log(FC)
Ribo log(FC)
RO log(FC)
lg[distance to the KanMX cassete]
YDL035C (GPR1)
YDL033C (SLM3)
FDR >= 0.05
Orientation: Head−to−Head Head−to−Tail Tail−to−Head Tail−to−Tail


## B

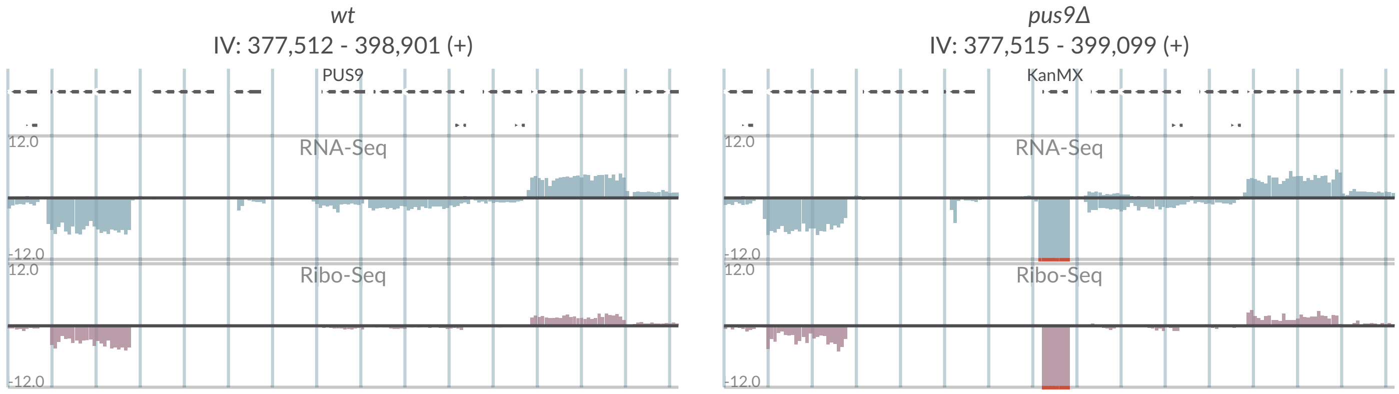

wt
IV: 377,512 - 398,901 (+)
PUS9
pus9Δ
IV: 377,515 - 399,099 (+)
KanMX
RNA-Seq
Ribo-Seq
12.0
-12.0

# S2-33

## A

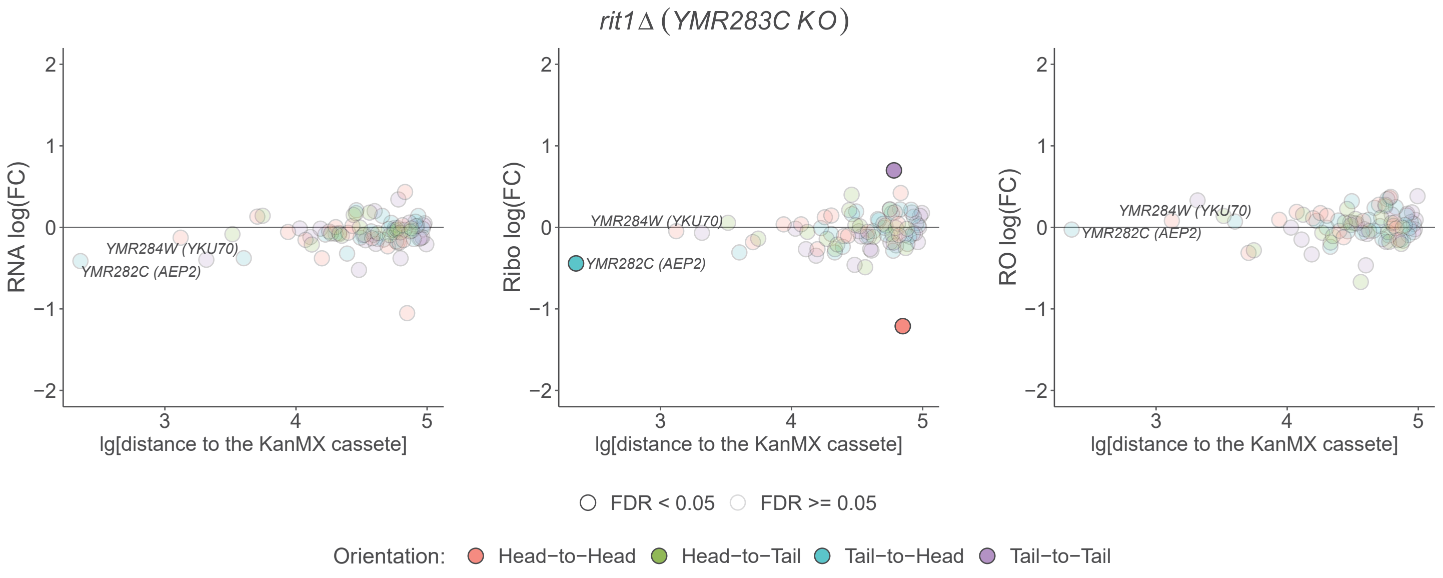

rit1Δ (YMR283C KO)
RNA log(FC)
Ribo log(FC)
RO log(FC)
YMR284W (YKU70)
YMR282C (AEP2)
lg[distance to the KanMX cassete]
FDR < 0.05
FDR >= 0.05
Orientation:
Head−to−Head
Head−to−Tail
Tail−to−Head
Tail−to−Tail


## B

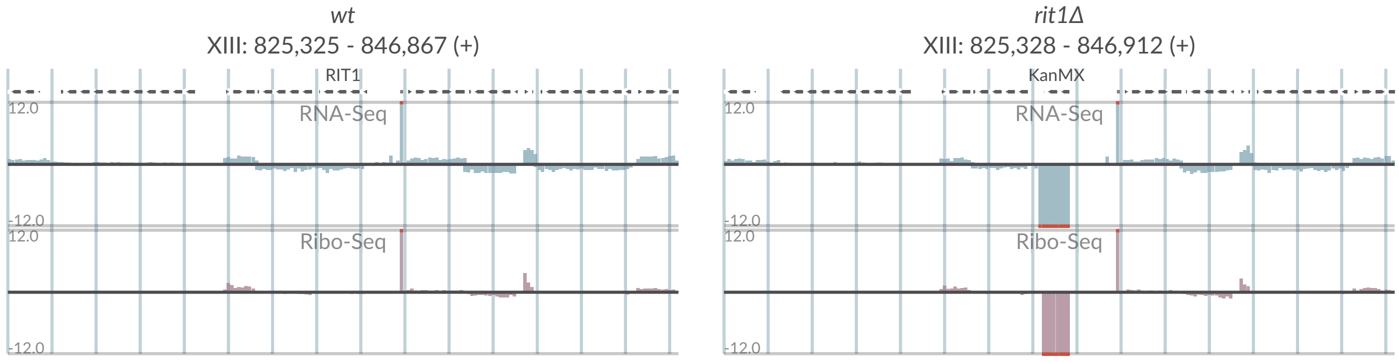

wt
XIII: 825,325 - 846,867 (+)
rit1Δ
XIII: 825,328 - 846,912 (+)
RIT1
KanMX
RNA-Seq
Ribo-Seq
12.0
-12.0

# S2-34

**A**

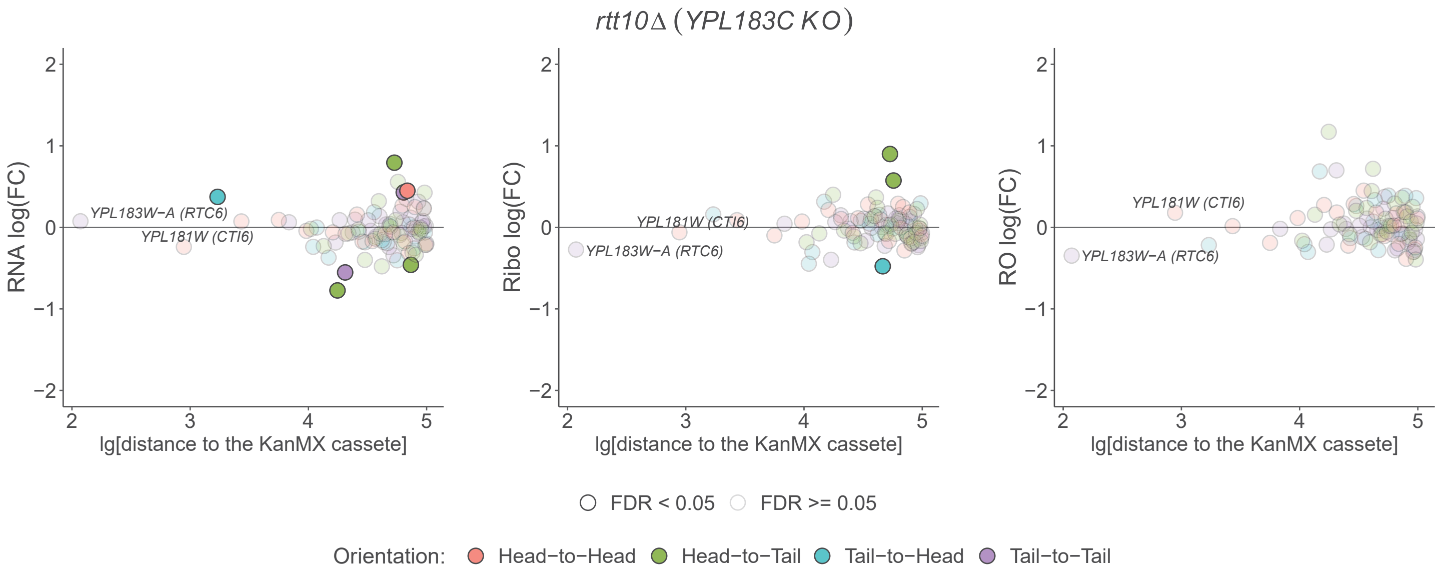

rtt10Δ (YPL183C KO)
RNA log(FC)
Ribo log(FC)
RO log(FC)
YPL183W−A (RTC6)
YPL181W (CTI6)
lg[distance to the KanMX cassete]
FDR < 0.05
FDR >= 0.05
Orientation:
Head−to−Head
Head−to−Tail
Tail−to−Head
Tail−to−Tail


**B**

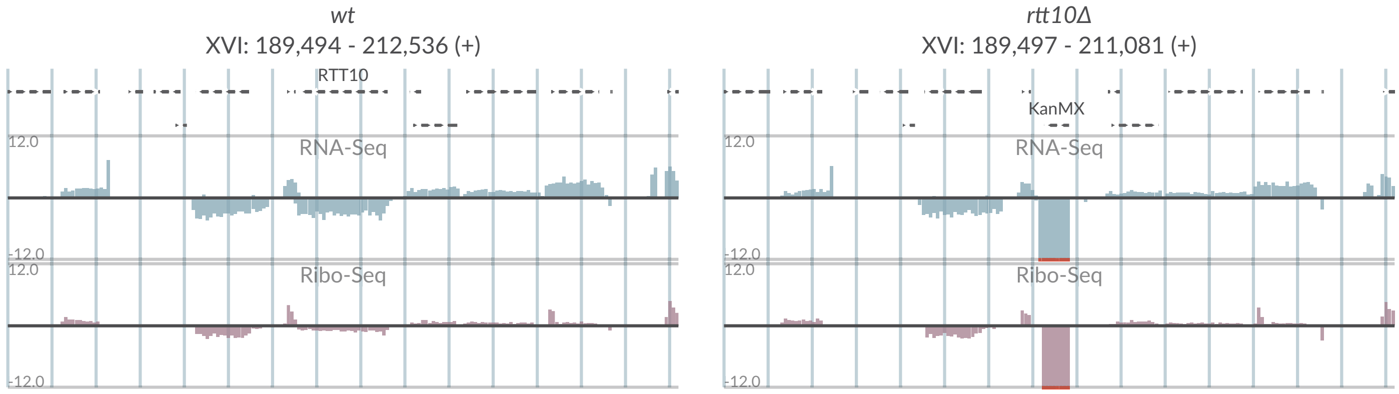

wt
XVI: 189,494 - 212,536 (+)
RTT10
RNA-Seq
Ribo-Seq
12.0
-12.0
rtt10Δ
XVI: 189,497 - 211,081 (+)
KanMX

# S2-35

## A

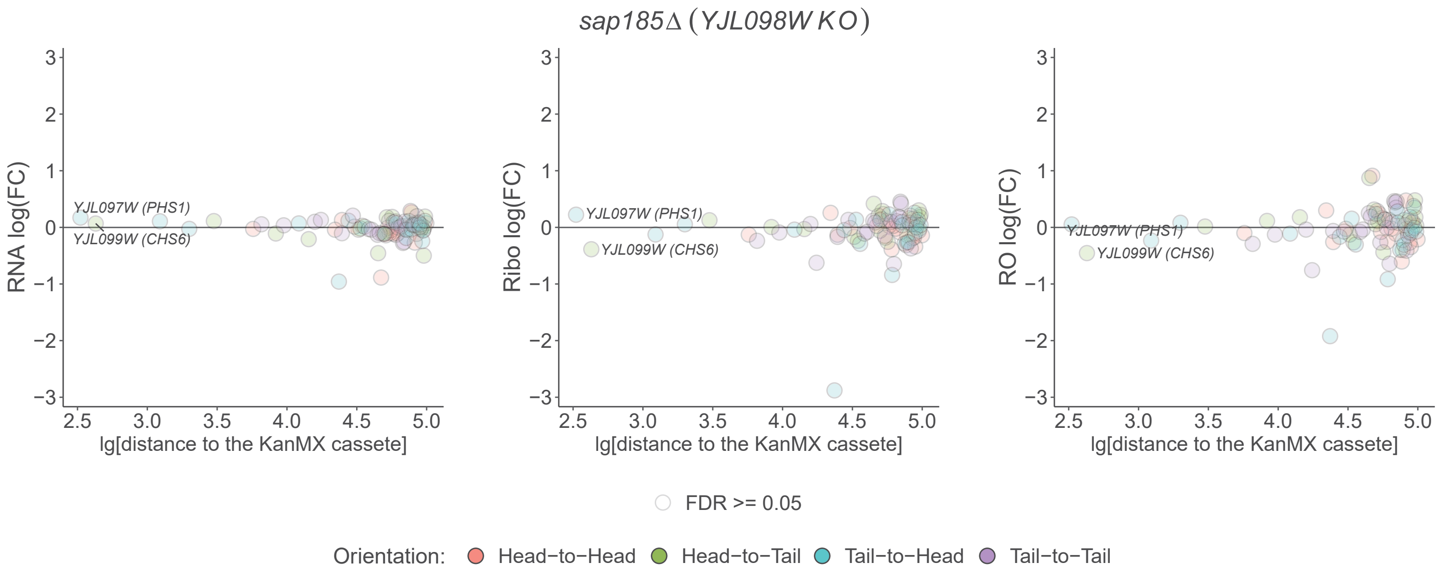

sap185Δ (YJL098W KO)
RNA log(FC)
Ribo log(FC)
RO log(FC)
YJL097W (PHS1)
YJL099W (CHS6)
lg[distance to the KanMX cassete]
FDR >= 0.05
Orientation: Head−to−Head Head−to−Tail Tail−to−Head Tail−to−Tail


## B

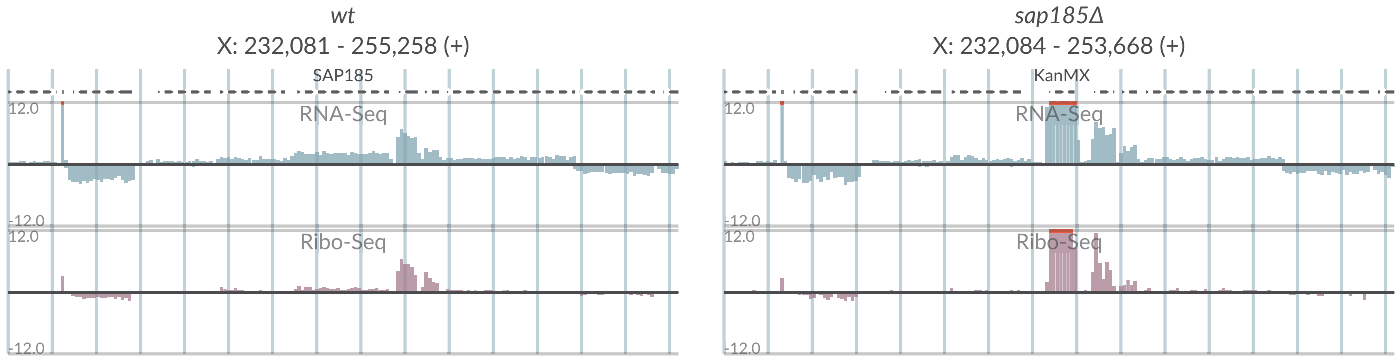

wt
X: 232,081 - 255,258 (+)
SAP185
RNA-Seq
Ribo-Seq
12.0
-12.0
sap185Δ
X: 232,084 - 253,668 (+)
KanMX

# S2-36

## A

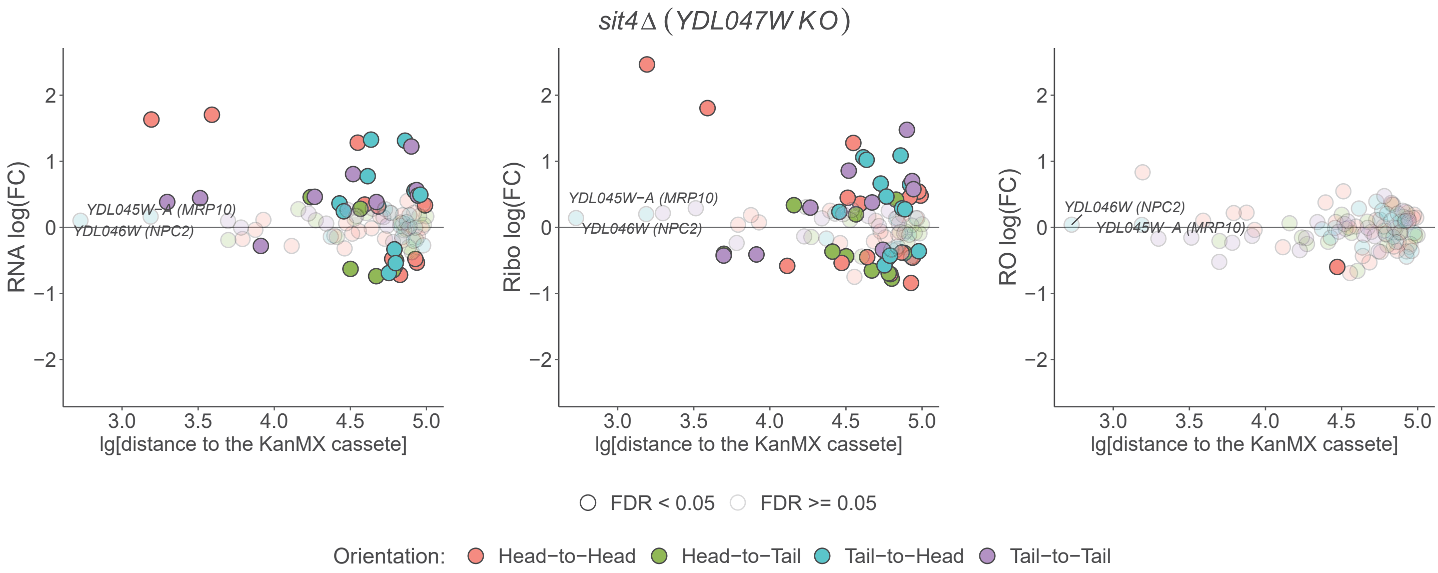

*sit4*Δ (*YDL047W KO*)
RNA log(FC)
Ribo log(FC)
RO log(FC)
lg[distance to the KanMX cassete]
YDL045W−A (MRP10)
YDL046W (NPC2)
FDR < 0.05
FDR >= 0.05
Orientation:
Head−to−Head
Head−to−Tail
Tail−to−Head
Tail−to−Tail


## B

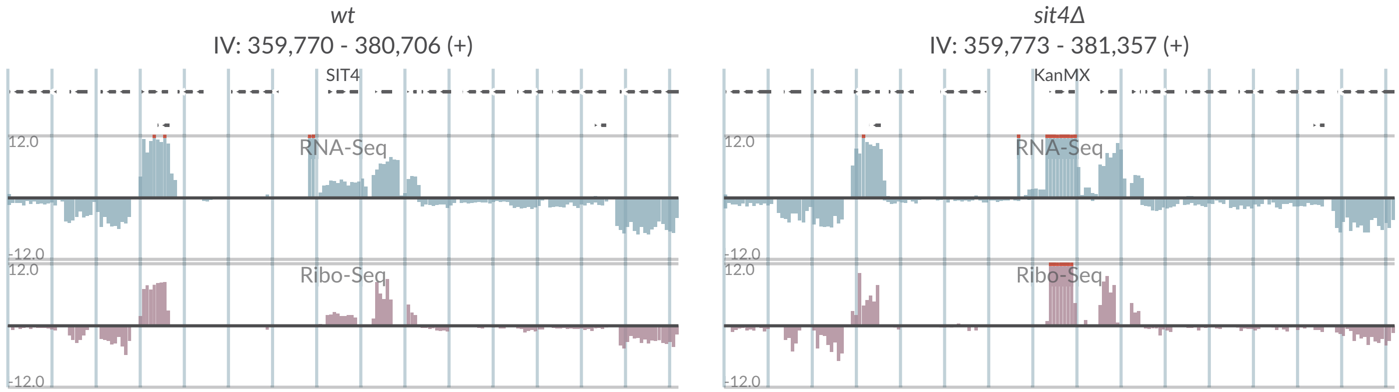

*wt*
IV: 359,770 - 380,706 (+)
SIT4
RNA-Seq
Ribo-Seq
*sit4Δ*
IV: 359,773 - 381,357 (+)
KanMX
RNA-Seq
Ribo-Seq

# S2-37

**A**

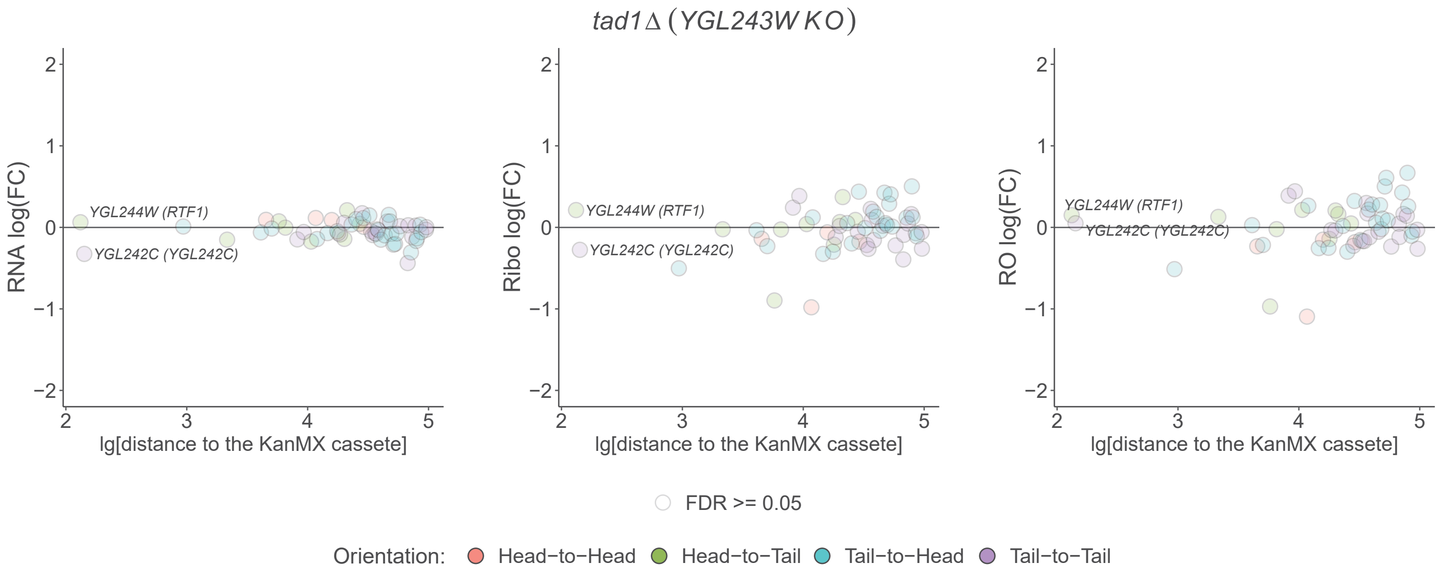

tad1Δ (YGL243W KO)
RNA log(FC)
Ribo log(FC)
RO log(FC)
YGL244W (RTF1)
YGL242C (YGL242C)
lg[distance to the KanMX cassete]
FDR >= 0.05
Orientation: Head−to−Head Head−to−Tail Tail−to−Head Tail−to−Tail


**B**

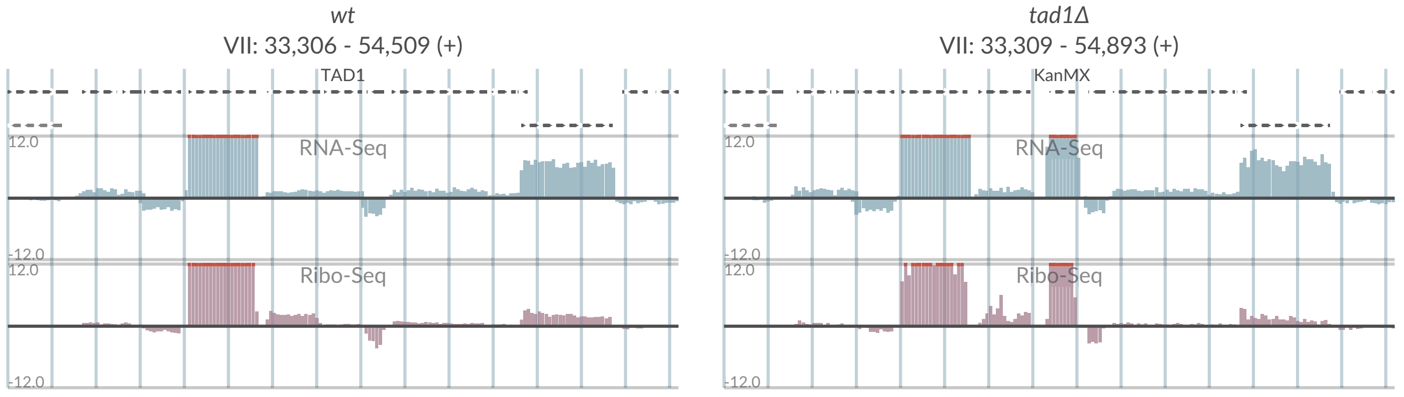

wt
VII: 33,306 - 54,509 (+)
TAD1
RNA-Seq
Ribo-Seq
tad1Δ
VII: 33,309 - 54,893 (+)
KanMX
RNA-Seq
Ribo-Seq

# S2-38

## A

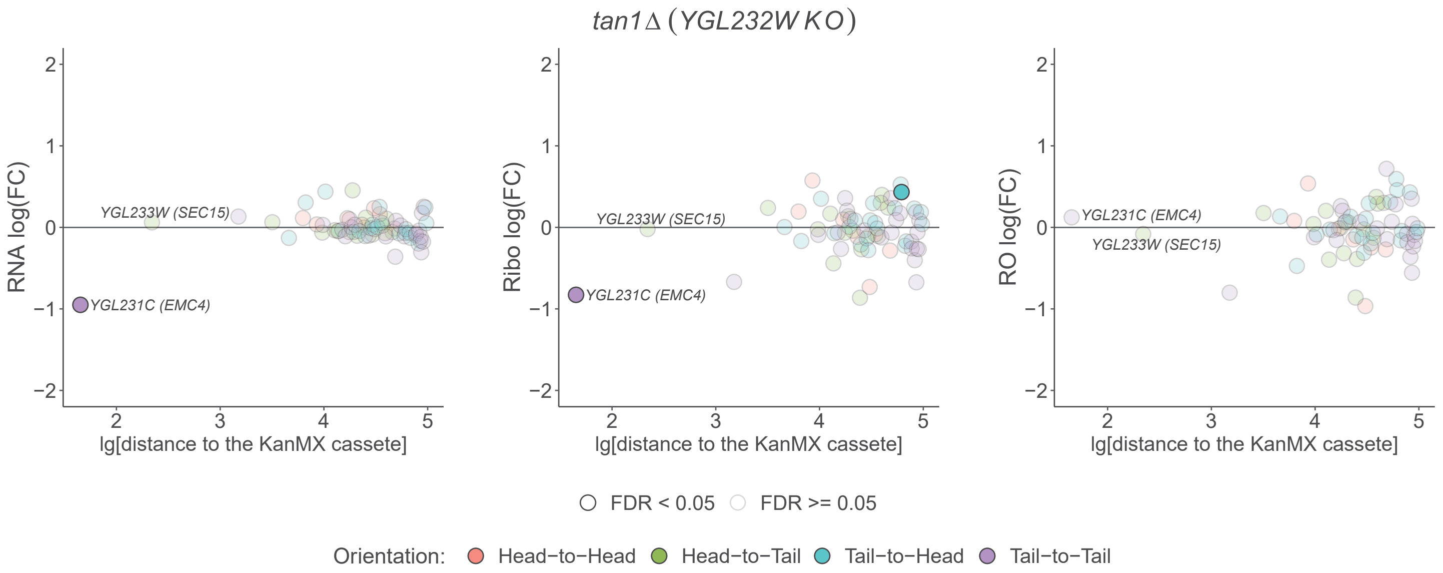

tan1Δ (YGL232W KO)
RNA log(FC)
Ribo log(FC)
RO log(FC)
lg[distance to the KanMX cassete]
YGL233W (SEC15)
YGL231C (EMC4)
FDR < 0.05
FDR >= 0.05
Orientation: Head−to−Head Head−to−Tail Tail−to−Head Tail−to−Tail


## B

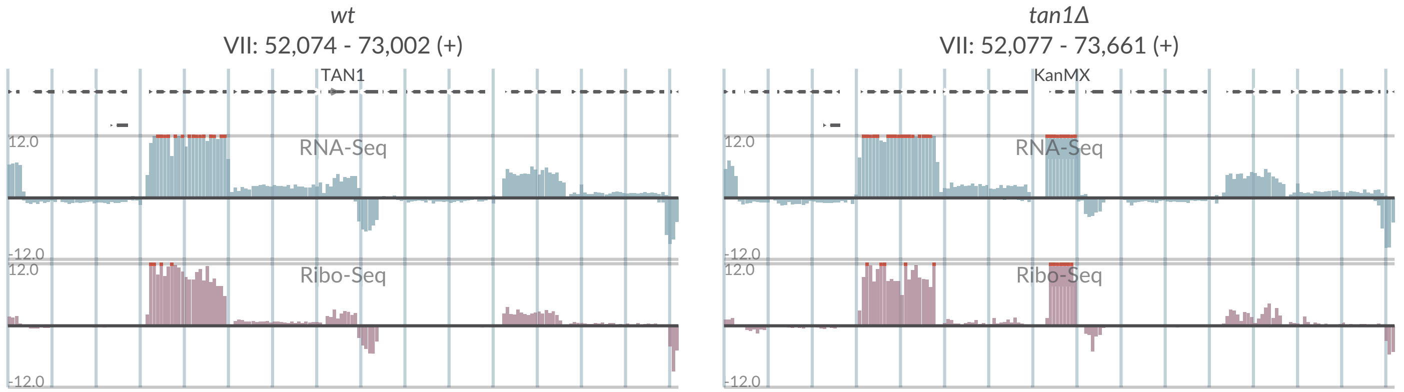

wt
VII: 52,074 - 73,002 (+)
TAN1
tan1Δ
VII: 52,077 - 73,661 (+)
KanMX
RNA-Seq
Ribo-Seq
12.0
-12.0

# S2-39

**A**

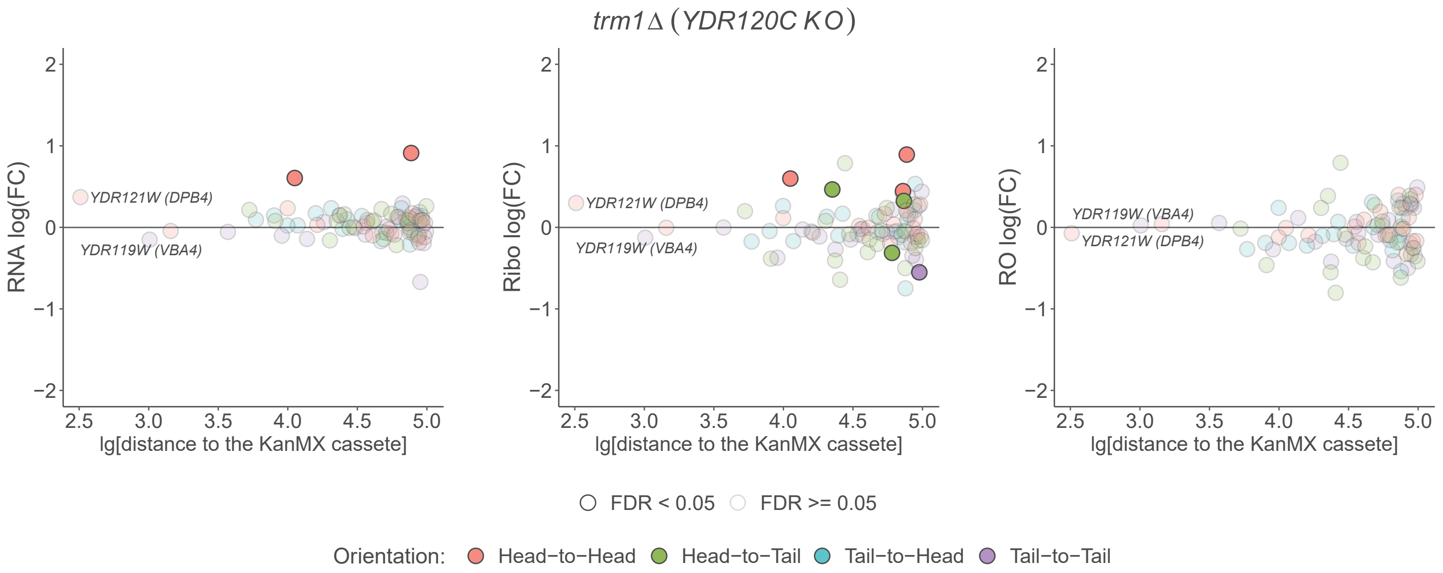
trm1Δ (YDR120C KO)
RNA log(FC)
Ribo log(FC)
RO log(FC)
YDR121W (DPB4)
YDR119W (VBA4)
lg[distance to the KanMX cassete]
FDR < 0.05
FDR >= 0.05
Orientation:
Head−to−Head
Head−to−Tail
Tail−to−Head
Tail−to−Tail

**B**

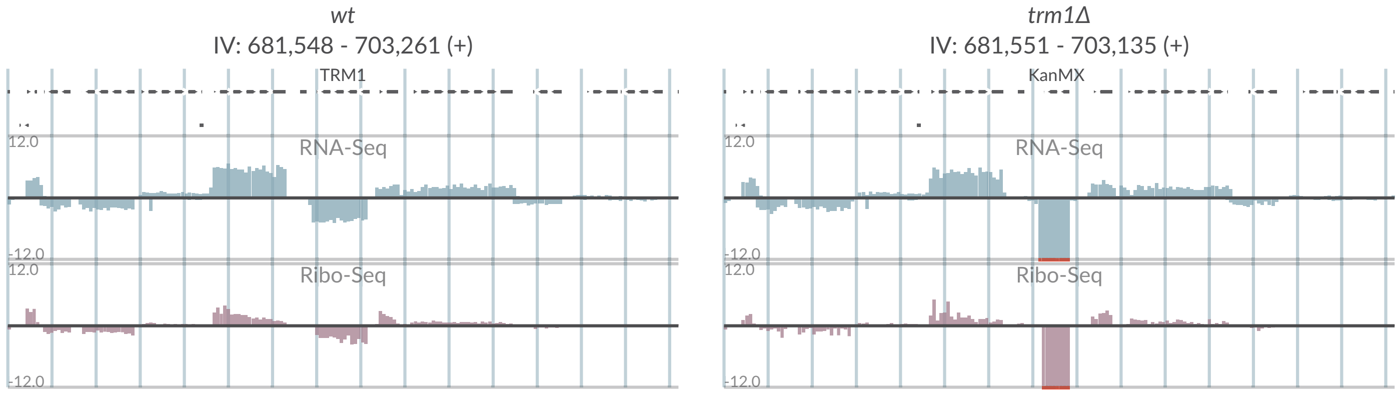
wt
IV: 681,548 - 703,261 (+)
TRM1
trm1Δ
IV: 681,551 - 703,135 (+)
KanMX
RNA-Seq
Ribo-Seq
12.0
-12.0

# S2-40

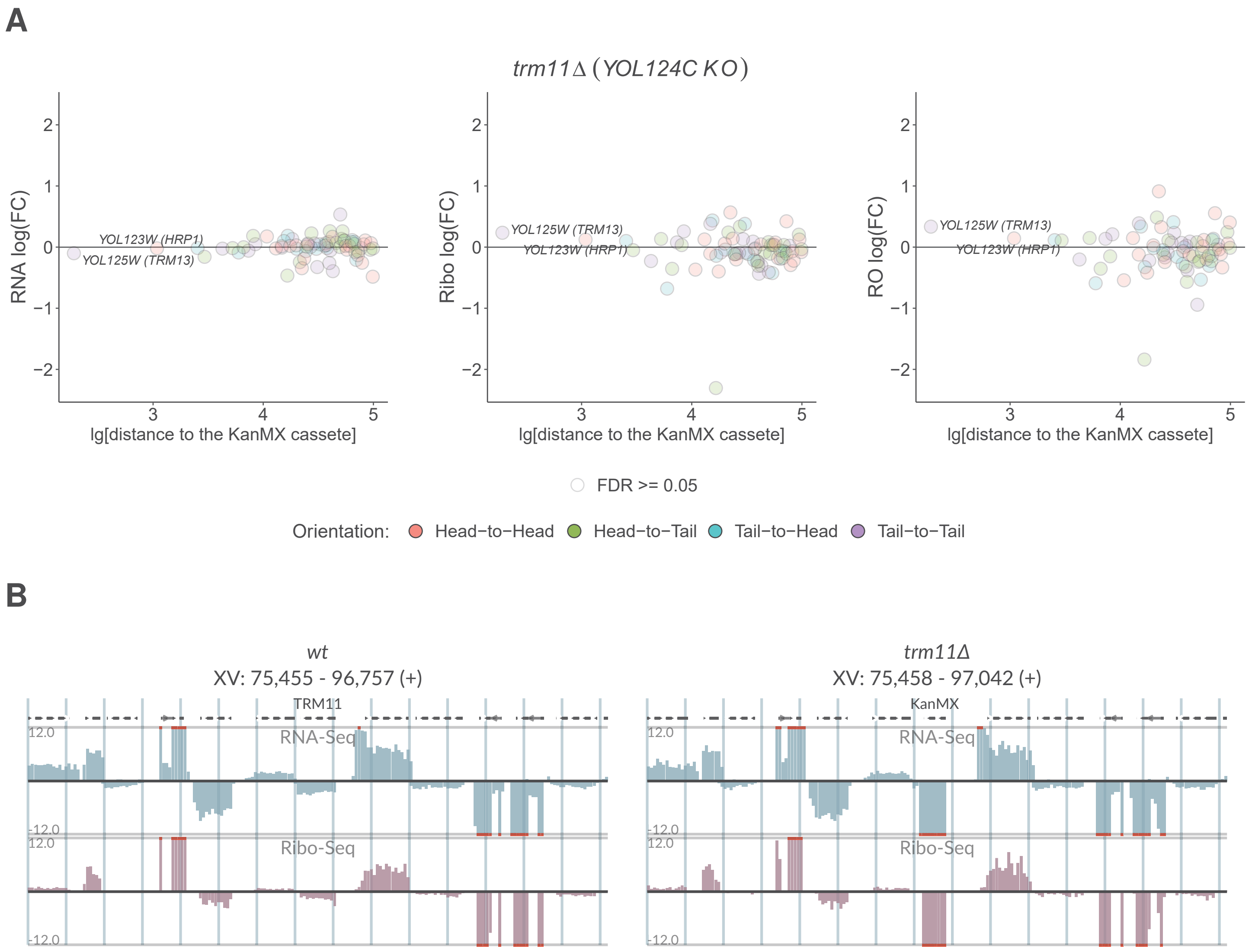
A
trm11Δ (YOL124C KO)
RNA log(FC)
Ribo log(FC)
RO log(FC)
YOL123W (HRP1)
YOL125W (TRM13)
lg[distance to the KanMX cassete]
FDR >= 0.05
Orientation: Head−to−Head Head−to−Tail Tail−to−Head Tail−to−Tail
B
wt
XV: 75,455 - 96,757 (+)
TRM11
trm11Δ
XV: 75,458 - 97,042 (+)
KanMX
RNA-Seq
Ribo-Seq
12.0
-12.0

# S2-41

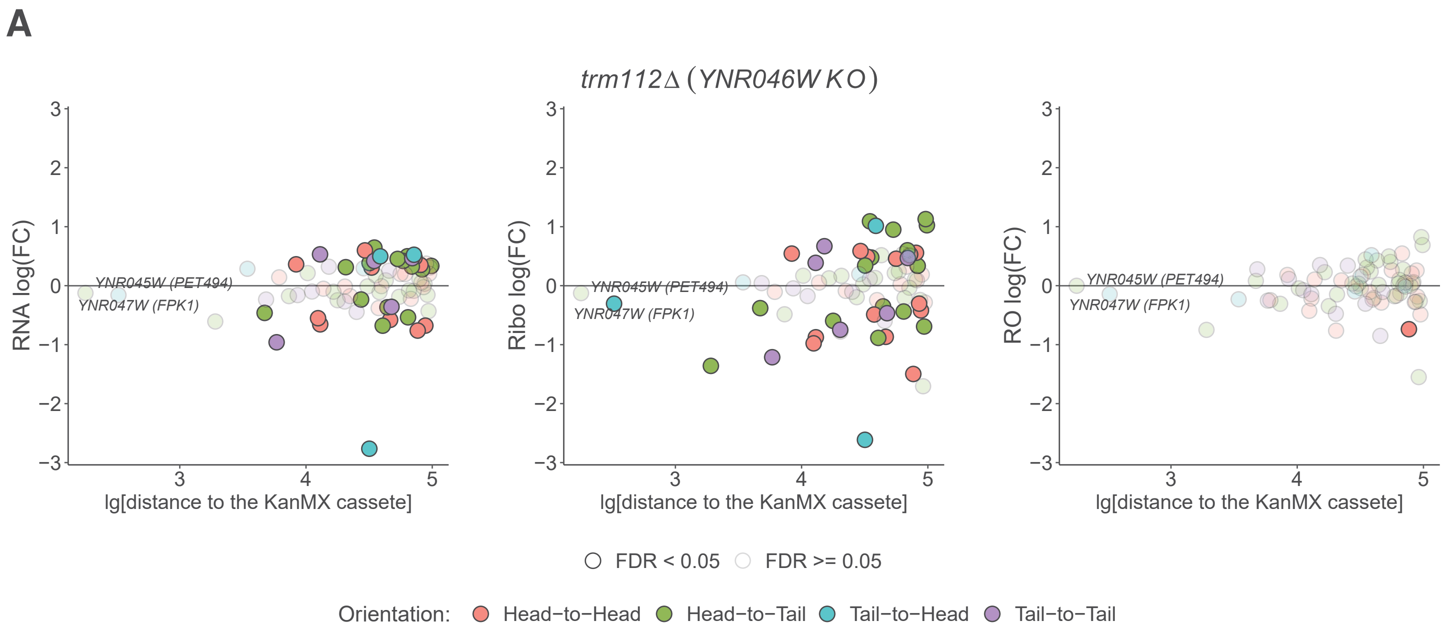
A
trm112Δ (YNR046W KO)
RNA log(FC)
Ribo log(FC)
RO log(FC)
YNR045W (PET494)
YNR047W (FPK1)
lg[distance to the KanMX cassete]
FDR < 0.05
FDR >= 0.05
Orientation:
Head−to−Head
Head−to−Tail
Tail−to−Head
Tail−to−Tail

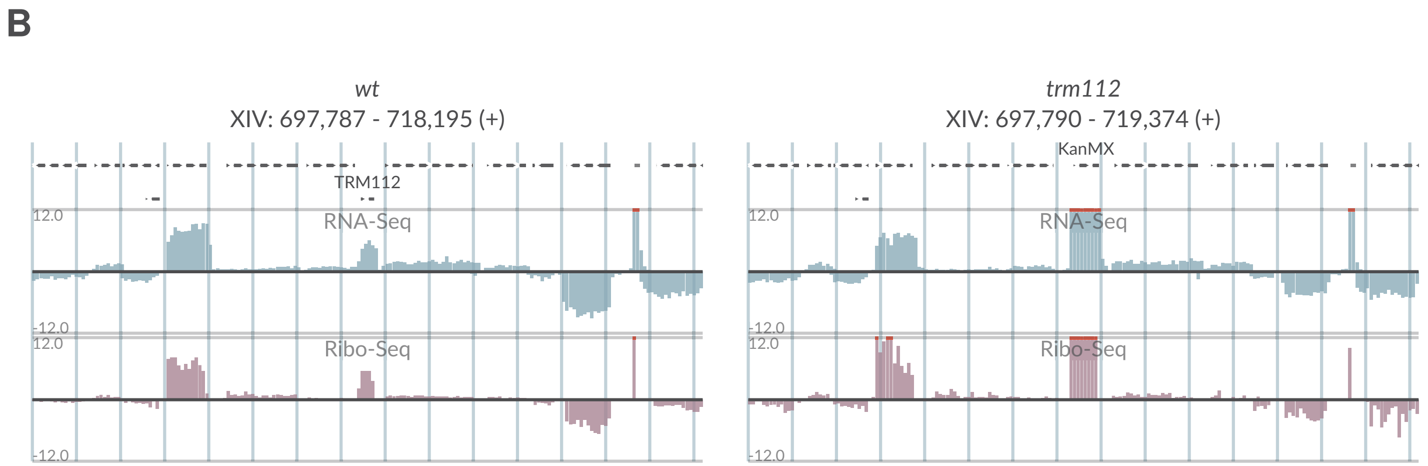
B
wt
XIV: 697,787 - 718,195 (+)
trm112
XIV: 697,790 - 719,374 (+)
TRM112
KanMX
RNA-Seq
Ribo-Seq
12.0
-12.0

# S2-42

**A**

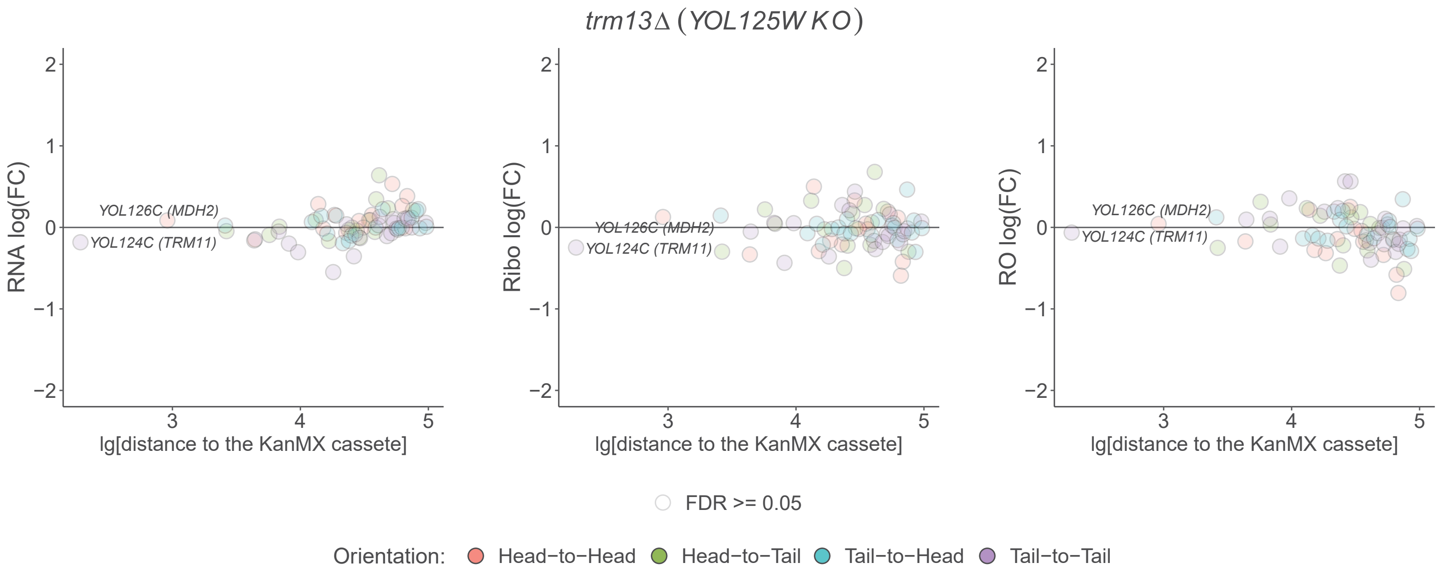

trm13Δ (YOL125W KO)
RNA log(FC)
Ribo log(FC)
RO log(FC)
YOL126C (MDH2)
YOL124C (TRM11)
lg[distance to the KanMX cassete]
FDR >= 0.05
Orientation: Head−to−Head Head−to−Tail Tail−to−Head Tail−to−Tail


**B**

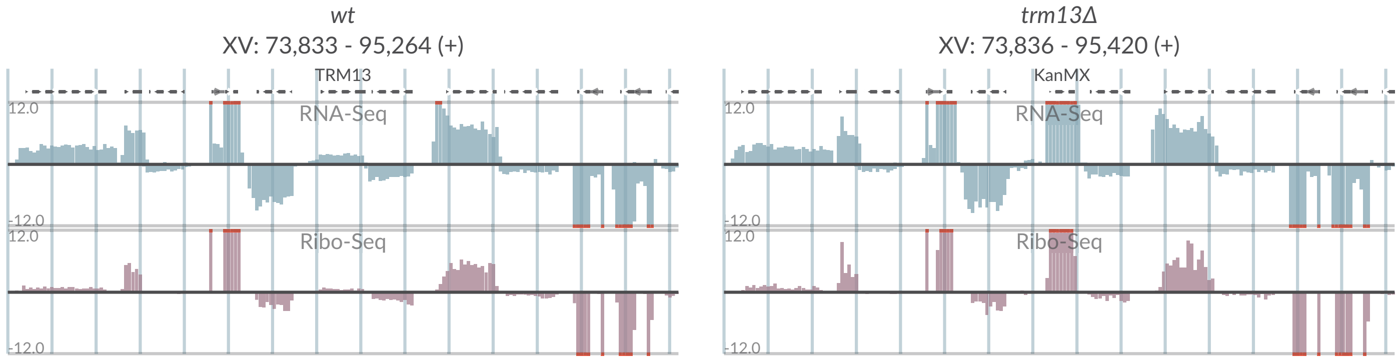

wt
XV: 73,833 - 95,264 (+)
TRM13
RNA-Seq
Ribo-Seq
trm13Δ
XV: 73,836 - 95,420 (+)
KanMX
RNA-Seq
Ribo-Seq

# S2-43

**A**

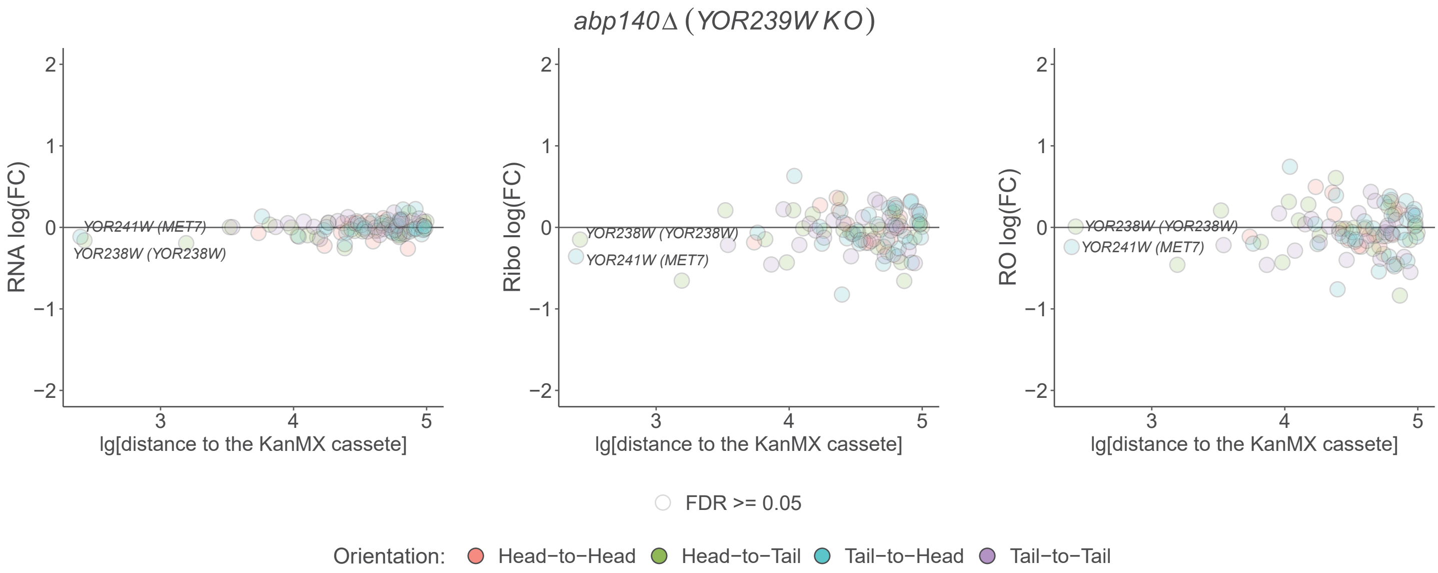

abp140Δ (YOR239W KO)
RNA log(FC)
Ribo log(FC)
RO log(FC)
lg[distance to the KanMX cassete]
YOR241W (MET7)
YOR238W (YOR238W)
FDR >= 0.05
Orientation: Head−to−Head Head−to−Tail Tail−to−Head Tail−to−Tail


**B**

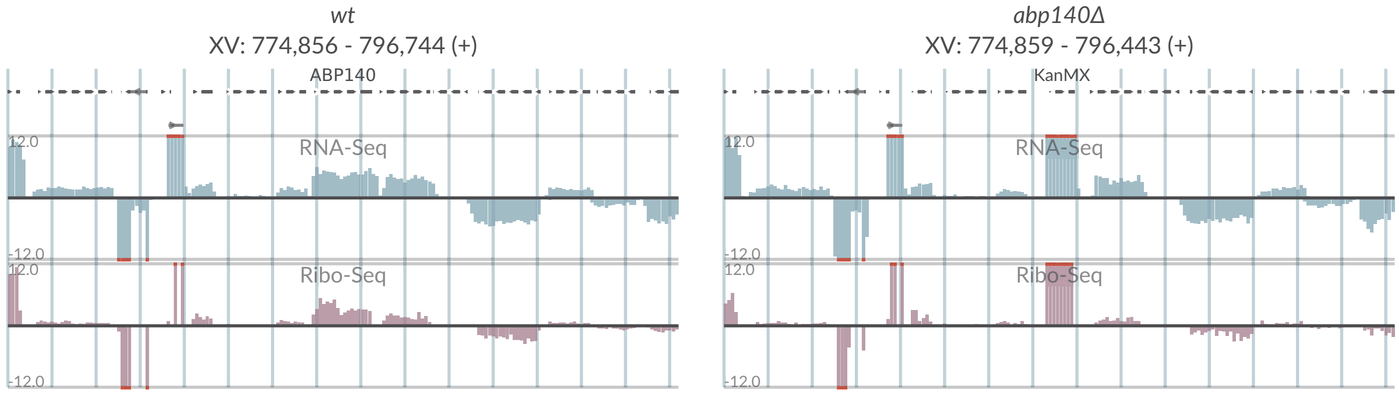

wt
XV: 774,856 - 796,744 (+)
ABP140
abp140Δ
XV: 774,859 - 796,443 (+)
KanMX
RNA-Seq
Ribo-Seq
12.0
-12.0

# S2-44

## A

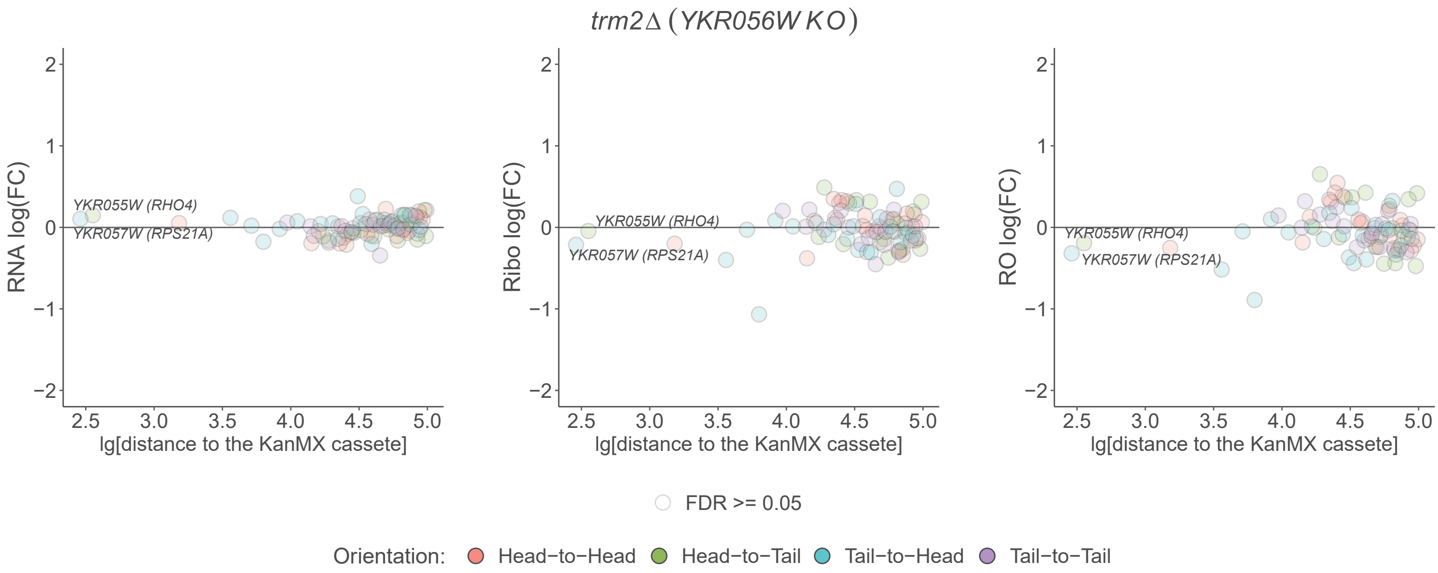

trm2Δ (YKR056W KO)
RNA log(FC)
Ribo log(FC)
RO log(FC)
YKR055W (RHO4)
YKR057W (RPS21A)
lg[distance to the KanMX cassete]
2
1
0
−1
−2
2.5
3.0
3.5
4.0
4.5
5.0
FDR >= 0.05
Orientation:
Head−to−Head
Head−to−Tail
Tail−to−Head
Tail−to−Tail


## B

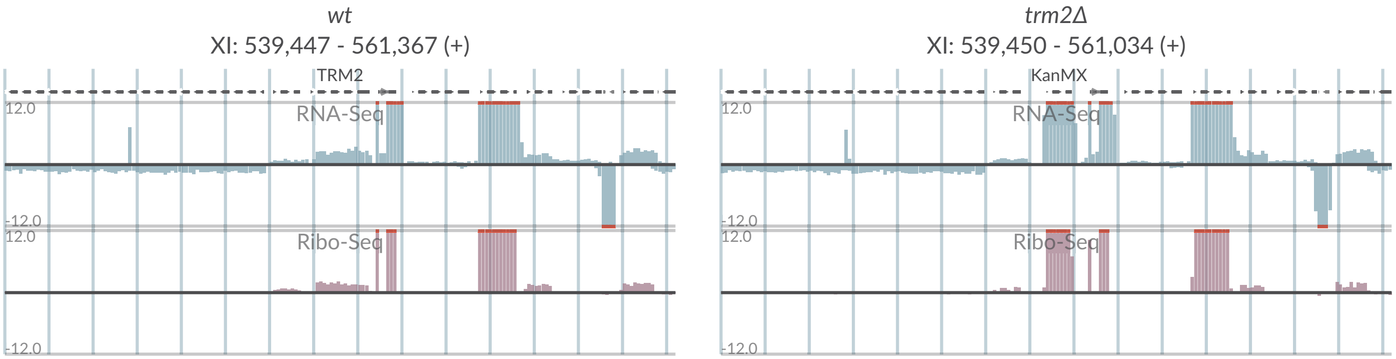

wt
XI: 539,447 - 561,367 (+)
TRM2
RNA-Seq
Ribo-Seq
12.0
-12.0
trm2Δ
XI: 539,450 - 561,034 (+)
KanMX
RNA-Seq
Ribo-Seq
12.0
-12.0

# S2-45

## A

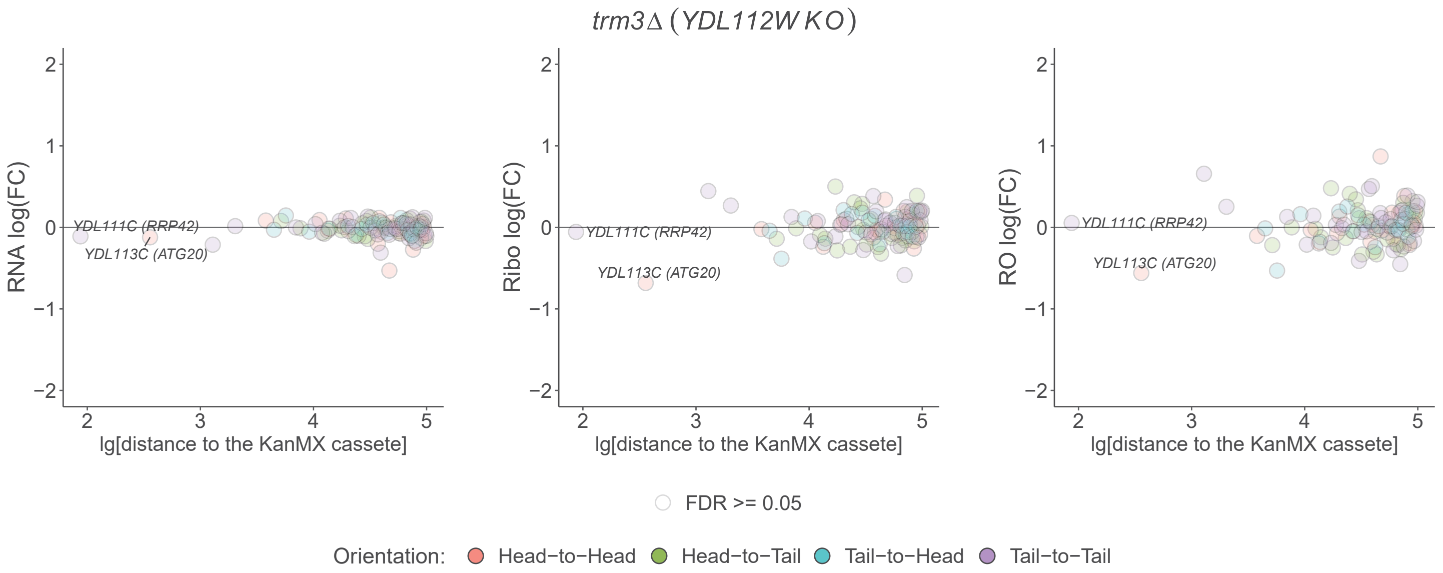
trm3Δ (YDL112W KO)
RNA log(FC)
Ribo log(FC)
RO log(FC)
YDL111C (RRP42)
YDL113C (ATG20)
lg[distance to the KanMX cassete]
FDR >= 0.05
Orientation: Head−to−Head Head−to−Tail Tail−to−Head Tail−to−Tail

## B

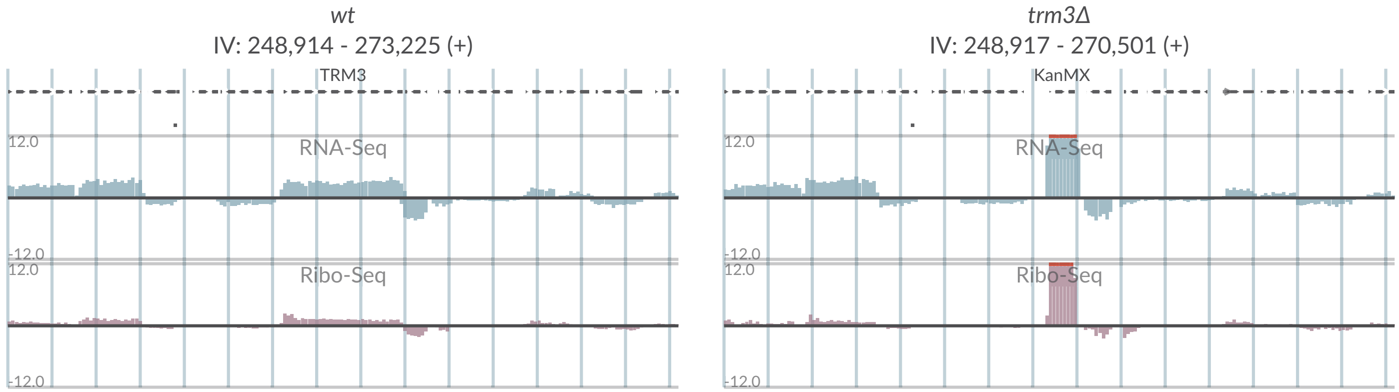
wt
IV: 248,914 - 273,225 (+)
TRM3
RNA-Seq
Ribo-Seq
trm3Δ
IV: 248,917 - 270,501 (+)
KanMX
RNA-Seq
Ribo-Seq

# S2-46

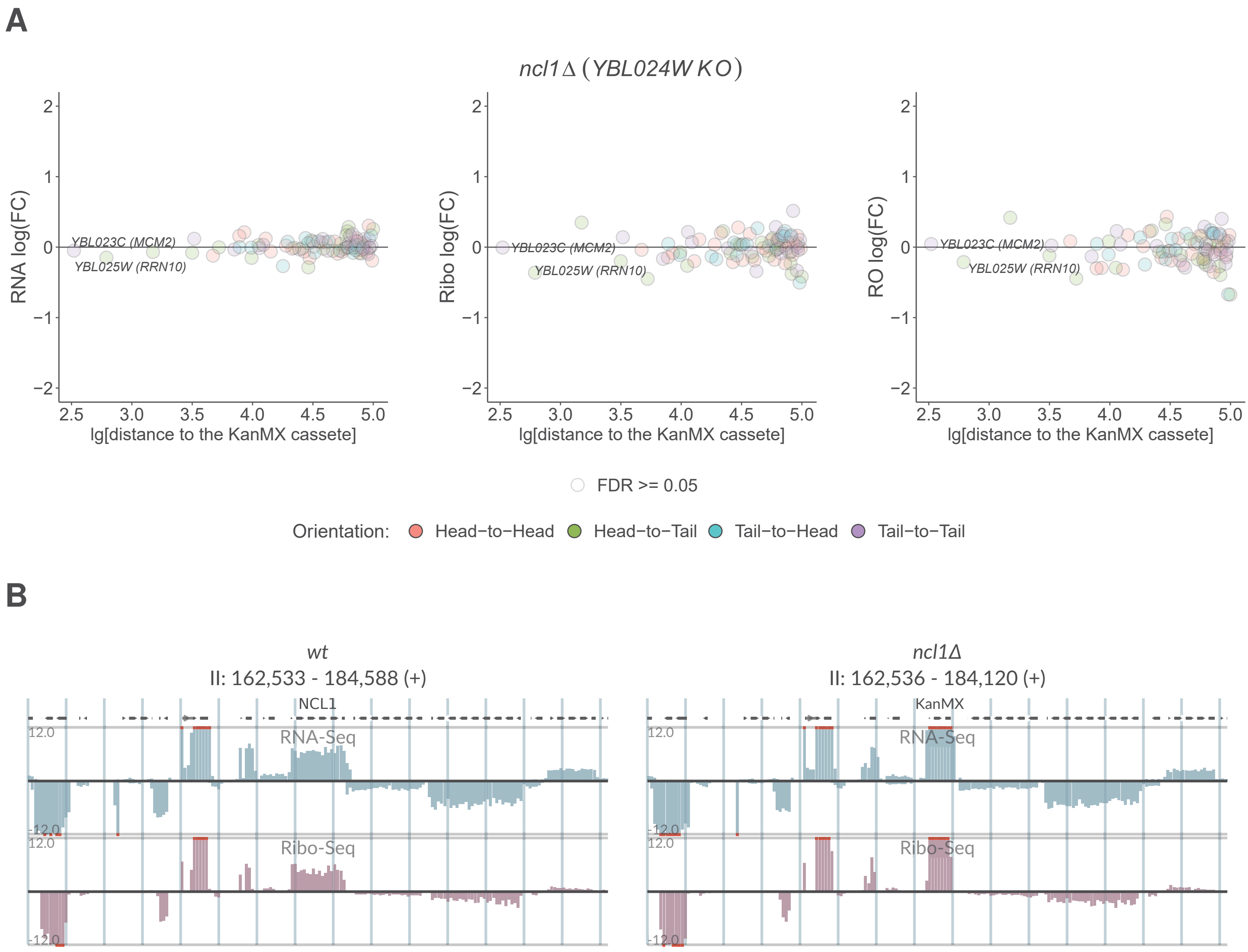
A
ncl1Δ (YBL024W KO)
RNA log(FC)
Ribo log(FC)
RO log(FC)
YBL023C (MCM2)
YBL025W (RRN10)
lg[distance to the KanMX cassete]
FDR >= 0.05
Orientation: Head−to−Head Head−to−Tail Tail−to−Head Tail−to−Tail
B
wt
II: 162,533 - 184,588 (+)
ncl1Δ
II: 162,536 - 184,120 (+)
NCL1
KanMX
RNA-Seq
Ribo-Seq
12.0
-12.0

# S2-47

## A

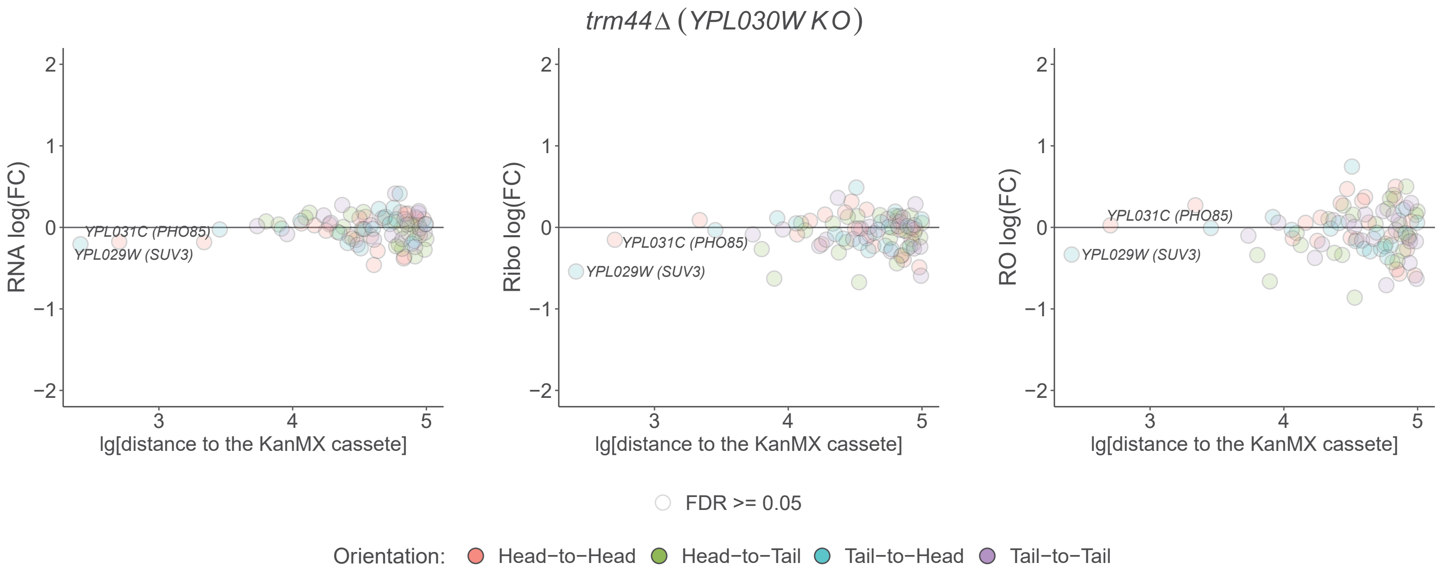

*trm44Δ (YPL030W KO)*
RNA log(FC)
Ribo log(FC)
RO log(FC)
YPL031C (PHO85)
YPL029W (SUV3)
lg[distance to the KanMX cassete]
FDR >= 0.05
Orientation: Head−to−Head Head−to−Tail Tail−to−Head Tail−to−Tail


## B

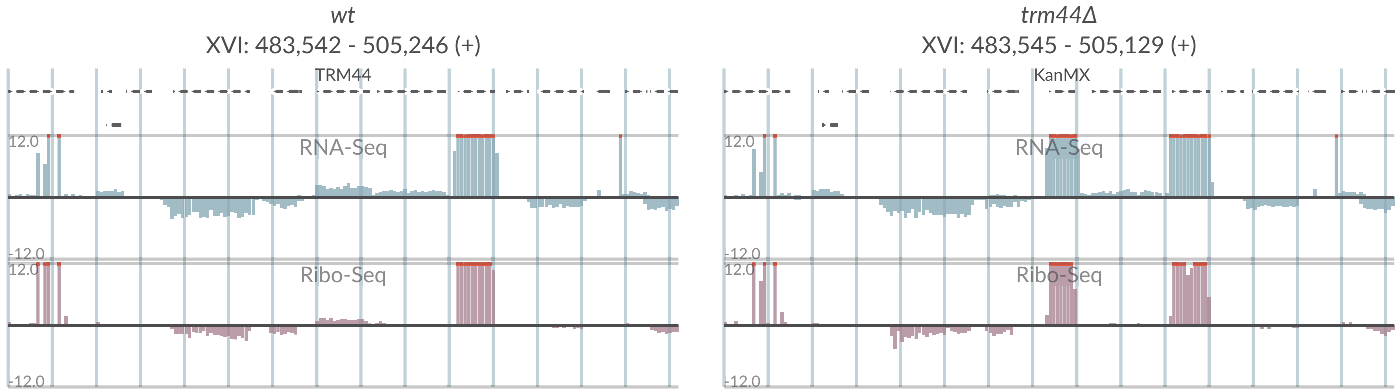

*wt*
XVI: 483,542 - 505,246 (+)
TRM44
RNA-Seq
Ribo-Seq
*trm44Δ*
XVI: 483,545 - 505,129 (+)
KanMX
RNA-Seq
Ribo-Seq

# S2-48

## A

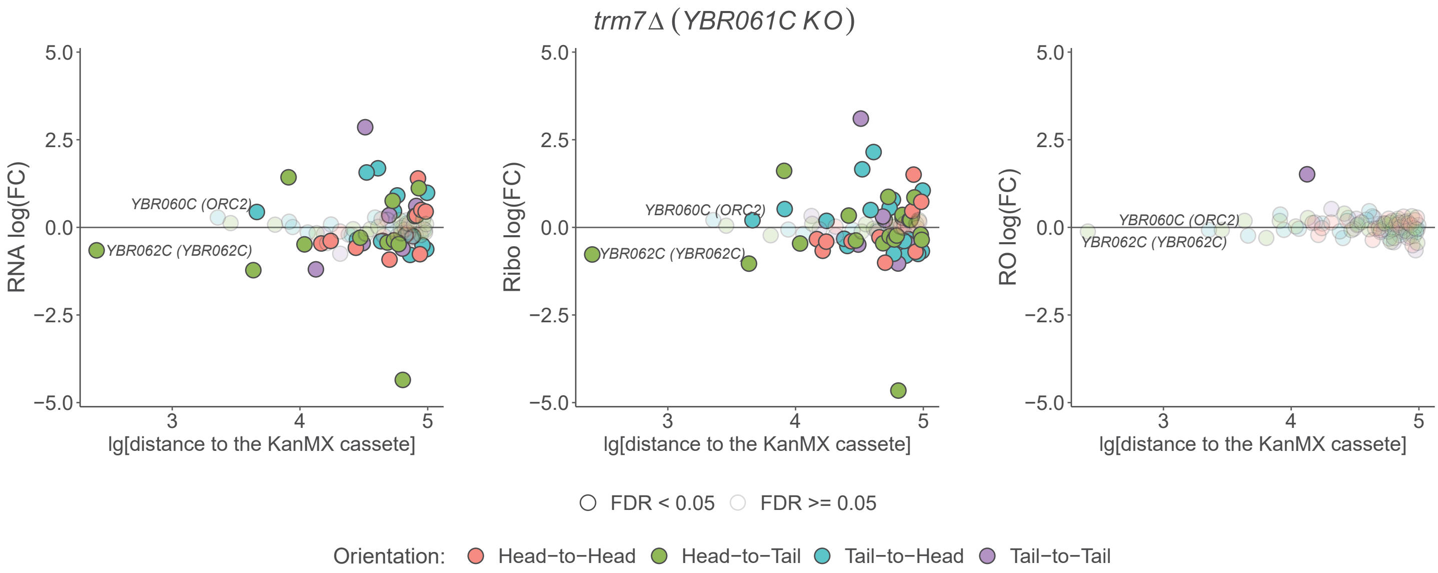
trm7Δ (YBR061C KO)
RNA log(FC)
Ribo log(FC)
RO log(FC)
lg[distance to the KanMX cassete]
YBR060C (ORC2)
YBR062C (YBR062C)
FDR < 0.05
FDR >= 0.05
Orientation:
Head−to−Head
Head−to−Tail
Tail−to−Head
Tail−to−Tail

## B

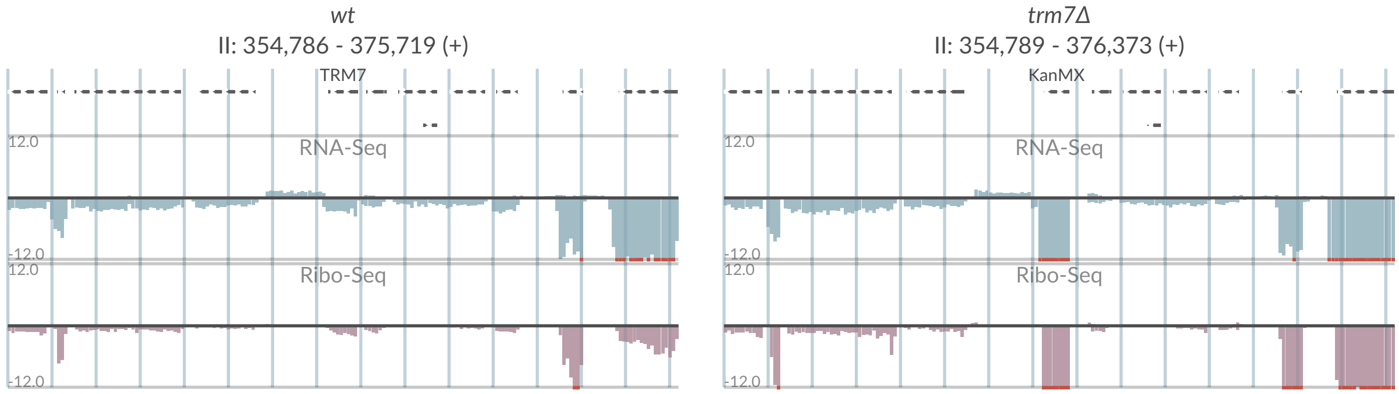
wt
II: 354,786 - 375,719 (+)
TRM7
trm7Δ
II: 354,789 - 376,373 (+)
KanMX
RNA-Seq
Ribo-Seq
12.0
-12.0

# S2-49

**A**

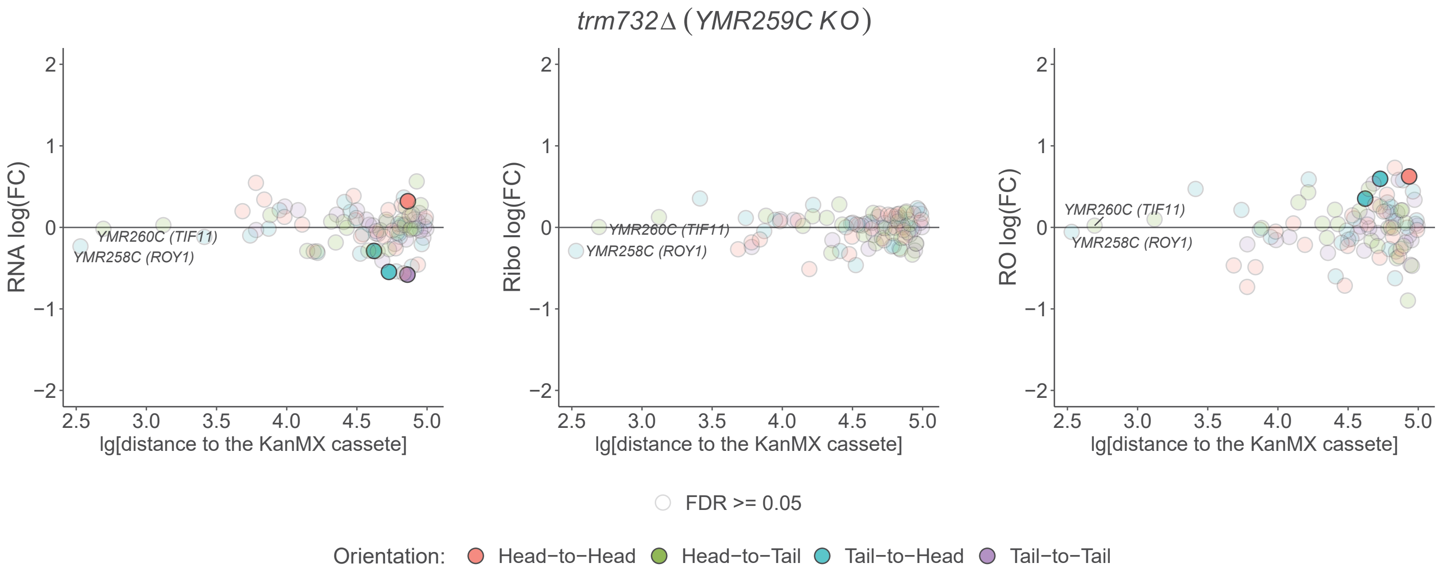
trm732Δ (YMR259C KO)
RNA log(FC)
Ribo log(FC)
RO log(FC)
lg[distance to the KanMX cassete]
YMR260C (TIF11)
YMR258C (ROY1)
FDR >= 0.05
Orientation: Head−to−Head Head−to−Tail Tail−to−Head Tail−to−Tail

**B**

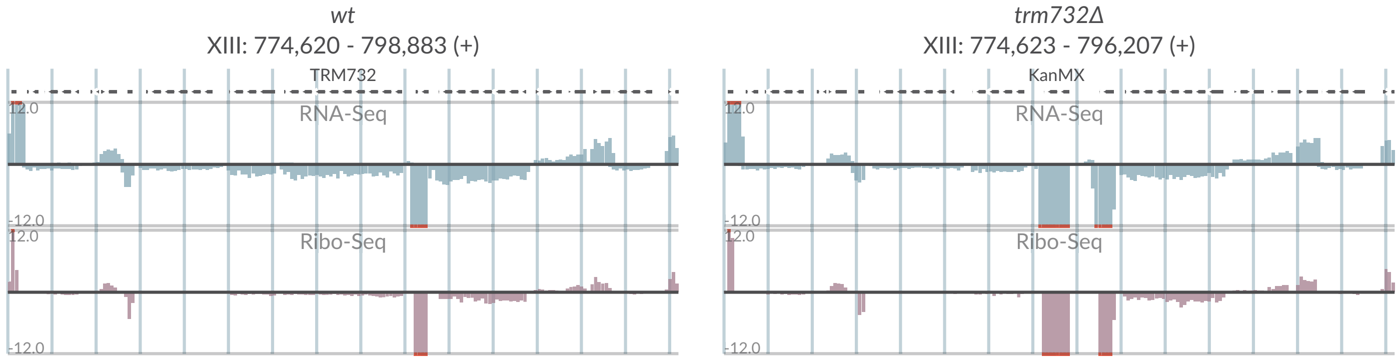
wt
XIII: 774,620 - 798,883 (+)
TRM732
RNA-Seq
Ribo-Seq
12.0
-12.0
trm732Δ
XIII: 774,623 - 796,207 (+)
KanMX
RNA-Seq
Ribo-Seq
12.0
-12.0

# S2-50

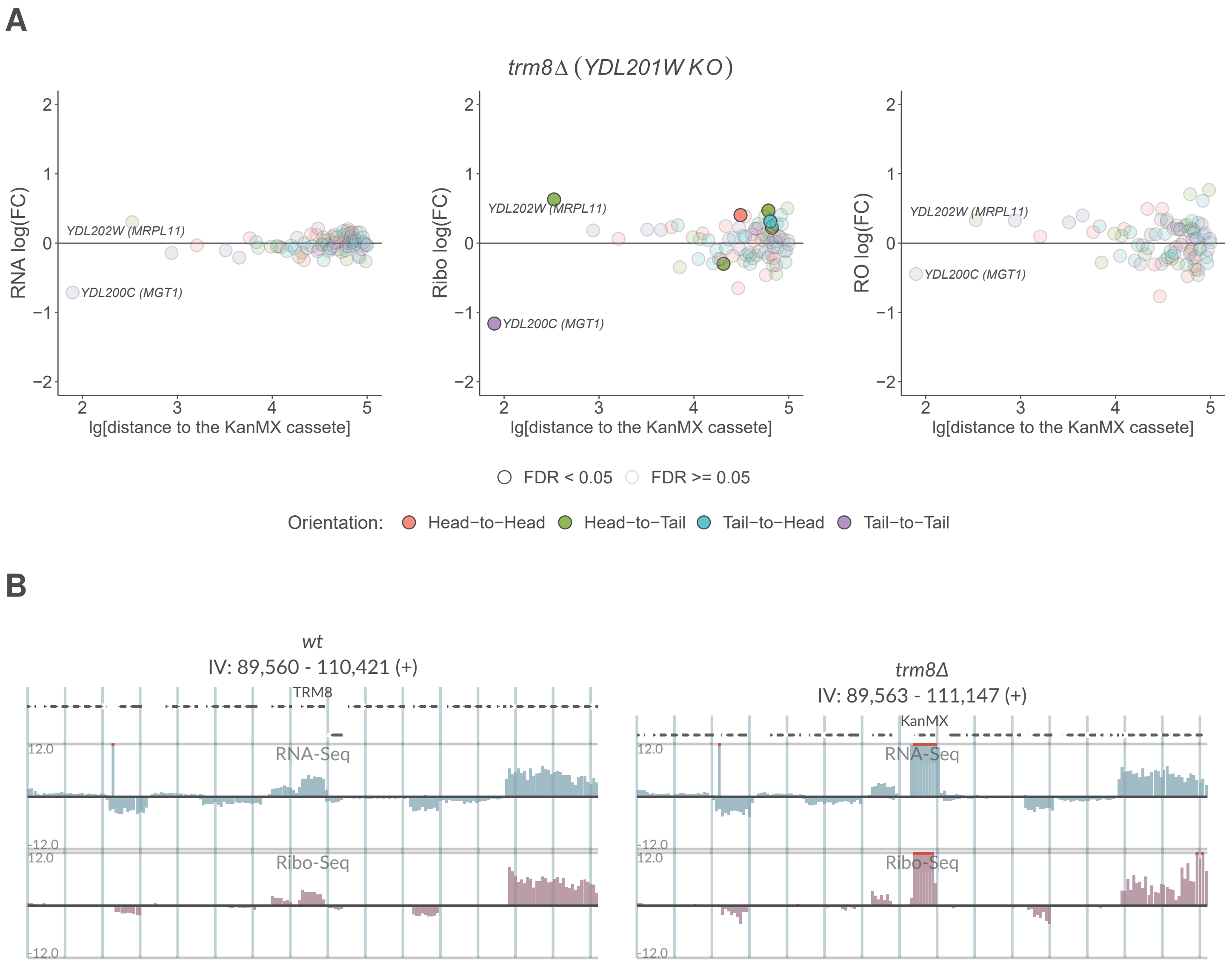
A
trm8Δ (YDL201W KO)
RNA log(FC)
Ribo log(FC)
RO log(FC)
lg[distance to the KanMX cassete]
YDL202W (MRPL11)
YDL200C (MGT1)
FDR < 0.05
FDR >= 0.05
Orientation:
Head−to−Head
Head−to−Tail
Tail−to−Head
Tail−to−Tail
B
wt
IV: 89,560 - 110,421 (+)
TRM8
trm8Δ
IV: 89,563 - 111,147 (+)
KanMX
RNA-Seq
Ribo-Seq
12.0
-12.0

# S2-51

**A**

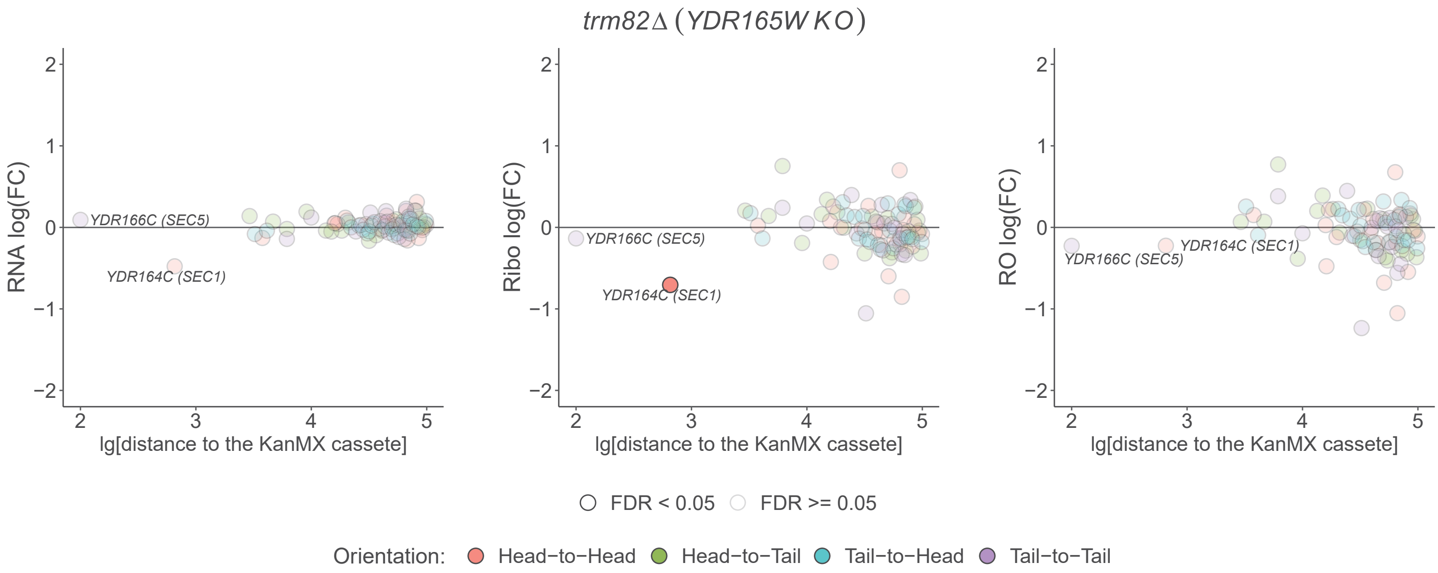
*trm82Δ* (*YDR165W KO*)
RNA log(FC)
Ribo log(FC)
RO log(FC)
lg[distance to the KanMX cassete]
YDR166C (SEC5)
YDR164C (SEC1)
FDR < 0.05
FDR >= 0.05
Orientation:
Head−to−Head
Head−to−Tail
Tail−to−Head
Tail−to−Tail

**B**

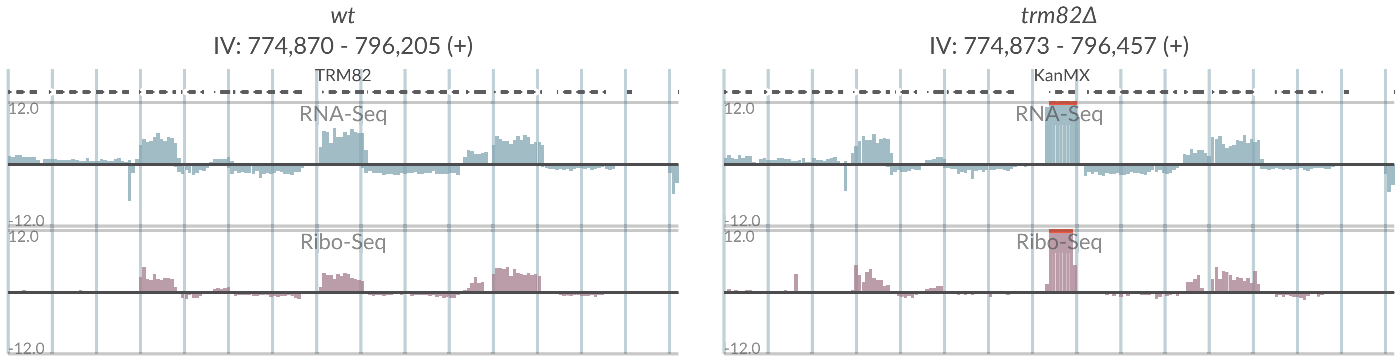
*wt*
IV: 774,870 - 796,205 (+)
TRM82
RNA-Seq
Ribo-Seq
*trm82Δ*
IV: 774,873 - 796,457 (+)
KanMX
RNA-Seq
Ribo-Seq

# S2-52

**A**

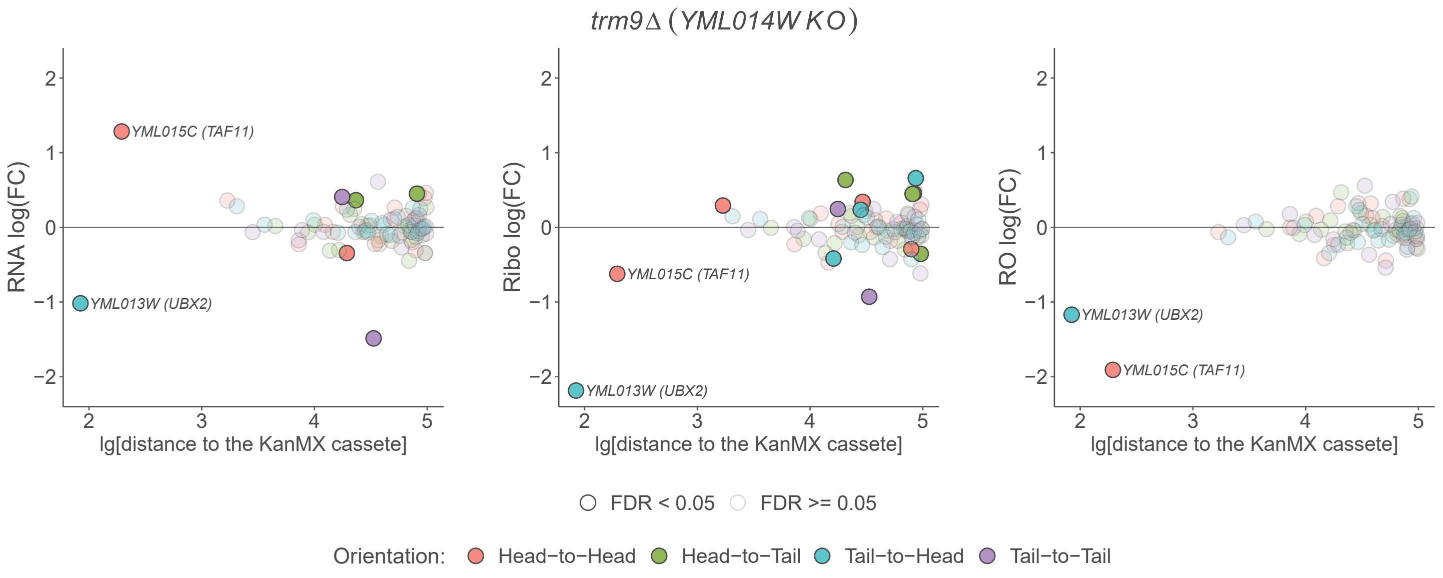

trm9Δ (YML014W KO)
RNA log(FC)
Ribo log(FC)
RO log(FC)
YML015C (TAF11)
YML013W (UBX2)
lg[distance to the KanMX cassete]
FDR < 0.05
FDR >= 0.05
Orientation:
Head−to−Head
Head−to−Tail
Tail−to−Head
Tail−to−Tail


**B**

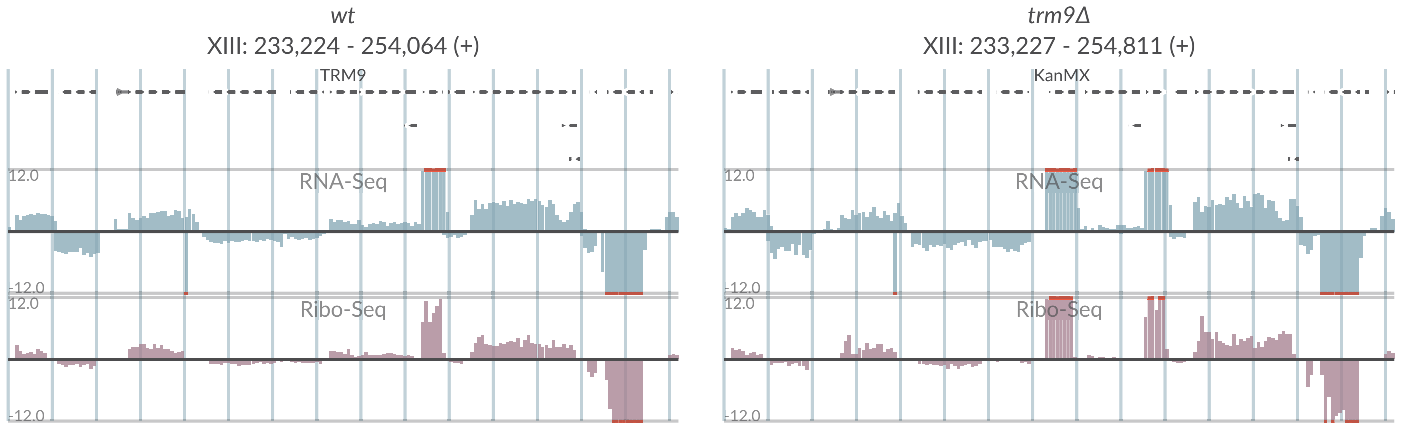

wt
XIII: 233,224 - 254,064 (+)
trm9Δ
XIII: 233,227 - 254,811 (+)
TRM9
KanMX
RNA-Seq
Ribo-Seq
12.0
-12.0

# S2-53

## A

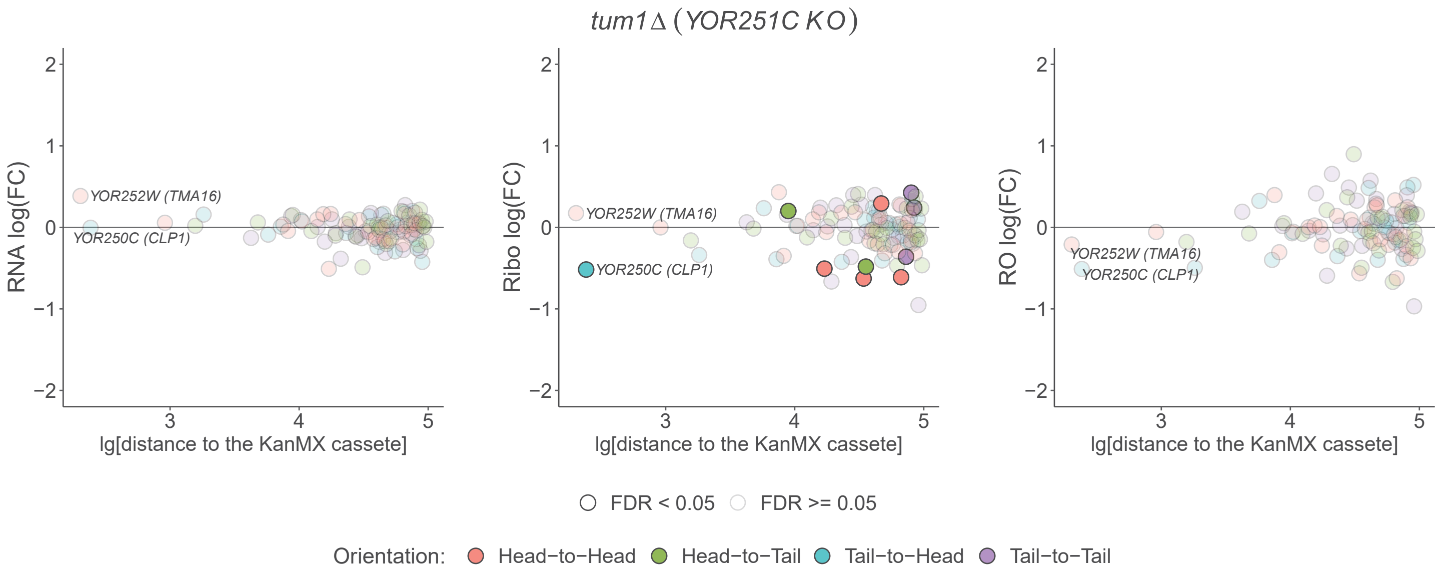

tum1Δ (YOR251C KO)
RNA log(FC)
Ribo log(FC)
RO log(FC)
YOR252W (TMA16)
YOR250C (CLP1)
lg[distance to the KanMX cassete]
FDR < 0.05
FDR >= 0.05
Orientation: Head−to−Head Head−to−Tail Tail−to−Head Tail−to−Tail


## B

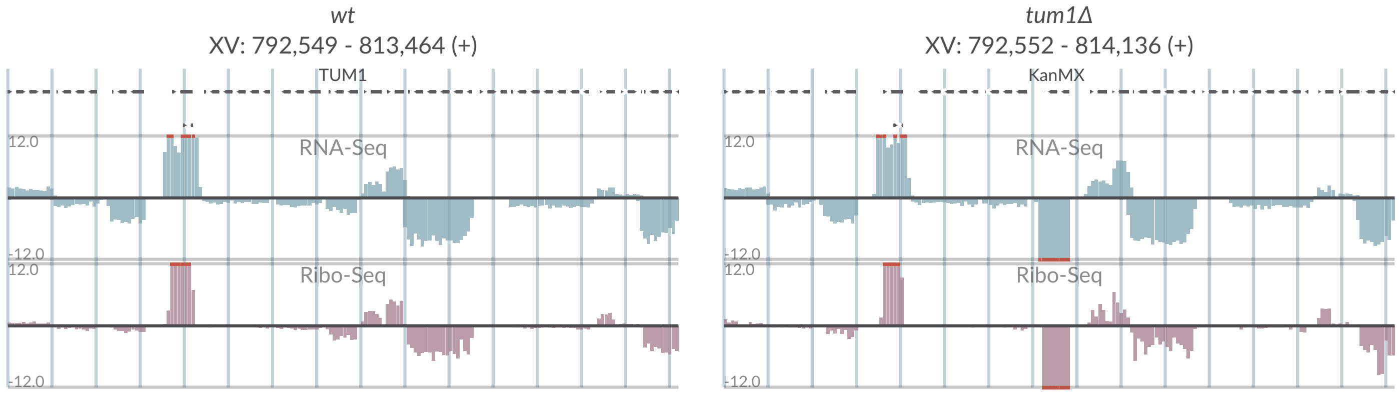

wt
XV: 792,549 - 813,464 (+)
TUM1
tum1Δ
XV: 792,552 - 814,136 (+)
KanMX
RNA-Seq
Ribo-Seq
12.0
-12.0

# S2-54

**A**

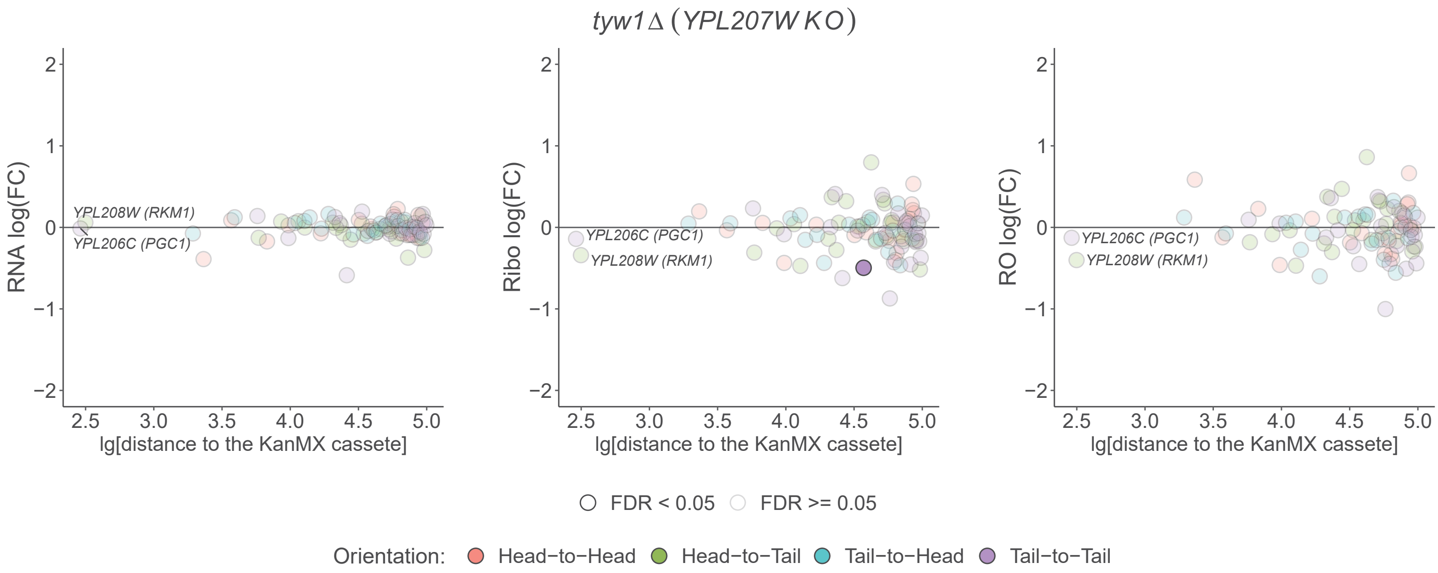
tyw1Δ (YPL207W KO)
RNA log(FC)
Ribo log(FC)
RO log(FC)
YPL208W (RKM1)
YPL206C (PGC1)
lg[distance to the KanMX cassete]
2
1
0
−1
−2
2.5
3.0
3.5
4.0
4.5
5.0
FDR < 0.05
FDR >= 0.05
Orientation:
Head−to−Head
Head−to−Tail
Tail−to−Head
Tail−to−Tail

**B**

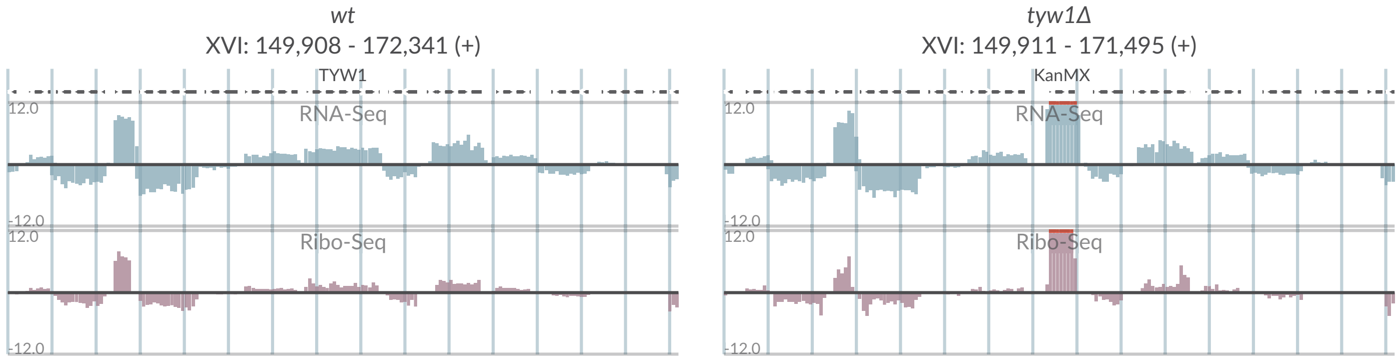
wt
XVI: 149,908 - 172,341 (+)
TYW1
tyw1Δ
XVI: 149,911 - 171,495 (+)
KanMX
RNA-Seq
Ribo-Seq
12.0
-12.0

# S2-55

**A**

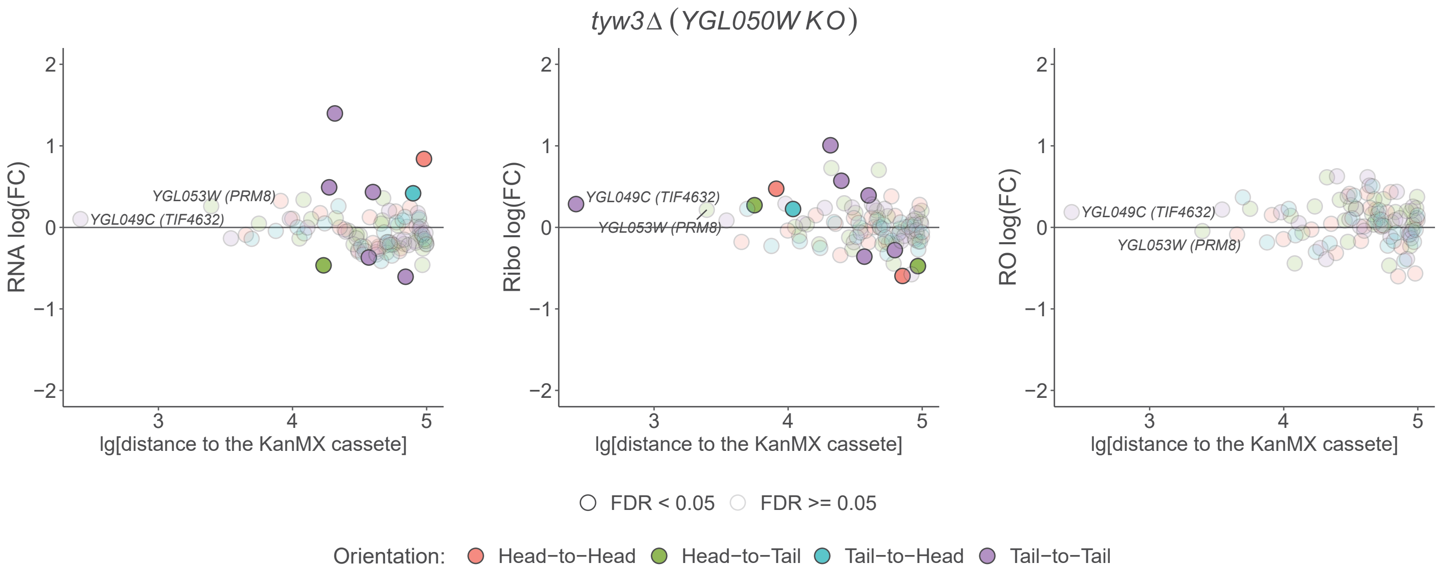

tyw3Δ (YGL050W KO)
RNA log(FC)
Ribo log(FC)
RO log(FC)
YGL053W (PRM8)
YGL049C (TIF4632)
lg[distance to the KanMX cassete]
FDR < 0.05
FDR >= 0.05
Orientation:
Head−to−Head
Head−to−Tail
Tail−to−Head
Tail−to−Tail


**B**

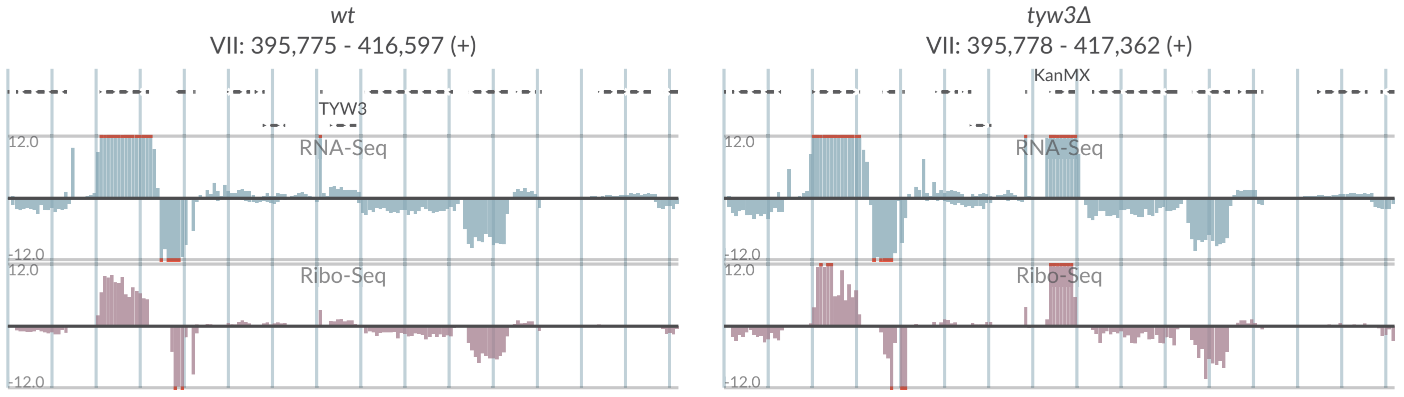

wt
VII: 395,775 - 416,597 (+)
TYW3
RNA-Seq
Ribo-Seq
tyw3Δ
VII: 395,778 - 417,362 (+)
KanMX

# S2-56

**A**

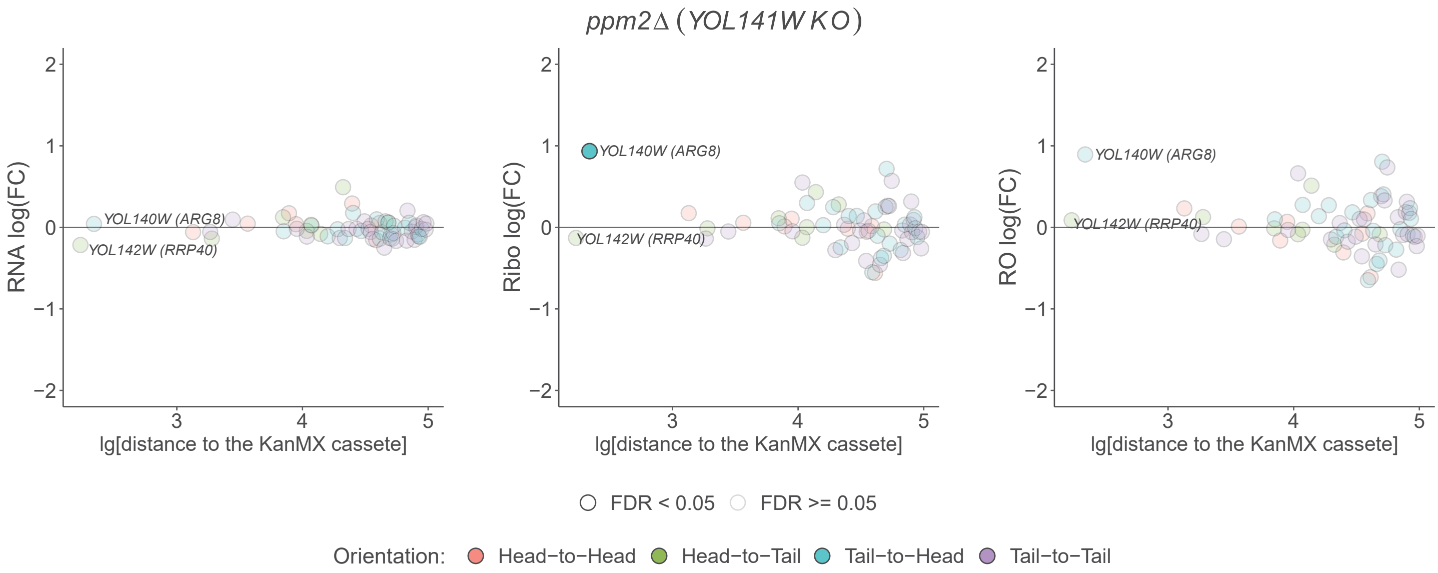

ppm2Δ (YOL141W KO)
RNA log(FC)
Ribo log(FC)
RO log(FC)
YOL140W (ARG8)
YOL142W (RRP40)
lg[distance to the KanMX cassete]
FDR < 0.05
FDR >= 0.05
Orientation:
Head−to−Head
Head−to−Tail
Tail−to−Head
Tail−to−Tail


**B**

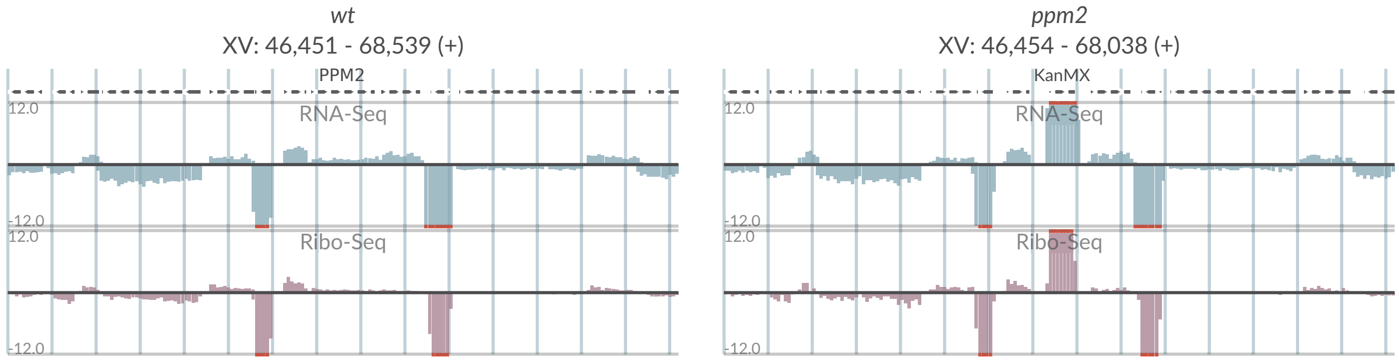

wt
XV: 46,451 - 68,539 (+)
PPM2
RNA-Seq
Ribo-Seq
ppm2
XV: 46,454 - 68,038 (+)
KanMX

# S2-57

**A**

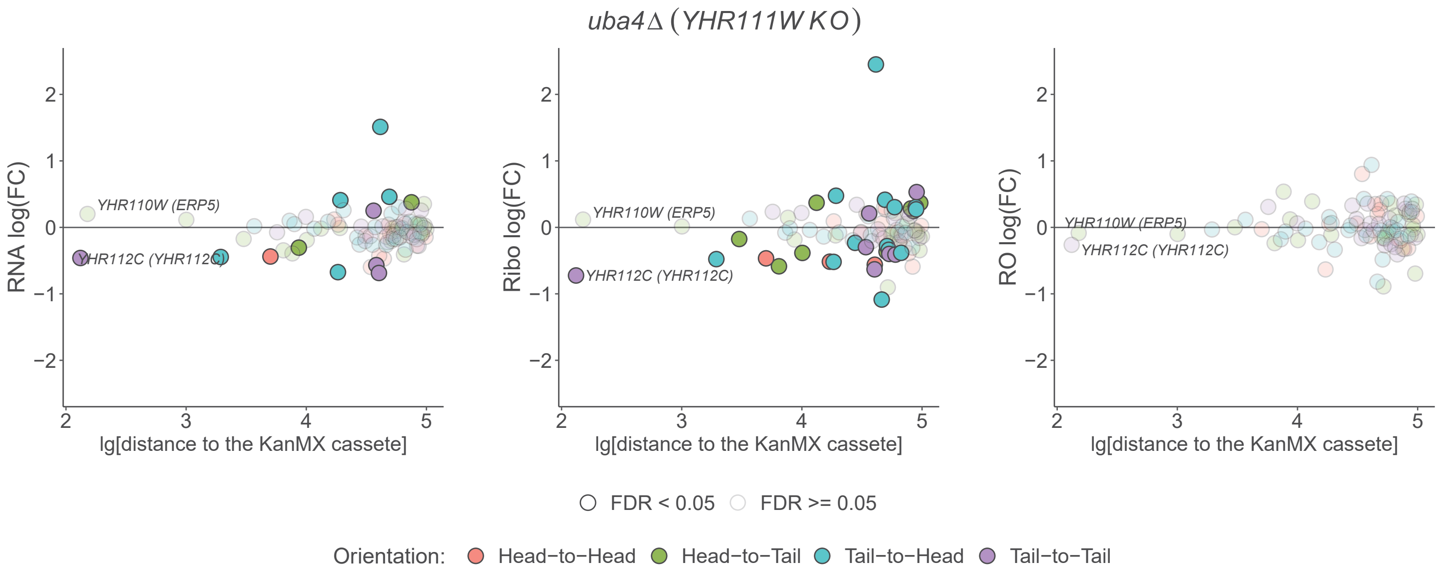

uba4Δ (YHR111W KO)
RNA log(FC)
Ribo log(FC)
RO log(FC)
YHR110W (ERP5)
YHR112C (YHR112C)
lg[distance to the KanMX cassete]
FDR < 0.05
FDR >= 0.05
Orientation:
Head−to−Head
Head−to−Tail
Tail−to−Head
Tail−to−Tail


**B**

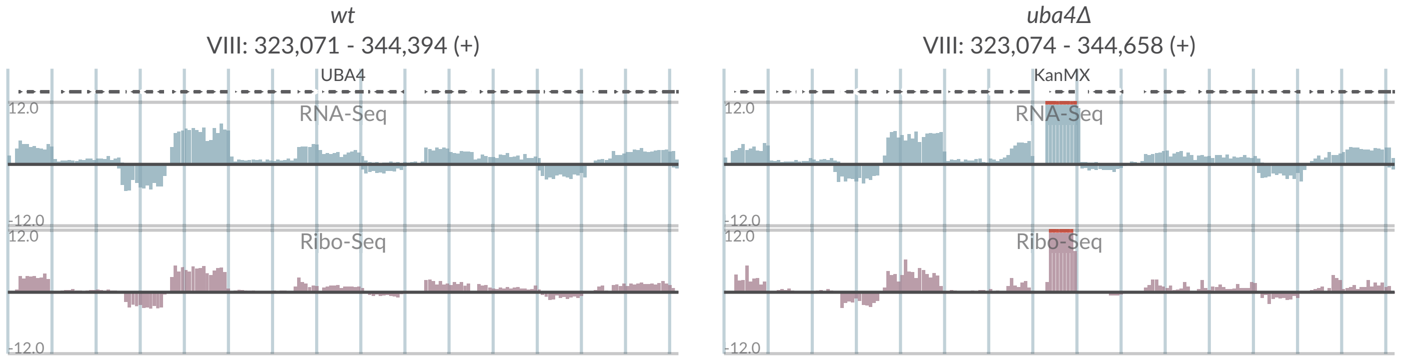

wt
VIII: 323,071 - 344,394 (+)
UBA4
RNA-Seq
Ribo-Seq
uba4Δ
VIII: 323,074 - 344,658 (+)
KanMX
RNA-Seq
Ribo-Seq

# S2-58

**A**

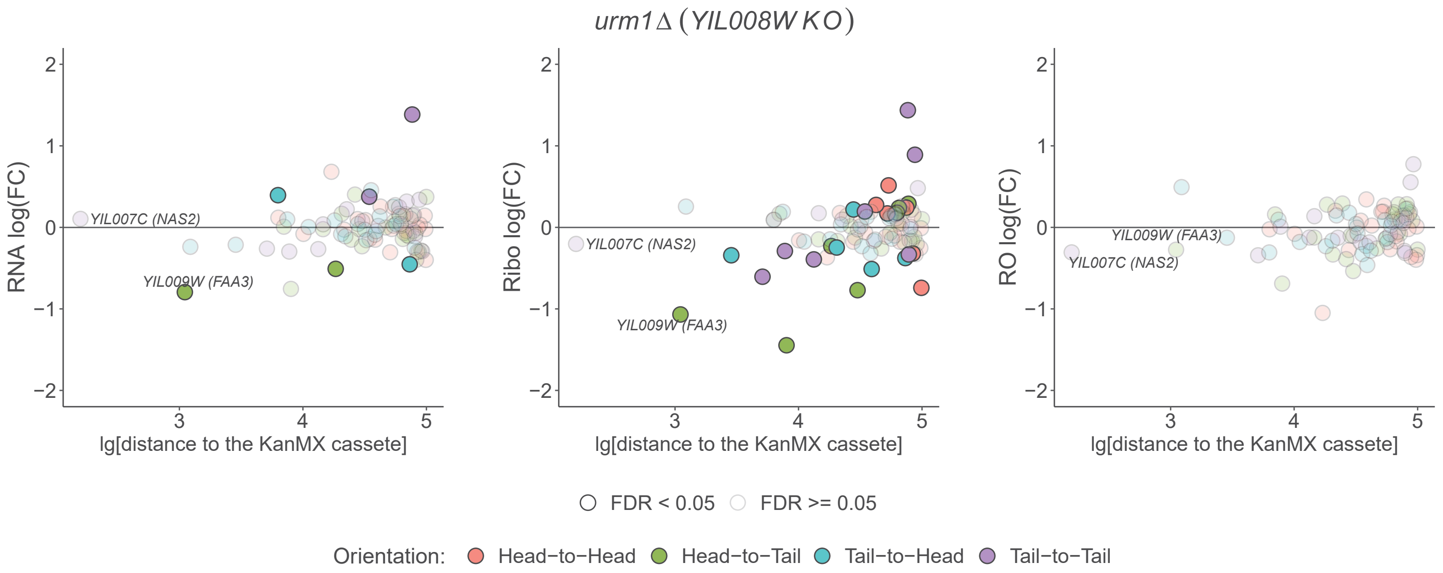
urm1Δ (YIL008W KO)
RNA log(FC)
Ribo log(FC)
RO log(FC)
YIL007C (NAS2)
YIL009W (FAA3)
lg[distance to the KanMX cassete]
FDR < 0.05
FDR >= 0.05
Orientation:
Head−to−Head
Head−to−Tail
Tail−to−Head
Tail−to−Tail

**B**

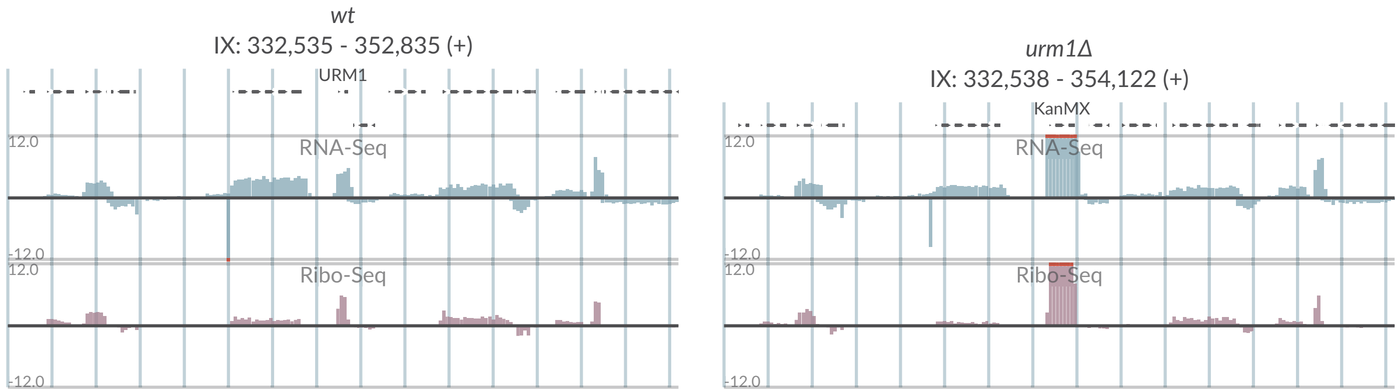
wt
IX: 332,535 - 352,835 (+)
URM1
RNA-Seq
Ribo-Seq
urm1Δ
IX: 332,538 - 354,122 (+)
KanMX
